## Supporting Information for "Eusociality through conflict dissolution"

*Proc. R. Soc. B*, DOI: 10.1098/rspb.2021.0386

Mauricio González-Forero[1*†] and Jorge Peña[2*†]

[1]School of Biology, University of St Andrews, St Andrews, UK

[2]Institute for Advanced Study in Toulouse, University of Toulouse Capitole, Toulouse, France

[*]Both authors contributed equally.

**8 Specification of Fig. 2, and additional figures** **73**

### Outline

This Supplementary Information contains the details of our evolutionary model and is organized as follows. First, in Evolutionary model set-up (section 1), we introduce assumptions, notions, and notation that will be used when building the model. Second, in Selection gradients (section 2), we build the population dynamics model that allows us to identify invasion fitness (i.e., the growth rate of a rare mutant subpopulation in a resident population at equilibrium); this enables us to calculate the selection gradients which provide the direction of selection. We obtain a generic expression of the selection gradient from a general formula of eigenvalue (here, invasion fitness) perturbation that writes the selection gradient in terms of reproductive value, stable mutant distribution, and the local sensitivity of mutant vital rates to marginal changes in trait values. Using the reproductive value and stable mutant distribution for our model, we obtain a generic yet simplified expression of the selection gradient for our model. We use this simplified expression to derive the selection gradient of the evolving traits we study (helping probability and reproductive effort). Third, in Inclusive fitness effects (section 3), we show that the selection gradients of all traits can be written in terms of inclusive fitness effects for all the model cases we consider. Fourth, in Conflict dissolution and benefit-cost ratio zones (section 4), we define conflict dissolution and show that a necessary condition for conflict dissolution via maternal reproductive specialization is that there is evolutionary synergy of reproductive effort on helping. Fifth, in Evolutionary synergy and trade-off alleviation (section 5), we show that such synergy is equivalent to trade-off alleviation by helpers if reproductive effort is optimal. Sixth, in Evolutionary dynamics (section 6), we postulate that the evolutionary dynamics satisfy a form of the "canonical equation" of adaptive dynamics. This enables us to use the derived selection gradients to write equations describing the evolutionary dynamics of the evolving traits. Seventh, in Specific functional forms (section 7), we specify functions for the vital rates and the joint helping probability which enables us to obtain numerical solutions for the evolutionary dynamics. Finally, in Specification of Fig. 2, and additional figures (section 8), we give the specification of functional forms and parameter values used to create the figures in the main text, and provide additional figures with results. Table S1 presents a summary of our notation.

Table S1: Summary of notation.

| Notation | Meaning |
|---|---|
| $p$ | Helping probability: probability that a first-brood offspring stays in the maternal nest and helps |
| $x$ | Maternal influence: maternal effort to induce first-brood offspring to become a helper |
| $y$ | Offspring resistance: offspring effort to resist the maternal influence |
| $z$ | Reproductive effort: maternal effort to produce second-brood offspring |
| $s_a$ | Offspring survival: probability that an offspring from brood $a \in \{1,2\}$ survives dispersal |
| $s_M$ | Parent survival: probability that a young couple becomes an old couple |
| $f_a$ | Fertility: number of offspring produced a couple of age $a \in \{1,2\}$ |
| $\sigma_{a,\ell}$ | Brood sex proportion: fraction of sex-$\ell$ offspring produced in brood $a \in \{1,2\}$ |
| $q_{\ell,i,k}$ | Transmission probability: probability that an offspring is of type $i \in \{\mathrm{r}, \mathrm{m}\}$ (resident or mutant) given it is of sex-$\ell$ and its parents are of type $k \in \{\mathrm{rm}, \mathrm{mr}\}$ (resident mother and mutant father or mutant mother and resident father) |
| $h$ | Expected number of helpers: expected number of helpers that a couple has |
| $F_{a,\ell,i,k}$ | Effective fertility: expected number of surviving reproductive, sex-$\ell$ offspring of type $i$ produced by an age-$a$ couple of type $k$ |
| $\Pi_{a,\ell,i,k}$ | Productivity: probability that a young couple survives to age $a$ times its effective fertility at that age |
| $N_{\ell,i}$ | Density of unmated individuals: number of unmated individuals of genotype $i$ and sex $\ell$ |
| $N_{a,k}$ | Density of couples: number of couples of age $a$ and type $k$ |
| $\mathcal{N}_k$ | Density of matings: number of matings of type $k$ before density dependence |
| $N$ | Fixed number of nesting sites in the population |
| $\alpha$ | Nest availability: density dependent probability that a new couple finds a nesting site |
| $\lambda$ | Invasion fitness: asymptotic growth rate of a rare mutant subpopulation in a resident population at demographic equilibrium |
| $\mathcal{S}_\zeta$ | Selection gradient of trait $\zeta$ |
| $\mathbf{u}$ | Stable mutant distribution: asymptotic distribution of neutral mutants |
| $\mathbf{v}$ | Reproductive values: long-term contribution by neutral mutants to the population |
| $\mathbf{G}$ | Genetic covariance matrix |
| $t$ | Ecological time |
| $\tau$ | Evolutionary time |
| $B$ | Marginal benefit of helping: marginal effect of helpers on late productivity |
| $C$ | Marginal cost of helping: marginal effect of helpers on early productivity |
| $D$ | Marginal productivity of late fertility: marginal effect of late fertility on late productivity |
| $\rho_{A,H,P}$ or $\rho_A$ | Relative reproductive worth for a random actor in $A$ relative to a random candidate helper in $H$ of a random payee in $P$ |

### 1 Evolutionary model set-up

#### 1.1 Basic assumptions and variables

**Adaptive dynamics assumptions.** We study the co-evolutionary dynamics of the helping probability $p$ of first-brood offspring and the reproductive effort $z$ devoted to the production of second-brood offspring by a mother. We do this by considering repeated invasion-fixation events of rare mutant alleles in a large population of resident alleles [1, 2, 3, 4]. We make the standard assumptions that each trait is controlled by a single locus, and that the effects of a mutation on trait values are marginally small and unbiased (i.e., a mutation is equally likely to increase or decrease the trait value). Given the small phenotypic effect of mutations and the large population size, a newly arisen mutation that is not neutral either becomes fixed or is eliminated. We also assume a standard separation of timescales. Specifically, we assume that mutation events are rare enough that natural selection either fixes or eliminates a non-neutral mutation before another mutation arises. The repetition of this mutant invasion sequence leads to evolutionary change in the resident phenotype. Thus, population dynamics occur in a fast "ecological" time scale $t$ (that we measure in discrete time) whereas evolutionary change occurs in a slow "evolutionary" timescale $\tau$ (that we measure in continuous time).

**Model cases.** We consider model cases that differ in three aspects. First, the genetic system (P, for "ploidy") can be either (i) diploid (P = D, in which case both sexes are diploid) or haplodiploid (P = HD, in which case females are diploid and males are haploid). Second, the individuals genetically controlling the helping behavior (C, for "control") can be either (i) offspring (C = O, for "offspring control"), (ii) the mother (C = M, for "maternal control"), or (iii) both mother and offspring (C = S, for "shared control"). Third, the sex of helpers (G, for "gender") can be either (i) female and male (G = B, for "both sexes help"), or (ii) exclusively female (G = F, for "only females help"). This yields twelve model cases (Fig. S1). For instance, in one model case the genetic system is diploid, helping is under offspring control, and both sexes help (D-O-B), which is relevant to termites if helping is under offspring control; in another model case, the genetic system is haplodiploid, helping is under shared control, and only females help (HD-S-F), which is relevant to eusocial hymenoptera if helping is under shared control. Although our focus is on model cases of shared control that allow us to study the evolutionary dynamics of parent-offspring conflict over helping, model cases of offspring control and maternal control serve as stepping stones in the building and analysis of model cases of shared control.

**Evolving traits.** For the model cases where helping is under either offspring or maternal control, we consider the coevolution of two traits: (i) the probability $p \in [0,1]$ that a first-brood offspring stays at the nest and becomes a helper, and (ii) the maternal reproductive effort $z \in \mathbb{R}_+^*$[1]. For all model cases, we assume that reproductive effort $z$ is exclusively under maternal control. Thus, when helping is under offspring or maternal control, we follow the evolution of the phenotypic vector $\mathbf{z} = (p, z)^\intercal$. For model cases where helping is under shared control, we consider the coevolution of three traits: maternal influence $x \in \mathbb{R}_+$, offspring resistance $y \in \mathbb{R}_+$, and maternal reproductive effort $z \in \mathbb{R}_+$. When considering helping under shared control, we assume

[1]Throughout, $\mathbb{R}_+$ refers to the set of non-negative reals, that is, $\mathbb{R}_+ = \{x \in \mathbb{R} | x \geq 0\}$. $\mathbb{R}_+^*$ refers to the set of positive reals, that is, $\mathbb{R}_+^* = \{x \in \mathbb{R} | x > 0\}$.

| | | | | | | | |
|---|---|---|---|---|---|---|---|
| | Sex of helpers (G) → | Both (B) | | | Female (F) | | |
| | Who controls help (C) → | Offspring (O) | Mother (M) | Shared (S) | Offspring (O) | Mother (M) | Shared (S) |
| Genetic system (P) | Diploid (D) | D-O-B | D-M-B | D-S-B | D-O-F | D-M-F | D-S-F |
| | Cases relevant to: | Termites<br>Snapping shrimp<br>Naked-mole rats | | | Social spiders<br>Ambrosia beetles | | |
| | Haplodiploid (HD) | HD-O-B | HD-M-B | HD-S-B | HD-O-F | HD-M-F | HD-S-F |
| | Cases relevant to: | Gall thrips | | | Eusocial hymenoptera | | |

Figure S1: Model cases we consider. Case relevance is based on Ross et al.[5] and Davies et al.[6].

that the helping probability $p(x, y)$ is a function of maternal influence $x$ and offspring resistance $y$ (i.e., $p(x, y)$ is a "joint phenotype" between mother and offspring; [7]). Thus, when helping is under shared control, we follow the evolution of the phenotypic vector $\mathbf{z} = (x, y, z)^{\intercal}$. For a given trait $\zeta$ (where $\zeta \in \{p, z\}$ for model cases of offspring and maternal control, and $\zeta \in \{x, y, z\}$ for model cases of shared control), we denote by $\zeta_{\mathrm{r}}$ the resident trait value and by $\zeta_{\mathrm{m}}$ the mutant trait value; similarly, we denote by $\mathbf{z}_{\mathrm{r}} = (\zeta_{\mathrm{r}})^{\intercal}$ the resident phenotypic vector and by $\mathbf{z}_{\mathrm{m}} = (\zeta_{\mathrm{m}})^{\intercal}$ the mutant phenotypic vector. By some abuse of notation, we also denote the resident trait value by $\zeta$ and the resident phenotypic vector by $\mathbf{z}$. It is then understood that $\zeta \equiv \zeta_{\mathrm{r}}$ and $\mathbf{z} \equiv \mathbf{z}_{\mathrm{r}}$.

**Life cycle.** We consider a finite but large population of individuals with a fixed number $N$ of nesting sites. Generations are overlapping, and the life cycle is lifetime monogamous with two offspring broods, as follows (Fig. S2). (i) In each nesting site, there is one singly mated female characterized by her genotype and the genotype of the male she mated or is mating with: we refer to a mated female and her mate as a "couple". We let $a$ index the age of a couple, so that $a = 1$ for a young couple and $a = 2$ for an old couple. We let $\ell$ denote the sex of an individual, so $\ell = ♀$ for a female and $\ell = ♂$ for a male. (ii) The female of a young couple produces and provides care for a fixed number $f_1$ of first-brood offspring, a proportion $\sigma_{1,\ell}$ of which are of sex $\ell$. A first-brood offspring of sex $\ell$ either remains at the nest with probability $p_\ell$ to become a non-reproductive helper, or disperses with probability $1 - p_\ell$. Each dispersed first-brood offspring survives dispersal with probability $s_1$ to become an unmated reproductive. Thus, a young couple produces $F_{1,\ell} = f_1\sigma_{1,\ell}(1 - p_\ell)s_1$ unmated reproductive offspring. (iii) A young couple either survives with probability $s_M$ to become an old couple or dies with probability $1 - s_M$. (iv) The female of an old couple produces a number $f_2$ of second-brood offspring, a proportion $\sigma_{2,\ell}$ of which are of sex $\ell$. A second-brood offspring always disperses, and survives dispersal with probability $s_2$ to become an unmated reproductive. Thus, an old couple produces $F_{2,\ell} = f_2\sigma_{2,\ell}s_2$ unmated reproductive offspring. We call $F_{a,\ell}$ the age-specific sex-specific effective fertility of a couple. Consequently, the expected number of sex-$\ell$ unmated reproductives produced by a couple through first-brood offspring is $\Pi_{1,\ell} = F_{1,\ell}$, and the expected number of sex-$\ell$ unmated reproductives produced by a couple through second-brood offspring is $\Pi_{2,\ell} = s_M F_{2,\ell}$. We call $\Pi_{a,\ell}$ the age-specific sex-specific productivity of a couple. (v) Old couples die. (vi) Unmated reproductives mate singly at random and establish nests subject to the availability of nesting sites, which is measured by $\alpha$. Mated reproductives that fail to establish a nest die.

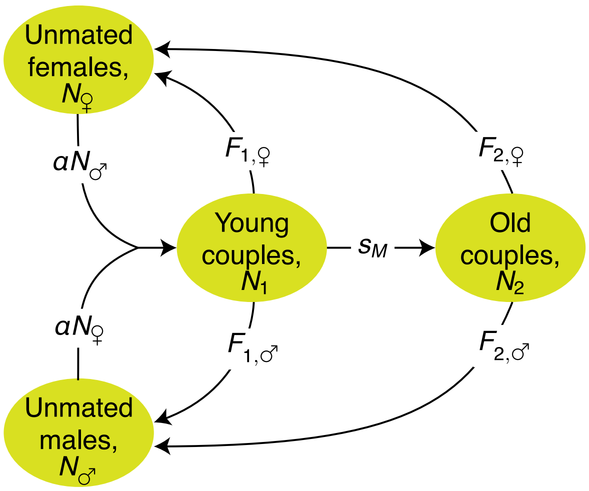

Figure S2: Resident life cycle. Unmated females and males mate once to become young couples that may survive to become old couples. Each couple occupies a single nesting site, the number of which is constant. The female of a young couple produces first-brood offspring and when the couple is old the female produces second-brood offspring. Each ellipse corresponds to a "demographic class" of individuals or of couples of individuals. Here $N_j$ is the number of individuals of demographic class $j$, $F_{a,\ell}$ is the effective fertility of a couple of age $a$ through sex-$\ell$ offspring, and $\alpha$ measures the density dependent probability that a new couple finds a nesting site.

**Genotypes.** Consideration of mutant genotypes leads to a complete life cycle comprising ten classes of individuals or of couples of individuals (Fig. S3). We let $i$ index the genotype of unmated individuals. The genotype $i$ of an unmated individual can be either r for a resident or m for a mutant, where due to the assumption that the mutant allele is rare, a mutant is heterozygous in diploids and in female haplodiploids, and hemizygous in male haplodiploids. Similarly, we let $k$ index the "type" of a couple, which comprises the genotype of the female and the genotype of the male of the couple in that order. That is, the type $k$ of a couple can be (i) rr when the female and male are both residents, (ii) rm when the female is resident and the male is mutant, or (iii) mr when the female is mutant and the male is resident. We do not need to consider the couple type mm comprising a mutant female and a mutant male, as the frequency of such type is negligible when the mutant allele is rare. For a couple of type $k$, we denote by ♀$(k)$ the genotype of the female and by ♂$(k)$ the genotype of the male in the couple, that is,

$$♀(k) = \begin{cases} \text{m} & \text{if } k = \text{mr} \\ \text{r} & \text{if } k = \text{rr or } k = \text{rm}, \end{cases} \tag{S1.1.1a}$$

and

$$♂(k) = \begin{cases} \text{m} & \text{if } k = \text{rm} \\ \text{r} & \text{if } k = \text{rr or } k = \text{mr}. \end{cases} \tag{S1.1.1b}$$

**Dependence of vital rates on the evolving traits.** We assume that early fertility $f_1$ and first-brood survival $s_1$ are constants. In contrast, we assume that the couple's survival $s_M$, the late fertility $f_2$, and the second-brood survival $s_2$ depend on the individuals' genotypes. Thus, the vital rates $s_M$, $f_2$, and $s_2$ are functions of

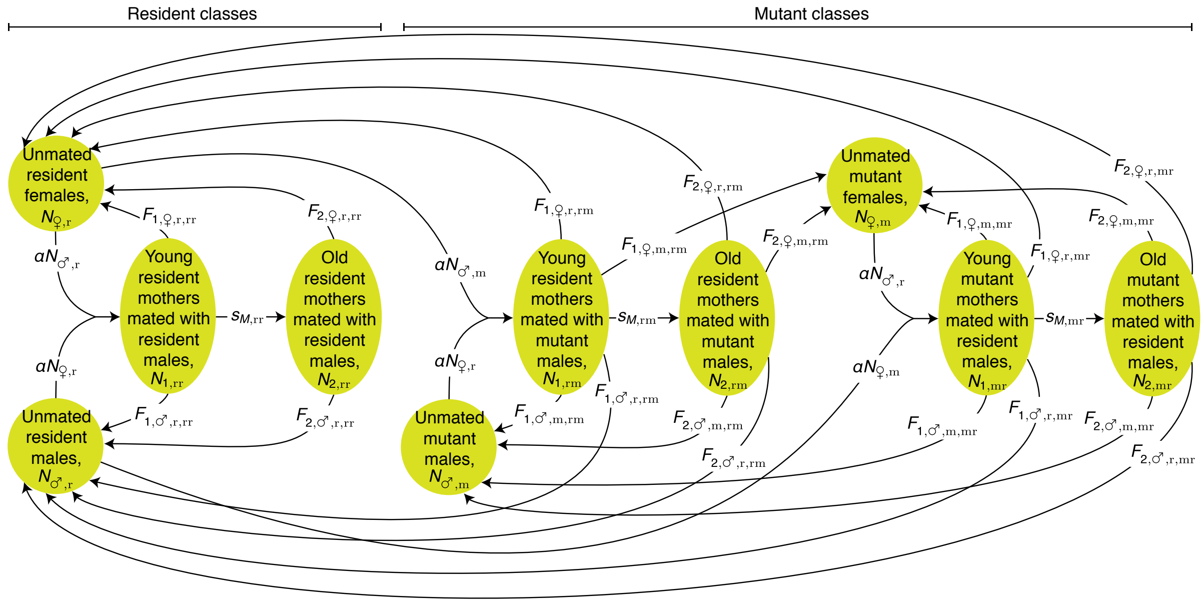

Figure S3: Resident-mutant life cycle. There are ten demographic classes, of which four exclusively involve resident genotypes and six involve mutant genotypes.

the evolving phenotype $\mathbf{z}$. More specifically, we assume that the vital rates $s_M$, $f_2$, and $s_2$ are functions of the expected number of helpers $h_k$ and the reproductive effort $z_k$ that an old couple of type $k$ has. We express $h_k$ in terms of genotypes in section 1.3 below. Regarding $z_k$, since reproductive effort is always under maternal control, the reproductive effort of an old couple of type $k$ is

$$z_k = z_{♀(k)}, \tag{S1.1.2a}$$

which, via equation (S1.1.1a), equals $z_\mathrm{r}$ ($\equiv z$) if the female in the couple is resident or $z_\mathrm{m}$ if she is mutant. With our notational conventions, this implies that

$$z_\mathrm{rr} = z_\mathrm{r} \equiv z \tag{S1.1.2b}$$

always holds.

**Brood sex proportions.** As previously stated, we denote by $\sigma_{a,\ell}$ the proportion of offspring of sex $\ell$ produced by a couple of age $a$. The brood sex proportions satisfy

$$\sum_{\ell\in\{♀,♂\}} \sigma_{a,\ell} = 1 \; \forall a \in \{1,2\} \tag{S1.1.3}$$

because each offspring is either a female or a male. In the following, we will also use the shorthand notation $\sigma_a \equiv \sigma_{a,♀}$, and refer to it as the sex proportion of brood $a$. Additionally, we will also write

$$\boldsymbol{\sigma}_a^\intercal = \left(\sigma_{a,♀}, \sigma_{a,♂}\right) \tag{S1.1.4}$$

for the vector collecting the sex proportions of brood $a$.

**Maximum number of helpers.** We denote the maximum number helpers by $\bar{h}$. For model cases where both sexes help (G = B),

$$\bar{h} = f_1. \tag{S1.1.5a}$$

For model cases where only females help (G = F),

$$\bar{h} = f_1 \sigma_1. \tag{S1.1.5b}$$

#### 1.2 Transmission and helping probabilities

**Transmission probability.** We denote by $q_{\ell,i,k}$ the probability that an offspring is of genotype $i$ given that it is of sex $\ell$ and that its parents are of type $k$. We refer to this conditional probability as the transmission probability, and list its values in Fig. S4. Although the transmission probability depends on the genetic system (diploid or haplodiploid), it invariably satisfies the following set of identities:

$$q_{\ell,\mathrm{r},\mathrm{rr}} = 1 \quad \forall \ell \in \{♀, ♂\}, \tag{S1.2.1a}$$

$$q_{\ell,\mathrm{m},\mathrm{rr}} = 0 \quad \forall \ell \in \{♀, ♂\}, \tag{S1.2.1b}$$

$$\sum_{i \in \{\mathrm{r},\mathrm{m}\}} q_{\ell,i,k} = 1 \quad \forall \ell \in \{♀, ♂\} \text{ and } \forall k \in \{\mathrm{rr}, \mathrm{rm}, \mathrm{mr}\}, \tag{S1.2.1c}$$

$$\sum_{\ell \in \left\{♀, ♂\right\}} q_{\ell,i,k} = 1 \quad \forall i \in \{\mathrm{r}, \mathrm{m}\} \text{ and } \forall k \in \{\mathrm{rm}, \mathrm{mr}\}. \tag{S1.2.1d}$$

Equations (S1.2.1a) and (S1.2.1b) state that all offspring of a resident couple (rr) are resident (r) regardless of their sex. Equation (S1.2.1c) holds because an offspring is either resident or mutant, regardless of its sex and the genotypes of its parents. Finally, (S1.2.1d) states that when parents have different genotypes (one being resident, the other mutant), and for each possible genotype of the offspring, the transmission probability is a probability distribution over the sexes of the offspring.

The ratio

$$\frac{q_{♀,\mathrm{m},\mathrm{rm}}}{q_{♂,\mathrm{m},\mathrm{mr}}} \tag{S1.2.2a}$$

will naturally arise in our analysis. This ratio can be interpreted as a measure of transmission asymmetry across sexes inherent to the genetic system, that is, a measure of how likely a mutant father is to transmit his mutant allele to a daughter (the numerator of (S1.2.2a), $q_{♀,\mathrm{m},\mathrm{rm}}$) compared to how likely a mutant mother is to transmit her allele to a son (the denominator of (S1.2.2a), $q_{♂,\mathrm{m},\mathrm{mr}}$). It can be checked that the ratio (S1.2.2a) simplifies to

$$\frac{q_{♀,\mathrm{m},\mathrm{rm}}}{q_{♂,\mathrm{m},\mathrm{mr}}} = \begin{cases} 1 & \text{for diploids (G = D)} \\ 2 & \text{for haplodiploids (G = HD)} \end{cases}. \tag{S1.2.2b}$$

Equation (S1.2.2b) states that there is no transmission asymmetry across sexes in diploids, but that in haplodiploids mutant fathers are twice as likely to transmit their mutant alleles to their daughters as mutant mothers are to transmit their mutant alleles to their sons. We will see that such transmission asymmetry means that, for diploids, a neutral mutation is asymptotically equally likely to occur in the female or the male of a couple; in contrast, for haplodiploids, a neutral mutation is asymptotically twice as likely to occur in the female rather than the male of a couple.

| Genetic system ↓ | Offspring sex, $\ell$ → / Parent type, $k$ → / Offspring genotype, $i$ ↓ | ♀ rr | ♀ rm | ♀ mr | ♂ rr | ♂ rm | ♂ mr |
|---|---|---|---|---|---|---|---|
| Diploid | r | 1 | 1/2 | 1/2 | 1 | 1/2 | 1/2 |
| | m | 0 | 1/2 | 1/2 | 0 | 1/2 | 1/2 |
| Haplodiploid | r | 1 | 0 | 1/2 | 1 | 1 | 1/2 |
| | m | 0 | 1 | 1/2 | 0 | 0 | 1/2 |

The elements of
(Eq. S1.2.1c)
(Eq. S1.2.1d)
(Eq. S1.2.1d)
sum to 1

Figure S4: Transmission probability. List of values for the conditional probability $q_{i,\ell,k}$ that an offspring is of genotype $i$ given that it is of sex $\ell$ and that its parents are of type $k$. Identities (S1.2.1c) and (S1.2.1d) are highlighted in color.

**Helping probability.** We denote by $p_{\ell,i,k}$ the probability that an offspring of sex $\ell$ and genotype $i$ produced by a couple of type $k$ does not disperse and instead stays at the nest to become a helper. We refer to this conditional probability as the helping probability and list its values in Fig. S5. The helping probability depends on (i) whether both sexes or only females help and (ii) whether helping is under offspring, maternal, or shared control. For model cases of shared control, we define the helping probability function

$$p : \mathbb{R}_+ \times \mathbb{R}_+ \to [0,1]$$
$$(x, y) \mapsto p(x, y),$$

such that $p(x, y)$ is the helping probability of an offspring when the mother exerts influence $x$ and the offspring exerts resistance $y$. We assume that $p$ is smooth, increasing in $x$, and decreasing in $y$, so that

$$\frac{\partial p}{\partial x} > 0, \quad \text{(S1.2.3a)}$$
$$\frac{\partial p}{\partial y} < 0 \quad \text{(S1.2.3b)}$$

hold for all the domain of $p(x, y)$. That is, an increase in maternal influence (resp. an increase in offspring resistance) increases (resp. decreases) the probability that a first-brood offspring becomes a helper.

#### 1.3 Expected number of helpers

**Expected number of helpers of a couple of type $k$.** As previously stated, the evolving phenotype $\mathbf{z}$ modulates the vital rates $s_M$, $f_2$, and $s_2$ because these vital rates are functions of the expected number of helpers $h_k$ and of the reproductive effort $z_k$ that each old couple of type $k$ has. We now derive an expression for $h_k$ in terms of individuals' genotypes. We start by using the definitions of the transmission probability $q_{\ell,i,k}$ and the helping probability $p_{\ell,i,k}$ (section 1.2) to write an expression for the expected proportion of helpers of sex $\ell$ among the first-brood offspring of a couple of type $k$, $p_{\ell,k}$, as

$$p_{\ell,k} = \sum_{i \in \{\mathrm{r},\mathrm{m}\}} q_{\ell,i,k} p_{\ell,i,k}. \quad \text{(S1.3.1)}$$

| | | Offspring sex, $\ell$ → | ♀ | | | ♂ | | |
|---|---|---|---|---|---|---|---|---|
| Sex of helpers ↓ | Who controls help ↓ | Parent type, $k$ → Offspring genotype, $i$ ↓ | rr | rm | mr | rr | rm | mr |
| Both, B | Offspring, O | r | $p$ | $p$ | $p$ | $p$ | $p$ | $p$ |
| | | m | $p_{\mathrm{m}}$ | $p_{\mathrm{m}}$ | $p_{\mathrm{m}}$ | $p_{\mathrm{m}}$ | $p_{\mathrm{m}}$ | $p_{\mathrm{m}}$ |
| | Mother, M | r | $p$ | $p$ | $p_{\mathrm{m}}$ | $p$ | $p$ | $p_{\mathrm{m}}$ |
| | | m | $p$ | $p$ | $p_{\mathrm{m}}$ | $p$ | $p$ | $p_{\mathrm{m}}$ |
| | Shared, S | r | $p(x,y)$ | $p(x,y)$ | $p(x_{\mathrm{m}},y)$ | $p(x,y)$ | $p(x,y)$ | $p(x_{\mathrm{m}},y)$ |
| | | m | $p(x,y_{\mathrm{m}})$ | $p(x,y_{\mathrm{m}})$ | $p(x_{\mathrm{m}},y_{\mathrm{m}})$ | $p(x,y_{\mathrm{m}})$ | $p(x,y_{\mathrm{m}})$ | $p(x_{\mathrm{m}},y_{\mathrm{m}})$ |
| Female, F | Offspring, O | r | $p$ | $p$ | $p$ | 0 | 0 | 0 |
| | | m | $p_{\mathrm{m}}$ | $p_{\mathrm{m}}$ | $p_{\mathrm{m}}$ | 0 | 0 | 0 |
| | Mother, M | r | $p$ | $p$ | $p_{\mathrm{m}}$ | 0 | 0 | 0 |
| | | m | $p$ | $p$ | $p_{\mathrm{m}}$ | 0 | 0 | 0 |
| | Shared, S | r | $p(x,y)$ | $p(x,y)$ | $p(x_{\mathrm{m}},y)$ | 0 | 0 | 0 |
| | | m | $p(x,y_{\mathrm{m}})$ | $p(x,y_{\mathrm{m}})$ | $p(x_{\mathrm{m}},y_{\mathrm{m}})$ | 0 | 0 | 0 |

Figure S5: Helping probability. List of values for the helping probability $p_{\ell,i,k}$ for the model cases considered.

The proportion of helpers of either sex among the first-brood offspring of a couple of type $k$ can then be written as

$$p_k = \sum_{\ell \in \{♀,♂\}} \sigma_{1,\ell} p_{\ell,k} = \sum_{\ell \in \{♀,♂\}} \sigma_{1,\ell} \sum_{i \in \{\mathrm{r},\mathrm{m}\}} q_{\ell,i,k} p_{\ell,i,k}, \tag{S1.3.2}$$

from which the expected number of helpers $h_k$ is derived as

$$h_k = f_1 p_k \tag{S1.3.3a}$$

$$= f_1 \sum_{\ell \in \{♀,♂\}} \sigma_{1,\ell} \sum_{i \in \{\mathrm{r},\mathrm{m}\}} q_{\ell,i,k} p_{\ell,i,k}. \tag{S1.3.3b}$$

**Expected number of helpers of a resident couple.** The expected number of helpers of a couple of type rr (i.e., the expected number of helpers per nest in a resident population) will be important in our analysis. We adopt a notational convention similar to the one we have adopted for the helping probability $p$, namely to use $h$ as (i) a generic variable referring to the expected number of helpers, (ii) as the value of such variable for the specific case of a couple of type rr (i.e., $h \equiv h_{\mathrm{rr}}$), and (iii) as a function of evolving traits whose output is the expected number of helpers, to be specified below. With these conventions, the expected number of helpers available to a couple of type rr can be written as

$$h_{\mathrm{rr}} \equiv h \tag{S1.3.4a}$$

$$= f_1 \sum_{\ell \in \{♀,♂\}} \sigma_{1,\ell} \sum_{i \in \{\mathrm{r},\mathrm{m}\}} q_{\ell,i,\mathrm{rr}} p_{\ell,i,\mathrm{rr}}$$

$$= f_1 \sum_{\ell \in \{♀,♂\}} \sigma_{1,\ell} p_{\ell,\mathrm{r},\mathrm{rr}}, \tag{S1.3.4b}$$

where the first equality follows from expression (S1.3.3b) with $k = \text{rr}$, and the last one from identities (S1.2.1a) and (S1.2.1b). By inspection of the values of the helping probability given in Fig. S5, and since $\sigma_{1,♀} + \sigma_{1,♂} = 1$ (S1.1.3) holds, expression (S1.3.4b) reduces to

$$h_{\text{rr}} \equiv h = h(p) = \bar{h}p \tag{S1.3.5a}$$

for model cases of offspring or maternal control, and to

$$h_{\text{rr}} \equiv h = h(x, y) = \bar{h}p(x, y) \tag{S1.3.5b}$$

for model cases of shared control. Here, $\bar{h} = f_1$ for model cases where both sexes help (S1.1.5a) and $\bar{h} = f_1\sigma_1$ for model cases where only females help (S1.1.5b). In expression (S1.3.5a) we have used the expected number of helpers function

$$\begin{aligned} h : [0,1] &\to [0, \bar{h}] \\ p &\mapsto \bar{h}p, \end{aligned}$$

such that $h(p) = \bar{h}p$, while $h(x, y)$ in expression (S1.3.5b) refers to the function

$$\begin{aligned} h : \mathbb{R}_+ \times \mathbb{R}_+ &\to [0, \bar{h}] \\ (x, y) &\mapsto \bar{h}p(x, y), \end{aligned}$$

such that $h(x, y) = \bar{h}p(x, y)$.

#### 1.4 Assumptions on vital rates

The mechanism of conflict dissolution that we identify rests on three critical assumptions. First, we assume that the late fertility of a mother can evolve (genetically or plastically). Second, we assume that mothers face life-history trade-offs (i) between fertility and survival to old age; (ii) between fertility and survival of second-brood offspring; or (iii) between fertility and both survival rates. Finally, we assume that such life-history trade-offs can be alleviated by helpers. We now formalize each of these assumptions.

**Late fertility of a couple of type $k$, $f_{2,k}$.** We assume that the number of second-brood offspring produced by a couple of type $k$, $f_{2,k}$, depends on the mother's reproductive effort, $z_k = z_{♀(k)}$ (S1.1.2a), via

$$f_{2,k} = f_2(z_k), \tag{S1.4.1}$$

where

$$\begin{aligned} f_2 : \mathbb{R}_+^* &\to \mathbb{R}_+^* \\ z &\mapsto f_2(z), \end{aligned} \tag{S1.4.2}$$

is a smooth function. Furthermore, we assume $f_2$ is strictly increasing; that is,

$$\frac{\mathrm{d}f_2}{\mathrm{d}z} > 0 \tag{S1.4.3}$$

holds for all $z \in \mathbb{R}_+^*$. Equations (S1.4.1) and (S1.4.3) respectively encapsulate the assumptions that mother's late fertility depends on the evolving mother's reproductive effort $z_k$, and that a larger reproductive effort implies a larger late fertility $f_{2,k}$.

**Survival probabilities $s_{M,k}$ and $s_{2,k}$.** We assume that the survival probabilities $s_{M,k}$ and $s_{2,k}$ can be written as functions of both the late fertility, $f_{2,k}$, and the expected number of helpers, $h_k$, of a couple of type $k$. More explicitly, we let the survival probabilities be given by

$$s_{M,k} = s_M(f_{2,k}, h_k) = s_M(f_2(z_k), h_k), \tag{S1.4.4a}$$

$$s_{2,k} = s_2(f_{2,k}, h_k) = s_2(f_2(z_k), h_k), \tag{S1.4.4b}$$

where the rightmost equalities follow from (S1.4.1), and where

$$\begin{aligned} s_M : S_M \times [0, \bar{h}] &\to (0,1) \\ (f_2, h) &\mapsto s_M(f_2, h), \end{aligned} \tag{S1.4.5a}$$

$$\begin{aligned} s_2 : S_2 \times [0, \bar{h}] &\to (0,1] \\ (f_2, h) &\mapsto s_2(f_2, h), \end{aligned} \tag{S1.4.5b}$$

are smooth functions decreasing in $f_2$[2]. In (S1.4.5), $S_M$ and $S_2$ are subsets of $\mathbb{R}_+^*$.

We assume that either $s_M$ or $s_2$ is decreasing in $f_2$, that is,

$$\frac{\partial s_M}{\partial f_2} < 0 \text{ or} \tag{S1.4.6a}$$

$$\frac{\partial s_2}{\partial f_2} < 0 \tag{S1.4.6b}$$

holds for all $f_2$ and all $h$ in the domains of these functions and where neither of the two derivatives is positive. Inequalities (S1.4.6) encapsulate the idea that mothers face a life-history trade-off between fertility and survival: all else being equal, a greater investment in late fertility $f_2$ from the part of the mother negatively affects at least one vital rate among $s_M$ and $s_2$.

Finally, we assume that either $s_M$ or $s_2$ is increasing in $h$, that is,

$$\frac{\partial s_M}{\partial h} > 0 \text{ or} \tag{S1.4.7a}$$

$$\frac{\partial s_2}{\partial h} > 0, \tag{S1.4.7b}$$

holds for all $f_2$ and all $h$ in the domains of these functions and where neither of the two derivatives is negative. Inequalities (S1.4.7) encapsulate the idea that helpers can increase the vital rates negatively affected by an increase in the mother's reproductive effort, thus potentially alleviating the trade-offs involved.

#### 1.5 Effective fertility

The early effective fertility $F_{1,\ell,i,k}$ gives the expected number of offspring of sex $\ell$ and genotype $i$ that successfully disperse and that are produced by a couple of age 1 and type $k$. The early effective fertility is given by

$$F_{1,\ell,i,k} = f_1 \sigma_{1,\ell} q_{\ell,i,k} (1 - p_{\ell,i,k}) s_1. \tag{S1.5.1}$$

[2]The upper bound of the codomain of $s_M$ is open so that $s_{M,\mathrm{rr}} < 1$ and the resident equilibrium of the resident system is stable, as we will show below. The lower bounds of the codomains of $s_M$ and $s_2$ are open so that, respectively, there are old couples and second-brood offspring can become reproductive.

Indeed, a young couple produces a fixed number $f_1$ of first-brood offspring, a proportion $\sigma_{1,\ell}$ of which are of sex $\ell$. Of these, a proportion $q_{\ell,i,k}$ is of genotype $i$, of which a proportion $(1-p_{\ell,i,k})s_1$ both disperses and survives dispersal. In particular, letting $i=\text{r}$, $k=\text{rr}$, and using identity (S1.2.1a), we find

$$F_{1,\ell,\text{r},\text{rr}} = f_1\sigma_{1,\ell}(1-p_{\ell,\text{r},\text{rr}})s_1 \tag{S1.5.2}$$

as an expression for the early effective fertility $F_{1,\ell,\text{r},\text{rr}}$ of a resident couple of type rr through offspring of genotype r and sex $\ell$ (i.e., the early rate of production of offspring of sex $\ell$ by a resident couple in a resident population).

An old couple of type $k$ produces a number of offspring $f_{2,k}$, a proportion $\sigma_{2,\ell}$ of which are of sex $\ell$. With probability $q_{\ell,i,k}$ one of such offspring of sex $\ell$ is of genotype $i$, with probability one it disperses (as we assume that all second-brood offspring disperse from their parental nest), and with probability $s_{2,k}$ it survives dispersal. It follows that the late effective fertility $F_{2,\ell,i,k}$ (giving the expected number of individuals of sex $\ell$ and genotype $i$ that successfully disperse and that are produced by a couple of age 2 and type $k$) is given by

$$F_{2,\ell,i,k} = f_{2,k}\sigma_{2,\ell}q_{\ell,i,k}s_{2,k}. \tag{S1.5.3}$$

Similarly to early effective fertility, the late effective fertility of a resident couple in a resident population evaluates to

$$F_{2,\ell,\text{r},\text{rr}} = f_{2,\text{rr}}\sigma_{2,\ell}s_{2,\text{rr}}. \tag{S1.5.4}$$

#### 1.6 Productivity

We will show that the selection gradient in our model can be conveniently written in terms of what we term the age-specific and sex-specific productivity of a couple. The productivity $\Pi_{\ell,i,k}$ of a $k$-type couple through offspring of sex $\ell$ and genotype $i$ is the expected lifetime number of unmated reproductive offspring of sex $\ell$ and genotype $i$ produced by a couple of type $k$. The productivity of a $k$-type couple through offspring of sex $\ell$ and genotype $i$ is given by the sum of a young couple's effective fertility and the old couple's effective fertility, the latter discounted by the probability $s_{M,k}$ that a young couple survives to old age. From this, we have

$$\Pi_{\ell,i,k} = F_{1,\ell,i,k} + s_{M,k}F_{2,\ell,i,k}. \tag{S1.6.1}$$

It will prove useful for our subsequent analysis to highlight the two summands of the previous expression with more dedicated notation. We will then alternatively write the productivity of a $k$-type couple through offspring of sex $\ell$ and genotype $i$ as

$$\Pi_{\ell,i,k} = \Pi_{1,\ell,i,k} + \Pi_{2,\ell,i,k}, \tag{S1.6.2}$$

where the first and second summands are respectively the early and late productivity of a couple of type $k$ through offspring of sex $\ell$ and genotype $i$. These are given by

$$\Pi_{1,\ell,i,k} = F_{1,\ell,i,k} = q_{\ell,i,k}\sigma_{1,\ell}f_1(1-p_{\ell,i,k})s_1, \tag{S1.6.3a}$$

$$\Pi_{2,\ell,i,k} = s_{M,k}F_{2,\ell,i,k} = q_{\ell,i,k}\sigma_{2,\ell}s_{M,k}f_{2,k}s_{2,k}, \tag{S1.6.3b}$$

where the second equalities follow from substituting the expressions for early and late effective fertility (equations (S1.5.1) and (S1.5.3)) into (S1.6.1) and rearranging.

We define the (total) early and late productivity of a type-$k$ couple as the sum of the productivities of each age over both sexes (female and male) and both genotypes (resident and mutant) of offspring. We can use previously established relationships between our variables to write down relatively simple expressions for these two quantities. The early productivity of a type-$k$ couple can be then written as:

$$\begin{aligned}
\Pi_{1,k} &= \sum_{\ell\in\{♀,♂\}} \sum_{i\in\{\mathrm{r,m}\}} \Pi_{1,\ell,i,k} && \text{(S1.6.4)}\\
&= \sum_{\ell\in\{♀,♂\}} \sum_{i\in\{\mathrm{r,m}\}} q_{\ell,i,k}\sigma_{1,\ell} f_1 (1-p_{\ell,i,k}) s_1 && \text{(S1.6.5)}\\
&= f_1 s_1 \sum_{\ell\in\{♀,♂\}} \sigma_{1,\ell} \sum_{i\in\{\mathrm{r,m}\}} q_{\ell,i,k}(1-p_{\ell,i,k})\\
&= f_1 s_1 \sum_{\ell\in\{♀,♂\}} \sigma_{1,\ell} \left( \sum_{i\in\{\mathrm{r,m}\}} q_{\ell,i,k} - \sum_{i\in\{\mathrm{r,m}\}} q_{\ell,i,k} p_{\ell,i,k} \right)\\
&= f_1 s_1 \sum_{\ell\in\{♀,♂\}} \sigma_{1,\ell} \left(1-p_{\ell,k}\right) && \text{(S1.6.6)}\\
&= f_1 s_1 \left( \sum_{\ell\in\{♀,♂\}} \sigma_{1,\ell} - \sum_{\ell\in\{♀,♂\}} \sigma_{1,\ell} p_{\ell,k} \right)\\
&= f_1 s_1 (1-p_k) && \text{(S1.6.7)}\\
&= \left(f_1 - h_k\right) s_1, && \text{(S1.6.8)}
\end{aligned}$$

where line (S1.6.5) follows from substituting (S1.6.3a) into (S1.6.4); line (S1.6.6) follows from identities (S1.2.1c) and (S1.3.1); line (S1.6.7) follows from identities (S1.1.3) and (S1.3.2); and line (S1.6.8) uses (S1.3.3a) and rearranges. Expression (S1.6.8) makes it explicit that the early productivity of a $k$-type couple is equal to the expected number of first-brood offspring that do not become helpers and instead disperse $(f_1 - h_k)$ times the probability that they survive dispersal $(s_1)$. To capture this in a general way, we define the *early productivity function*

$$\begin{aligned}
\Pi_1 : [0, f_1] &\to \mathbb{R}_+ \\
h &\mapsto \left(f_1 - h\right) s_1,
\end{aligned} \qquad \text{(S1.6.9)}$$

such that $\Pi_1(h) = \left(f_1 - h\right) s_1$.

Similarly, the late productivity of a type-$k$ couple can be written as:

$$\begin{aligned}
\Pi_{2,k} &= \sum_{\ell\in\{♀,♂\}} \sum_{i\in\{\mathrm{r,m}\}} \Pi_{2,\ell,i,k} && \text{(S1.6.10)}\\
&= \sum_{\ell\in\{♀,♂\}} \sum_{i\in\{\mathrm{r,m}\}} q_{\ell,i,k}\sigma_{2,\ell} s_{M,k} f_{2,k} s_{2,k} && \text{(S1.6.11)}\\
&= s_{M,k} f_{2,k} s_{2,k} \sum_{\ell\in\{♀,♂\}} \sigma_{2,\ell} \sum_{i\in\{\mathrm{r,m}\}} q_{\ell,i,k}\\
&= s_{M,k} f_{2,k} s_{2,k} \sum_{\ell\in\{♀,♂\}} \sigma_{2,\ell} && \text{(S1.6.12)}\\
&= s_{M,k} f_{2,k} s_{2,k}, && \text{(S1.6.13)}
\end{aligned}$$

where line (S1.6.11) follows from substituting (S1.6.3b) into (S1.6.10); line (S1.6.12) follows from identity (S1.2.1c); and line (S1.6.13) follows from identity (S1.1.3).

The (total) productivity of a couple of type $k$ is the sum of its early and late productivities, that is

$$\Pi_k = \Pi_{1,k} + \Pi_{2,k}. \tag{S1.6.14}$$

Two further identities concerning productivities are worth pointing out. First, note that, by substituting (S1.4.1) and (S1.4.4) into (S1.6.13), the late productivity of a couple of type $k$ is given by

$$\Pi_{2,k} = s_M(f_2(z_k), h_k) f_2(z_k) s_2(f_2(z_k), h_k).$$

This motivates our introduction of the *late productivity function*

$$\begin{aligned} \Pi_2 : \mathbb{R}_+^* \times [0, f_1] &\to \mathbb{R}_+^* \\ (f_2, h) &\mapsto s_M(f_2, h) f_2 s_2(f_2, h), \end{aligned} \tag{S1.6.15}$$

such that $\Pi_2(f_2, h) = s_M(f_2, h) f_2 s_2(f_2, h)$. The late productivity of a couple of type $k$ can then be written as

$$\Pi_{2,k} = \Pi_2(f_{2,k}, h_k). \tag{S1.6.16}$$

Second, substituting equation (S1.6.1) into (S1.6.2) and by identity (S1.6.13) we find that the productivity of a $k$-type mother through offspring of sex $\ell$ and genotype $i$ (S1.6.2) can be also written as

$$\Pi_{\ell,i,k} = q_{\ell,i,k} \left[ \sigma_{1,\ell} f_1 (1 - p_{\ell,i,k}) s_1 + \sigma_{2,\ell} \Pi_{2,k} \right]. \tag{S1.6.17}$$

In particular, and by setting $i = \mathrm{r}$ and $k = \mathrm{rr}$ in the previous expression, the productivity of a rr-type mother through offspring of sex $\ell$ and type r (i.e., the productivity of a mother through offspring of sex $\ell$ in a resident population) is given by

$$\begin{aligned} \Pi_{\ell,\mathrm{r},\mathrm{rr}} &= q_{\ell,\mathrm{r},\mathrm{rr}} \left[ \sigma_{1,\ell} f_1 (1 - p_{\ell,\mathrm{r},\mathrm{rr}}) s_1 + \sigma_{2,\ell} \Pi_{2,\mathrm{rr}} \right] \\ &= \sigma_{1,\ell} f_1 (1 - p_{\ell,\mathrm{r},\mathrm{rr}}) s_1 + \sigma_{2,\ell} \Pi_{2,\mathrm{rr}}, \end{aligned} \tag{S1.6.18}$$

where the second equality follows from identity (S1.2.1a).

### 2 Selection gradients

We now derive the selection gradients for our model. To do this, we proceed in nine steps. First, we build a population dynamics model of a resident population and a rare mutant subpopulation (Resident-mutant population dynamics; section 2.1). Second, we find the unique stable resident equilibrium where the mutant is absent (Resident population dynamics and resident equilibrium; section 2.2). Third, we identify invasion fitness, which is the growth rate of a rare mutant population around such resident equilibrium (Invasion fitness; section 2.3). Fourth, we write a general expression for the selection gradient, which gives the direction of selection in phenotypic space, by applying a general result on the sensitivity of the leading eigenvalue of irreducible and nonnegative matrices [8, 9, 1, 10]. This expression gives the selection gradient in terms of marginal effects of the mutant on vital rates weighted by reproductive values and the components of the stable mutant distribution (Selection gradient (generic form); section 2.4). Fifth, we calculate the neutral mutant submatrix required to obtain such reproductive values and stable mutant distribution (Neutral mutant submatrix, $\mathbf{J}^{\circ}_{\mathrm{mut}}$; section 2.5). Sixth, we find the reproductive values and stable mutant distribution for our model (Reproductive values and stable distribution; section 2.6). Seventh, using the particular form of the reproductive values and the stable mutant distribution for our model, we obtain a simplified expression of the selection gradient in terms of a couple's productivity weighted by reproductive values and stable mutant proportions of different classes (Selection gradient (generic, simplified form); section 2.7). Eighth, using such simplified selection gradient, we obtain the selection gradient of traits affecting helping (Selection gradient of traits affecting helping; section 2.8). Finally, we obtain the selection gradient of reproductive effort (Selection gradient of reproductive effort; section 2.9).

#### 2.1 Resident-mutant population dynamics

Having set up some of our general notation, we are ready to write the equations describing the population dynamics of our model, which we let occur in discrete time.

Let $N_{\ell,i}(t)$ denote the number of (dispersed) unmated reproductives of sex $\ell \in \{♀, ♂\}$ and genotype $i \in \{\mathrm{r}, \mathrm{m}\}$ at "ecological" time $t$, so that $N_{♀,\mathrm{r}}(t)$, $N_{♀,\mathrm{m}}(t)$, $N_{♂,\mathrm{r}}(t)$, and $N_{♂,\mathrm{m}}(t)$ represent, respectively, the number of unmated resident females, mutant females, resident males, and mutant males at time $t$. Likewise, let $N_{a,k}(t)$ denote the number of couples of age $a \in \{1,2\}$ and type $k \in \{\mathrm{rr}, \mathrm{rm}, \mathrm{mr}\}$ at time $t$. The variables $N_{\ell,i}$ and $N_{a,k}$ for $\ell \in \{♀, ♂\}$, $i \in \{\mathrm{r}, \mathrm{m}\}$, $a \in \{1,2\}$, and $k \in \{\mathrm{rr}, \mathrm{rm}, \mathrm{mr}\}$ constitute the dynamic variables (ten in total) of the population dynamics part of our model (Fig. S3). We collect these variables in the 10-dimensional vector

$$\mathbf{N}(t) = \begin{pmatrix} \mathbf{N}_{\mathrm{r}}(t) \\ \mathbf{N}_{\mathrm{m}}(t) \end{pmatrix}, \tag{S2.1.1}$$

concatenating the resident and the mutant population vectors, respectively given by

$$\mathbf{N}_{\mathrm{r}}(t) = (N_{♀,\mathrm{r}}(t), N_{♂,\mathrm{r}}(t), N_{1,\mathrm{rr}}(t), N_{2,\mathrm{rr}}(t))^{\intercal}, \tag{S2.1.2}$$

and

$$\mathbf{N}_{\mathrm{m}}(t) = (N_{♀,\mathrm{m}}(t), N_{♂,\mathrm{m}}(t), N_{1,\mathrm{rm}}(t), N_{1,\mathrm{mr}}(t), N_{2,\mathrm{rm}}(t), N_{2,\mathrm{mr}}(t))^{\intercal}. \tag{S2.1.3}$$

We now write down the equations that allow us to project such variables from time $t$ to time $t+1$, and, recursively, to any future time step.

Let

$$\mathcal{N}_k(t) = N_{♀,♀(k)}(t) N_{♂,♂(k)}(t) \tag{S2.1.4}$$

denote the product of unmated females of genotype ♀$(k)$ and unmated males of genotype ♂$(k)$ (see definitions (S1.1.1a) and (S1.1.1b)), which evaluates to

$$\mathcal{N}_{\text{rr}}(t) = N_{♀,\text{r}}(t) N_{♂,\text{r}}(t), \tag{S2.1.5a}$$

$$\mathcal{N}_{\text{rm}}(t) = N_{♀,\text{r}}(t) N_{♂,\text{m}}(t), \tag{S2.1.5b}$$

$$\mathcal{N}_{\text{mr}}(t) = N_{♀,\text{m}}(t) N_{♂,\text{r}}(t). \tag{S2.1.5c}$$

Assuming random mating, the number of matings at time $t$ giving rise to young couples of type $k$ is proportional to $\mathcal{N}_k(t)$. Hence,

$$N_{1,k}(t+1) = \alpha(\mathbf{N}(t)) \mathcal{N}_k(t), \tag{S2.1.6}$$

where $\alpha(\mathbf{N}(t))$ (an expression for which we derive in equation (S2.1.9) below) measures nesting site availability and enforces the density-dependence condition that the total number of couples (i.e., nests) in the population is equal to the total number of nesting sites, $N$, that is,

$$\sum_{k\in\{\text{rr},\text{rm},\text{mr}\}} \sum_{a\in\{1,2\}} N_{a,k}(t+1) = N. \tag{S2.1.7}$$

Each young couple of type $k$ becomes an old couple at the next time step with probability $s_{M,k}$. Hence,

$$N_{2,k}(t+1) = s_{M,k} N_{1,k}(t). \tag{S2.1.8}$$

Substituting (S2.1.6) and (S2.1.8) into (S2.1.7), $\alpha(\mathbf{N}(t))$ in (S2.1.6) can be written in terms of our variables as

$$\alpha(\mathbf{N}(t)) = \frac{N - \sum_{k\in\{\text{rr},\text{rm},\text{mr}\}} s_{M,k} N_{1,k}(t)}{\sum_{k\in\{\text{rr},\text{rm},\text{mr}\}} \mathcal{N}_k(t)}. \tag{S2.1.9}$$

In turn, the number of dispersed unmated individuals of sex $\ell$ and genotype $i$ at time $t+1$ is given by

$$N_{\ell,i}(t+1) = \sum_{k\in\{\text{rr},\text{rm},\text{mr}\}} \sum_{a\in\{1,2\}} N_{a,k}(t) F_{a,\ell,i,k}, \tag{S2.1.10}$$

where $F_{a,\ell,i,k}$ is the expected number of individuals of sex $\ell$ and genotype $i$ that successfully disperse and that are produced by a couple of age $a$ and type $k$. The quantity $F_{a,\ell,i,k}$ is the effective fertility defined in section 1.5 (see expressions (S1.5.1) and (S1.5.3)).

Recursions (S2.1.6), (S2.1.8), and (S2.1.10) describe the population dynamics of our model: recursion (S2.1.6) describes mating, recursion (S2.1.8) describes mated-pair survival, and recursion (S2.1.10) describes reproduction. It is convenient to write this set of equations in matrix notation as

$$\mathbf{N}(t+1) = \mathbf{A}(\mathbf{N}(t)) \mathbf{N}(t), \tag{S2.1.11}$$

where the projection matrix

$$\mathbf{A}(\mathbf{N}(t)) = \begin{pmatrix} \mathbf{A}_{\text{rr}}(\mathbf{N}(t)) & \mathbf{A}_{\text{rm}} \\ \mathbf{A}_{\text{mr}}(\mathbf{N}(t)) & \mathbf{A}_{\text{mm}}(\mathbf{N}(t)) \end{pmatrix}, \tag{S2.1.12}$$

comprises the submatrices

$$\mathbf{A}_{\mathrm{rr}}(\mathbf{N}(t)) = \begin{pmatrix} 0 & 0 & F_{1,♀,\mathrm{r},\mathrm{rr}} & F_{2,♀,\mathrm{r},\mathrm{rr}} \\ 0 & 0 & F_{1,♂,\mathrm{r},\mathrm{rr}} & F_{2,♂,\mathrm{r},\mathrm{rr}} \\ \alpha(\mathbf{N}(t))N_{♂,\mathrm{r}}(t) & \alpha(\mathbf{N}(t))N_{♀,\mathrm{r}}(t) & 0 & 0 \\ 0 & 0 & s_{M,\mathrm{rr}} & 0 \end{pmatrix}, \tag{S2.1.13a}$$

$$\mathbf{A}_{\mathrm{rm}} = \begin{pmatrix} 0 & 0 & F_{1,♀,\mathrm{r},\mathrm{rm}} & F_{1,♀,\mathrm{r},\mathrm{mr}} & F_{2,♀,\mathrm{r},\mathrm{rm}} & F_{2,♀,\mathrm{r},\mathrm{mr}} \\ 0 & 0 & F_{1,♂,\mathrm{r},\mathrm{rm}} & F_{1,♂,\mathrm{r},\mathrm{mr}} & F_{2,♂,\mathrm{r},\mathrm{rm}} & F_{2,♂,\mathrm{r},\mathrm{mr}} \\ 0 & 0 & 0 & 0 & 0 & 0 \\ 0 & 0 & 0 & 0 & 0 & 0 \end{pmatrix}, \tag{S2.1.13b}$$

$$\mathbf{A}_{\mathrm{mr}}(\mathbf{N}(t)) = \begin{pmatrix} 0 & 0 & 0 & 0 \\ 0 & 0 & 0 & 0 \\ \alpha(\mathbf{N}(t))N_{♂,\mathrm{m}}(t) & 0 & 0 & 0 \\ 0 & \alpha(\mathbf{N}(t))N_{♀,\mathrm{m}}(t) & 0 & 0 \\ 0 & 0 & 0 & 0 \\ 0 & 0 & 0 & 0 \end{pmatrix}, \tag{S2.1.13c}$$

$$\mathbf{A}_{\mathrm{mm}}(\mathbf{N}(t)) = \begin{pmatrix} 0 & 0 & F_{1,♀,\mathrm{m},\mathrm{rm}} & F_{1,♀,\mathrm{m},\mathrm{mr}} & F_{2,♀,\mathrm{m},\mathrm{rm}} & F_{2,♀,\mathrm{m},\mathrm{mr}} \\ 0 & 0 & F_{1,♂,\mathrm{m},\mathrm{rm}} & F_{1,♂,\mathrm{m},\mathrm{mr}} & F_{2,♂,\mathrm{m},\mathrm{rm}} & F_{2,♂,\mathrm{m},\mathrm{mr}} \\ 0 & \alpha(\mathbf{N}(t))N_{♀,\mathrm{r}}(t) & 0 & 0 & 0 & 0 \\ \alpha(\mathbf{N}(t))N_{♂,\mathrm{r}}(t) & 0 & 0 & 0 & 0 & 0 \\ 0 & 0 & s_{M,\mathrm{rm}} & 0 & 0 & 0 \\ 0 & 0 & 0 & s_{M,\mathrm{mr}} & 0 & 0 \end{pmatrix}. \tag{S2.1.13d}$$

#### 2.2 Resident population dynamics and resident equilibrium

In the absence of the mutant allele, $\mathbf{N}_{\mathrm{m}}(t) = (0,\dots,0)^{\intercal}$ holds, and the population dynamics (S2.1.11) reduces to the resident system

$$\mathbf{N}_{\mathrm{r}}(t+1) = \mathbf{A}_{\mathrm{rr}}(\mathbf{N}_{\mathrm{r}}(t))\mathbf{N}_{\mathrm{r}}(t), \tag{S2.2.1}$$

with

$$\alpha(\mathbf{N}_{\mathrm{r}}(t)) = \frac{N - s_{M,\mathrm{rr}}N_{1,\mathrm{rr}}(t)}{\mathcal{N}_{\mathrm{rr}}(t)}. \tag{S2.2.2}$$

Substituting (S2.2.2) into the projection matrix $\mathbf{A}_{\mathrm{rr}}(\mathbf{N}_{\mathrm{r}}(t))$ (S2.1.13a), performing the matrix multiplication in (S2.2.1), and simplifying, yields

$$N_{♀,\mathrm{r}}(t+1) = F_{1,♀,\mathrm{r},\mathrm{rr}}N_{1,\mathrm{rr}}(t) + F_{2,♀,\mathrm{r},\mathrm{rr}}N_{2,\mathrm{rr}}(t), \tag{S2.2.3a}$$

$$N_{♂,\mathrm{r}}(t+1) = F_{1,♂,\mathrm{r},\mathrm{rr}}N_{1,\mathrm{rr}}(t) + F_{2,♂,\mathrm{r},\mathrm{rr}}N_{2,\mathrm{rr}}(t), \tag{S2.2.3b}$$

$$N_{1,\mathrm{rr}}(t+1) = N - s_{M,\mathrm{rr}}N_{1,\mathrm{rr}}(t), \tag{S2.2.3c}$$

$$N_{2,\mathrm{rr}}(t+1) = s_{M,\mathrm{rr}}N_{1,\mathrm{rr}}(t). \tag{S2.2.3d}$$

At an equilibrium $\mathbf{N}_{\mathrm{r}}^* = (N^*_{♀,\mathrm{r}}, N^*_{♂,\mathrm{r}}, N^*_{1,\mathrm{rr}}, N^*_{2,\mathrm{rr}})^\intercal$, the system satisfies

$$N_{♀,\mathrm{r}}(t+1) = N_{♀,\mathrm{r}}(t) = N^*_{♀,\mathrm{r}}, \tag{S2.2.4a}$$

$$N_{♂,\mathrm{r}}(t+1) = N_{♂,\mathrm{r}}(t) = N^*_{♂,\mathrm{r}}, \tag{S2.2.4b}$$

$$N_{1,\mathrm{rr}}(t+1) = N_{1,\mathrm{rr}}(t) = N^*_{1,\mathrm{rr}}, \tag{S2.2.4c}$$

$$N_{2,\mathrm{rr}}(t+1) = N_{2,\mathrm{rr}}(t) = N^*_{2,\mathrm{rr}}. \tag{S2.2.4d}$$

Substituting (S2.2.4) into (S2.2.3) and solving the resulting linear system of equations, we find that the system admits a unique equilibrium given by

$$N^*_{♀,\mathrm{r}} = \frac{N}{1+s_{M,\mathrm{rr}}}\left(F_{1,♀,\mathrm{r,rr}} + s_{M,\mathrm{rr}} F_{2,♀,\mathrm{r,rr}}\right) = \frac{N}{1+s_{M,\mathrm{rr}}}\Pi_{♀,\mathrm{r,rr}}, \tag{S2.2.5a}$$

$$N^*_{♂,\mathrm{r}} = \frac{N}{1+s_{M,\mathrm{rr}}}\left(F_{1,♂,\mathrm{r,rr}} + s_{M,\mathrm{rr}} F_{2,♂,\mathrm{r,rr}}\right) = \frac{N}{1+s_{M,\mathrm{rr}}}\Pi_{♂,\mathrm{r,rr}}, \tag{S2.2.5b}$$

$$N^*_{1,\mathrm{rr}} = \frac{N}{1+s_{M,\mathrm{rr}}}, \tag{S2.2.5c}$$

$$N^*_{2,\mathrm{rr}} = \frac{N}{1+s_{M,\mathrm{rr}}} s_{M,\mathrm{rr}}, \tag{S2.2.5d}$$

where the second equality in expressions (S2.2.5a) and (S2.2.5b) follows from identity (S1.6.1), which links effective fertilities and productivities.

This equilibrium is locally stable. To show this, we perform a local stability analysis [1] of the resident system (S2.2.1) at the resident equilibrium (S2.2.5). Evaluating the Jacobian matrix of (S2.2.1) at (S2.2.5) we obtain the local stability matrix

$$\mathbf{J}_{\mathrm{res}} = \left.\left(\frac{\partial \mathbf{N}_{\mathrm{r}}(t+1)}{\partial N_{♀,\mathrm{r}}(t)}, \frac{\partial \mathbf{N}_{\mathrm{r}}(t+1)}{\partial N_{♂,\mathrm{r}}(t)}, \frac{\partial \mathbf{N}_{\mathrm{r}}(t+1)}{\partial N_{1,\mathrm{rr}}(t)}, \frac{\partial \mathbf{N}_{\mathrm{r}}(t+1)}{\partial N_{2,\mathrm{rr}}(t)}\right)\right|_{\mathbf{N}_{\mathrm{r}}=\mathbf{N}_{\mathrm{r}}^*} \tag{S2.2.6a}$$

$$= \begin{pmatrix} 0 & 0 & F_{1,♀,\mathrm{r,rr}} & F_{2,♀,\mathrm{r,rr}} \\ 0 & 0 & F_{1,♂,\mathrm{r,rr}} & F_{2,♂,\mathrm{r,rr}} \\ 0 & 0 & -s_{M,\mathrm{rr}} & 0 \\ 0 & 0 & s_{M,\mathrm{rr}} & 0 \end{pmatrix}. \tag{S2.2.6b}$$

This matrix has a block-triangular form composed of four 2 × 2 submatrices; because of this block-triangular form, the eigenvalues of $\mathbf{J}_{\mathrm{res}}$ correspond to the eigenvalues of the submatrices along the diagonal. As these submatrices are both triangular, their eigenvalues are the values along their main diagonals. It follows that the eigenvalues of $\mathbf{J}_{\mathrm{res}}$ are zero (with multiplicity three) and $-s_{M,\mathrm{rr}}$. Since we assume that $s_{M,\mathrm{rr}} < 1$, the absolute value of the leading eigenvalue of $\mathbf{J}_{\mathrm{res}}$ is less than one, proving the local stability of $\mathbf{N}_{\mathrm{r}}^*$. We conclude that the resident equilibrium is locally stable in the absence of the mutant allele.

From (S2.2.5a) and (S2.2.5b), we have that the sex ratio among unmated reproductives at the resident equilibrium is given by the ratio of sex-specific productivities, that is,

$$\frac{N^*_{♂,\mathrm{r}}}{N^*_{♀,\mathrm{r}}} = \frac{\Pi_{♂,\mathrm{r,rr}}}{\Pi_{♀,\mathrm{r,rr}}}. \tag{S2.2.7}$$

#### 2.3 Invasion fitness

We now identify invasion fitness, that is, the asymptotic growth rate of a rare mutant population introduced at the resident equilibrium

$$\mathbf{N}^* = (\mathbf{N}_\mathrm{r}^*, \mathbf{0}), \tag{S2.3.1}$$

where $\mathbf{N}_\mathrm{r}^*$ corresponds to (S2.2.5). To a first-order approximation, the population dynamics around the resident equilibrium are governed by the local stability matrix

$$\mathbf{J} = \left( \frac{\partial \mathbf{N}(t+1)}{\partial N_{♀,\mathrm{r}}(t)}, \frac{\partial \mathbf{N}(t+1)}{\partial N_{♂,\mathrm{r}}(t)}, \ldots, \frac{\partial \mathbf{N}(t+1)}{\partial N_{2,\mathrm{rm}}(t)}, \frac{\partial \mathbf{N}(t+1)}{\partial N_{2,\mathrm{mr}}(t)} \right) \Bigg|_{\mathbf{N}=\mathbf{N}^*}, \tag{S2.3.2}$$

that is, the Jacobian matrix of (S2.1.11) evaluated at the resident equilibrium (S2.3.1). Taking the partial derivatives, it can be checked that this Jacobian has the block-triangular form [1]:

$$\mathbf{J} = \begin{pmatrix} \mathbf{J}_\mathrm{res} & \mathbf{V} \\ \mathbf{0} & \mathbf{J}_\mathrm{mut} \end{pmatrix}, \tag{S2.3.3}$$

featuring submatrices $\mathbf{0}$ (a $6 \times 4$ matrix of zeros), $\mathbf{J}_\mathrm{res}$ (the $4 \times 4$ matrix given by equation (S2.2.6b)), $\mathbf{V}$ (a $4 \times 6$ matrix), and

$$\mathbf{J}_\mathrm{mut} = \begin{pmatrix} 0 & 0 & F_{1,♀,\mathrm{m,rm}} & F_{1,♀,\mathrm{m,mr}} & F_{2,♀,\mathrm{m,rm}} & F_{2,♀,\mathrm{m,mr}} \\ 0 & 0 & F_{1,♂,\mathrm{m,rm}} & F_{1,♂,\mathrm{m,mr}} & F_{2,♂,\mathrm{m,rm}} & F_{2,♂,\mathrm{m,mr}} \\ 0 & \dfrac{1}{\Pi_{♂,\mathrm{r,rr}}} & 0 & 0 & 0 & 0 \\ \dfrac{1}{\Pi_{♀,\mathrm{r,rr}}} & 0 & 0 & 0 & 0 & 0 \\ 0 & 0 & s_{M,\mathrm{rm}} & 0 & 0 & 0 \\ 0 & 0 & 0 & s_{M,\mathrm{mr}} & 0 & 0 \end{pmatrix} \tag{S2.3.4}$$

(a $6 \times 6$ matrix). Given the block-triangular form of $\mathbf{J}$ (S2.3.3), the mutant submatrix $\mathbf{J}_\mathrm{mut}$ governs the mutant population dynamics around the resident equilibrium.

Invasion fitness is given by the leading eigenvalue $\lambda$ of $\mathbf{J}_\mathrm{mut}$. Since raising $\mathbf{J}_\mathrm{mut}$ to a sufficiently high power yields matrices with all entries being positive, $\mathbf{J}_\mathrm{mut}$ is nonnegative, irreducible, and primitive. It follows from the Perron-Frobenius theorem that $\lambda$ is real and positive [9], and that invasion fitness is well defined. Then, a rare mutant allele invades if and only if the absolute value of the invasion fitness is larger than one.

#### 2.4 Selection gradient (generic form)

We now use our identification of invasion fitness to obtain a general expression of the selection gradient, which gives the direction of selection. Invasion fitness can be written as $\lambda = \lambda(\mathbf{z}_\mathrm{m}, \mathbf{z})$ to highlight the fact that it is a function of both mutant and resident phenotypes because so are the entries of $\mathbf{J}_\mathrm{mut}$. Here, $\mathbf{z}_\mathrm{m} = (\zeta_\mathrm{m})^\intercal = (p_\mathrm{m}, z_\mathrm{m})^\intercal$ and $\mathbf{z} = (\zeta)^\intercal = (p, z)^\intercal$ for model cases of offspring or maternal control, or $\mathbf{z}_\mathrm{m} = (\zeta_\mathrm{m})^\intercal = (x_\mathrm{m}, y_\mathrm{m}, z_\mathrm{m})^\intercal$ and $\mathbf{z} = (\zeta)^\intercal = (x, y, z)^\intercal$ for model cases of shared control.

We assume that mutations have small phenotypic effects (i.e., we assume that selection is $\delta$-weak; [11]). Then, invasion fitness can be approximated by a first-order Taylor expansion of $\lambda(\mathbf{z}_\mathrm{m}, \mathbf{z})$ with respect to $\mathbf{z}_\mathrm{m}$

around $\mathbf{z}$ to obtain

$$\lambda(\mathbf{z}_{\mathrm{m}},\mathbf{z}) \approx 1 + (\mathbf{z}_{\mathrm{m}} - \mathbf{z})^{\mathsf{T}}\boldsymbol{\mathcal{S}}(\mathbf{z}),$$

where we have used the fact that $\lambda(\mathbf{z},\mathbf{z}) = 1$ (since mutant alleles coding for the same trait as the resident are neutral), and where the selection gradient of $\mathbf{z}$ is given by

$$\boldsymbol{\mathcal{S}}(\mathbf{z}) = \begin{pmatrix} \mathcal{S}_p(\mathbf{z}) \\ \mathcal{S}_z(\mathbf{z}) \end{pmatrix} = \begin{pmatrix} \left.\dfrac{\partial\lambda}{\partial p_{\mathrm{m}}}\right|_{\mathbf{z}_{\mathrm{m}}=\mathbf{z}} \\ \left.\dfrac{\partial\lambda}{\partial z_{\mathrm{m}}}\right|_{\mathbf{z}_{\mathrm{m}}=\mathbf{z}} \end{pmatrix}, \tag{S2.4.1}$$

for model cases of offspring and maternal control, or by

$$\boldsymbol{\mathcal{S}}(\mathbf{z}) = \begin{pmatrix} \mathcal{S}_x(\mathbf{z}) \\ \mathcal{S}_y(\mathbf{z}) \\ \mathcal{S}_z(\mathbf{z}) \end{pmatrix} = \begin{pmatrix} \left.\dfrac{\partial\lambda}{\partial x_{\mathrm{m}}}\right|_{\mathbf{z}_{\mathrm{m}}=\mathbf{z}} \\ \left.\dfrac{\partial\lambda}{\partial y_{\mathrm{m}}}\right|_{\mathbf{z}_{\mathrm{m}}=\mathbf{z}} \\ \left.\dfrac{\partial\lambda}{\partial z_{\mathrm{m}}}\right|_{\mathbf{z}_{\mathrm{m}}=\mathbf{z}} \end{pmatrix} \tag{S2.4.2}$$

for model cases of shared control.

To calculate the selection gradient of $\zeta$, $\mathcal{S}_\zeta(\mathbf{z})$, (where $\zeta \in \{p,z\}$ for offspring and maternal control; $\zeta \in \{x,y,z\}$ for shared control), that is,

$$\mathcal{S}_\zeta(\mathbf{z}) = \left.\frac{\partial\lambda}{\partial\zeta_{\mathrm{m}}}\right|_{\mathbf{z}_{\mathrm{m}}=\mathbf{z}}, \tag{S2.4.3}$$

we use a classic result on perturbations of the leading eigenvalue of irreducible and nonnegative matrices. This result implies that the selection gradient of $\zeta$ (S2.4.3) can be written as [9, 1]

$$\mathcal{S}_\zeta(\mathbf{z}) = \frac{\mathbf{v}^{\mathsf{T}} \left.\dfrac{\partial\mathbf{J}_{\mathrm{mut}}}{\partial\zeta_{\mathrm{m}}}\right|_{\mathbf{z}_{\mathrm{m}}=\mathbf{z}} \mathbf{u}}{\mathbf{v}^{\mathsf{T}}\mathbf{u}}, \tag{S2.4.4}$$

where $\mathbf{v}$ and $\mathbf{u}$ are, respectively, the left and right eigenvectors associated to the leading eigenvalue of the neutral mutant submatrix $\mathbf{J}^\circ_{\mathrm{mut}}$, which equals one. Henceforth, we will denote by $X^\circ$ a variable $X$ considered under neutrality, that is

$$X^\circ \equiv X|_{\mathbf{z}_{\mathrm{m}}=\mathbf{z}}, \tag{S2.4.5}$$

for any variable $X$. With this convention,

$$\mathbf{J}^\circ_{\mathrm{mut}} \equiv \mathbf{J}_{\mathrm{mut}}|_{\mathbf{z}_{\mathrm{m}}=\mathbf{z}} \tag{S2.4.6}$$

$$= \begin{pmatrix} 0 & 0 & F^\circ_{1,♀,\mathrm{m,rm}} & F^\circ_{1,♀,\mathrm{m,mr}} & F^\circ_{2,♀,\mathrm{m,rm}} & F^\circ_{2,♀,\mathrm{m,mr}} \\ 0 & 0 & F^\circ_{1,♂,\mathrm{m,rm}} & F^\circ_{1,♂,\mathrm{m,mr}} & F^\circ_{2,♂,\mathrm{m,rm}} & F^\circ_{2,♂,\mathrm{m,mr}} \\ 0 & \dfrac{1}{\Pi^\circ_{♂,\mathrm{r,rr}}} & 0 & 0 & 0 & 0 \\ \dfrac{1}{\Pi^\circ_{♀,\mathrm{r,rr}}} & 0 & 0 & 0 & 0 & 0 \\ 0 & 0 & s^\circ_{M,\mathrm{rm}} & 0 & 0 & 0 \\ 0 & 0 & 0 & s^\circ_{M,\mathrm{mr}} & 0 & 0 \end{pmatrix}. \tag{S2.4.7}$$

#### 2.5 Neutral mutant submatrix, $\mathbf{J}^{\circ}_{\text{mut}}$

To calculate the dominant left and right eigenvectors **v** and **u** of the neutral mutant submatrix $\mathbf{J}^{\circ}_{\text{mut}}$, we now calculate the entries of $\mathbf{J}^{\circ}_{\text{mut}}$, together with other variables and rates considered under neutrality. All of these neutral variables and rates can be written in terms of variables and rates of resident individuals in a resident population.

**Neutral reproductive effort, $z^{\circ}_k$.** We start by calculating $z^{\circ}_k$, that is, the neutral reproductive effort exerted by the female of an old couple of type $k$. For all $k \in \{\text{rr}, \text{rm}, \text{mr}\}$, this is given by

$$z^{\circ}_k \equiv z_k|_{\mathbf{z}_\text{m}=\mathbf{z}} = z_{♀(k)}\big|_{\mathbf{z}_\text{m}=\mathbf{z}} = z = z_{\text{rr}}, \tag{S2.5.1}$$

where the first equality follows from the definitions of $z_k$ (S1.1.2a) and neutrality (S2.4.5); the second equality follows from the definition of $♀(k)$ (S1.1.1a) and our notational convention (S1.1.2b); and the last equality follows again from our convention (S1.1.2b).

**Neutral expected number of helpers, $h^{\circ}_k$.** We proceed now to calculate $h^{\circ}_k$, that is, the expected number of helpers for an old couple of type $k$ evaluated at neutrality. Let us first note that, by inspection of the values of the helping probabilities given in Figure S5, the following identity holds:

$$p^{\circ}_{\ell,i,k} \equiv p_{\ell,i,k}\big|_{\mathbf{z}_\text{m}=\mathbf{z}} = p_{\ell,\text{r},\text{rr}}. \tag{S2.5.2}$$

Taking this into account, we can then write, for all $k \in \{\text{rr}, \text{rm}, \text{mr}\}$,

$$
\begin{aligned}
h^{\circ}_k &= \left( f_1 \sum_{\ell \in \{♀,♂\}} \sigma_{1,\ell} \sum_{i \in \{\text{r},\text{m}\}} q_{\ell,i,k} p_{\ell,i,k} \right)\Bigg|_{\mathbf{z}_\text{m}=\mathbf{z}} && \text{(S2.5.3a)}\\
&= f_1 \sum_{\ell \in \{♀,♂\}} \sigma_{1,\ell} \sum_{i \in \{\text{r},\text{m}\}} q_{\ell,i,k} \left(p_{\ell,i,k}\right)\big|_{\mathbf{z}_\text{m}=\mathbf{z}} && \text{(S2.5.3b)}\\
&= f_1 \sum_{\ell \in \{♀,♂\}} \sigma_{1,\ell} \sum_{i \in \{\text{r},\text{m}\}} q_{\ell,i,k} p_{\ell,\text{r},\text{rr}} && \text{(S2.5.3c)}\\
&= f_1 \sum_{\ell \in \{♀,♂\}} \sigma_{1,\ell} p_{\ell,\text{r},\text{rr}} \sum_{i \in \{\text{r},\text{m}\}} q_{\ell,i,k} && \text{(S2.5.3d)}\\
&= f_1 \sum_{\ell \in \{♀,♂\}} \sigma_{1,\ell} p_{\ell,\text{r},\text{rr}} && \text{(S2.5.3e)}\\
&= h_{\text{rr}} = h && \text{(S2.5.3f)}
\end{aligned}
$$

where the first line (S2.5.3b) follows from substituting (S1.3.3b) and the definition of neutrality (S2.4.5); the second line (S2.5.3b) follows from the fact that only the probabilities $p_{\ell,i,k}$ are functions of the evolving traits **z**; the third line (S2.5.3c) applies identity (S2.5.2); the fifth line (S2.5.3e) applies identity (S1.2.1c); and the final line (S2.5.3f) follows from (S1.3.4b).

**Neutral vital rates ($f^{\circ}_{2,k}$, $s^{\circ}_{M,k}$, and $s^{\circ}_{2,k}$).** The entries of $\mathbf{J}^{\circ}_{\text{mut}}$ as given in equation (S2.4.6) depend on the values of the different vital rates under neutrality, that is, on $f^{\circ}_{2,k}$, $s^{\circ}_{M,k}$, and $s^{\circ}_{2,k}$. We calculate these values now.

The late fertility of the female of a couple of type $k$ under neutrality is given by

$$f^{\circ}_{2,k} = f_2(z_k)\big|_{\mathbf{z}_\mathrm{m}=\mathbf{z}} = f_2\left(z_k|_{\mathbf{z}_\mathrm{m}=\mathbf{z}}\right) = f_2(z_\mathrm{rr}) = f_{2,\mathrm{rr}}, \tag{S2.5.4}$$

where the first equality follows from substituting equation (S1.4.1) and from the definition of neutrality (S2.4.5); the second equality holds because the function $f_2$ (S1.4.2) is the same for all $k$; the third equality follows from equation (S2.5.1); and the last equality follows from (S1.4.1) with $k = \mathrm{rr}$.

The survival of a couple of type $k$ under neutrality is given by

$$s^{\circ}_{M,k} = s_M(f_{2,k}, h_k)\big|_{\mathbf{z}_\mathrm{m}=\mathbf{z}} = s_M\left(f^{\circ}_{2,k}, h^{\circ}_k\right) = s_M\left(f_{2,\mathrm{rr}}, h_\mathrm{rr}\right) = s_{M,\mathrm{rr}}, \tag{S2.5.5}$$

where the first equality follows from substituting equation (S1.4.4a) and from the definition of neutrality (S2.4.5); the second equality holds because the function $s_M$ (S1.4.5a) is the same for all $k$; the third equality follows from equation (S2.5.4) and (S2.5.3f); and the last equality follows from (S1.4.4a) with $k = \mathrm{rr}$. Thus, the probabilities $s^{\circ}_{M,\mathrm{rm}}$ and $s^{\circ}_{M,\mathrm{mr}}$ featuring in $\mathbf{J}^{\circ}_\mathrm{mut}$ (S2.4.6) simplify to

$$s^{\circ}_{M,\mathrm{rm}} = s^{\circ}_{M,\mathrm{mr}} = s_{M,\mathrm{rr}}. \tag{S2.5.6}$$

Analogous reasoning leads to the following expression for the survival of the second-brood offspring of a couple of type $k$ under neutrality:

$$s^{\circ}_{2,k} = s_2(f_{2,k}, h_k)\big|_{\mathbf{z}_\mathrm{m}=\mathbf{z}} = s_2\left(f^{\circ}_{2,k}, h^{\circ}_k\right) = s_2\left(f_{2,\mathrm{rr}}, h_\mathrm{rr}\right) = s_{2,\mathrm{rr}}. \tag{S2.5.7}$$

**Neutral effective fertility, $F^{\circ}_{a,\ell,i,k}$.** The nonzero entries in the first two rows of $\mathbf{J}^{\circ}_\mathrm{mut}$ (S2.4.6) are effective fertilities (defined in section 1.5) under neutrality. We find explicit expressions for these effective fertilities below.

First, for all $\ell$, all $i$, and all $k$, the early effective fertility under neutrality, $F^{\circ}_{1,\ell,i,k}$, simplifies to

$$\begin{aligned}
F^{\circ}_{1,\ell,i,k} &= \left(f_1\sigma_{1,\ell}q_{\ell,i,k}(1-p_{\ell,i,k})s_1\right)\big|_{\mathbf{z}_\mathrm{m}=\mathbf{z}} \\
&= q_{\ell,i,k}f_1\sigma_{1,\ell}(1-p_{\ell,\mathrm{r},\mathrm{rr}})s_1 \\
&= q_{\ell,i,k}F_{1,\ell,\mathrm{r},\mathrm{rr}}
\end{aligned} \tag{S2.5.8}$$

where the first equality follows from substituting the expression for $F_{1,\ell,i,k}$ (S1.5.1) and the definition of neutrality (S2.4.5); the second equality follows from (S2.5.2); and the final equality follows from (S1.5.2).

Likewise, for all $\ell$, all $i$, and all $k$, the late effective fertility under neutrality, $F^{\circ}_{2,\ell,i,k}$, simplifies to

$$\begin{aligned}
F^{\circ}_{2,\ell,i,k} &= \left(f_{2,k}\sigma_{2,\ell}q_{\ell,i,k}s_{2,k}\right)\big|_{\mathbf{z}_\mathrm{m}=\mathbf{z}} \\
&= q_{\ell,i,k}\sigma_{2,\ell}f^{\circ}_{2,k}s^{\circ}_{2,k} \\
&= q_{\ell,i,k}\sigma_{2,\ell}f_{2,\mathrm{rr}}s_{2,\mathrm{rr}} \\
&= q_{\ell,i,k}F_{2,\ell,\mathrm{r},\mathrm{rr}}
\end{aligned} \tag{S2.5.9}$$

where we have substituted the expressions for $f^{\circ}_{2,k}$ and $s^{\circ}_{2,k}$ given by equations (S2.5.4) and (S2.5.7), and the expression for $F_{2,\ell,\mathrm{r},\mathrm{rr}}$ given by (S1.5.4).

Equations (S2.5.8) and (S2.5.9) state that the effective fertility of a young or old couple that has a neutral mutation equals the corresponding effective fertility of a resident couple multiplied by the probability that the mutant couple produces an offspring of the relevant genotype and relevant sex.

**Neutral productivity, $\Pi^{\circ}_{\ell,i,k}$.** When simplifying the expression for the selection gradient, it will be useful to have the expression for the neutral productivity $\Pi^{\circ}_{\ell,i,k}$ of a $k$-type couple through offspring of sex $\ell$ and genotype $i$. To calculate it, we start from the expression for $\Pi_{\ell,i,k}$ (equation (S1.6.1)), evaluate at neutrality, and simplify using the expressions for the neutral effective fertilities (equations (S2.5.8) and (S2.5.9)) and couple survival (equation (S2.5.5)) to obtain

$$\begin{aligned}\Pi^{\circ}_{\ell,i,k} &= \left(F_{1,\ell,i,k} + s_{M,k}F_{2,\ell,i,k}\right)\big|_{\mathbf{z}_{\mathrm{m}}=\mathbf{z}} \\ &= F^{\circ}_{1,\ell,i,k} + s^{\circ}_{M,k}F^{\circ}_{2,\ell,i,k} \\ &= q_{\ell,i,k}\left(F_{1,\ell,\mathrm{r},\mathrm{rr}} + s_{M,\mathrm{rr}}F_{2,\ell,\mathrm{r},\mathrm{rr}}\right) \\ &= q_{\ell,i,k}\Pi_{\ell,\mathrm{r},\mathrm{rr}} \end{aligned} \tag{S2.5.10}$$

where the last line follows from identifying the expression for the productivity of a couple of type rr through resident offspring, $\Pi_{\ell,\mathrm{r},\mathrm{rr}}$ given by equation (S1.6.18). In particular, and because of identity (S1.2.1a) we recover

$$\Pi^{\circ}_{\ell,\mathrm{r},\mathrm{rr}} = \Pi_{\ell,\mathrm{r},\mathrm{rr}}. \tag{S2.5.11}$$

**Neutral mutant submatrix, $\mathbf{J}^{\circ}_{\mathrm{mut}}$.** Putting together our previous results in this subsection 2.5, we write the neutral mutant submatrix, $\mathbf{J}^{\circ}_{\mathrm{mut}}$ (S2.4.6) as

$$\begin{aligned}\mathbf{J}^{\circ}_{\mathrm{mut}} &\equiv \mathbf{J}_{\mathrm{mut}}\big|_{\mathbf{z}_{\mathrm{m}}=\mathbf{z}} \\ &= \begin{pmatrix} 0 & 0 & q_{♀,\mathrm{m},\mathrm{rm}}F_{1,♀,\mathrm{r},\mathrm{rr}} & q_{♀,\mathrm{m},\mathrm{mr}}F_{1,♀,\mathrm{r},\mathrm{rr}} & q_{♀,\mathrm{m},\mathrm{rm}}F_{2,♀,\mathrm{r},\mathrm{rr}} & q_{♀,\mathrm{m},\mathrm{mr}}F_{2,♀,\mathrm{r},\mathrm{rr}} \\ 0 & 0 & q_{♂,\mathrm{m},\mathrm{rm}}F_{1,♂,\mathrm{r},\mathrm{rr}} & q_{♂,\mathrm{m},\mathrm{mr}}F_{1,♂,\mathrm{r},\mathrm{rr}} & q_{♂,\mathrm{m},\mathrm{rm}}F_{2,♂,\mathrm{r},\mathrm{rr}} & q_{♂,\mathrm{m},\mathrm{mr}}F_{2,♂,\mathrm{r},\mathrm{rr}} \\ 0 & \frac{1}{\Pi_{♂,\mathrm{r},\mathrm{rr}}} & 0 & 0 & 0 & 0 \\ \frac{1}{\Pi_{♀,\mathrm{r},\mathrm{rr}}} & 0 & 0 & 0 & 0 & 0 \\ 0 & 0 & s_{M,\mathrm{rr}} & 0 & 0 & 0 \\ 0 & 0 & 0 & s_{M,\mathrm{rr}} & 0 & 0 \end{pmatrix}. \end{aligned} \tag{S2.5.12}$$

#### 2.6 Reproductive values and stable distribution

Having calculated the neutral mutant submatrix, $\mathbf{J}^{\circ}_{\mathrm{mut}}$, we are ready to calculate its (dominant) left and right eigenvectors. These are the vectors $\mathbf{v}$ (S2.6.1) and $\mathbf{u}$ (S2.6.14) appearing in our expression for the selection gradient $\mathcal{S}_{\zeta}(\mathbf{z})$ of a generic trait $\zeta$ given by equation (S2.4.4). The biological meaning of these vectors is the following [1]. The left eigenvector $\mathbf{v}$ lists the reproductive values of neutral mutants, with reproductive values measuring the relative long-term contribution of individuals in a mutant class to the future mutant population. The right eigenvector $\mathbf{u}$ is the stable class distribution of neutral mutants, which measures the relative asymptotic distribution of neutral mutants among classes. By the Perron-Frobenius theorem, the vectors $\mathbf{u}$ and $\mathbf{v}$ are positive [9]. We will show that the selection gradient (S2.4.4) can be simplified so that it only depends on two entries of $\mathbf{u}$ (namely, $u_{1,\mathrm{rm}}$ and $u_{1,\mathrm{mr}}$) and two entries of $\mathbf{v}$ (namely, $v_{♀,\mathrm{m}}$ and $v_{♂,\mathrm{m}}$). Thus, without loss of generality, we choose $\mathbf{u}$ and $\mathbf{v}$ so that $u_{1,\mathrm{rm}} + u_{1,\mathrm{mr}} = 1$ (i.e., $u_{1,k}$ is the stable proportion of mutant young couples of type $k$) and $v_{♂,\mathrm{m}} = 1$ (i.e., the reproductive value of mutant males is arbitrarily set to one). Doing so we slightly depart from common use in demographic models, where $\mathbf{u}$ is often chosen so that the whole vector

$\mathbf{u}$ is a probability distribution, that is, so that $\mathbf{1}^\mathsf{T}\mathbf{u} = 1$ (where $\mathbf{1}$ is a vector of ones), and where $\mathbf{v}$ is sometimes chosen so that the whole vector $\mathbf{v}^\mathsf{T}\mathbf{u}$ is a probability distribution, that is, so that $\mathbf{v}^\mathsf{T}\mathbf{u} = 1$. Regardless, we will continue referring to the vector $\mathbf{u}$ as the stable *distribution*.

**Reproductive values, v.** We start by calculating the left eigenvector

$$\mathbf{v}^\mathsf{T} = \left(v_{♀,\mathrm{m}}, v_{♂,\mathrm{m}}, v_{1,\mathrm{rm}}, v_{1,\mathrm{mr}}, v_{2,\mathrm{rm}}, v_{2,\mathrm{mr}}\right), \tag{S2.6.1}$$

giving the neutral reproductive values of mutants in each class. From the definition of a left eigenvector, and since the leading eigenvalue of $\mathbf{J}^\circ_{\mathrm{mut}}$ is one, $\mathbf{v}$ is defined by

$$\mathbf{v}^\mathsf{T}\mathbf{J}^\circ_{\mathrm{mut}} = \mathbf{v}^\mathsf{T}. \tag{S2.6.2}$$

Performing the matrix multiplication stated in (S2.6.2) with $\mathbf{J}^\circ_{\mathrm{mut}}$ given by equation (S2.4.6), we obtain the system of equations

$$v_{♀,\mathrm{m}} = \frac{v_{1,\mathrm{mr}}}{\Pi^\circ_{♀,\mathrm{r},\mathrm{rr}}}, \tag{S2.6.3a}$$

$$v_{♂,\mathrm{m}} = \frac{v_{1,\mathrm{rm}}}{\Pi^\circ_{♂,\mathrm{r},\mathrm{rr}}}, \tag{S2.6.3b}$$

$$v_{1,\mathrm{rm}} = F^\circ_{1,♀,\mathrm{m},\mathrm{rm}} v_{♀,\mathrm{m}} + F^\circ_{1,♂,\mathrm{m},\mathrm{rm}} v_{♂,\mathrm{m}} + s^\circ_{M,\mathrm{rm}} v_{2,\mathrm{rm}}, \tag{S2.6.3c}$$

$$v_{1,\mathrm{mr}} = F^\circ_{1,♀,\mathrm{m},\mathrm{mr}} v_{♀,\mathrm{m}} + F^\circ_{1,♂,\mathrm{m},\mathrm{mr}} v_{♂,\mathrm{m}} + s^\circ_{M,\mathrm{mr}} v_{2,\mathrm{mr}}, \tag{S2.6.3d}$$

$$v_{2,\mathrm{rm}} = F^\circ_{2,♀,\mathrm{m},\mathrm{rm}} v_{♀,\mathrm{m}} + F^\circ_{2,♂,\mathrm{m},\mathrm{rm}} v_{♂,\mathrm{m}}, \tag{S2.6.3e}$$

$$v_{2,\mathrm{mr}} = F^\circ_{2,♀,\mathrm{m},\mathrm{mr}} v_{♀,\mathrm{m}} + F^\circ_{2,♂,\mathrm{m},\mathrm{mr}} v_{♂,\mathrm{m}}. \tag{S2.6.3f}$$

From these equations we can write down two equivalent expressions for the reproductive values of young mutant couples ($v_{1,\mathrm{mr}}$ and $v_{1,\mathrm{rm}}$) in terms of the reproductive values of mutant unmated reproductives ($v_{♀,\mathrm{m}}$ and $v_{♂,\mathrm{m}}$). First, isolating $v_{1,\mathrm{mr}}$ and $v_{1,\mathrm{rm}}$ from, respectively, equations (S2.6.3a) and (S2.6.3b), and using (S2.5.11), we obtain

$$v_{1,\mathrm{mr}} = \Pi_{♀,\mathrm{r},\mathrm{rr}} v_{♀,\mathrm{m}}, \tag{S2.6.4a}$$

$$v_{1,\mathrm{rm}} = \Pi_{♂,\mathrm{r},\mathrm{rr}} v_{♂,\mathrm{m}}. \tag{S2.6.4b}$$

Second, substituting the expressions for the reproductive value of old couples of type rm, $v_{2,\mathrm{rm}}$ (S2.6.3e), and the reproductive value of old couples of type mr, $v_{2,\mathrm{mr}}$ (S2.6.3f), into the equations for the reproductive value of young mutant couples (equations (S2.6.3c) and (S2.6.3d)), rearranging, and using the definition of productivities $\Pi_{\ell,i,k}$ (S1.6.1), we get

$$v_{1,\mathrm{mr}} = \Pi^\circ_{♀,\mathrm{m},\mathrm{mr}} v_{♀,\mathrm{m}} + \Pi^\circ_{♂,\mathrm{m},\mathrm{mr}} v_{♂,\mathrm{m}}, \tag{S2.6.5a}$$

$$v_{1,\mathrm{rm}} = \Pi^\circ_{♀,\mathrm{m},\mathrm{rm}} v_{♀,\mathrm{m}} + \Pi^\circ_{♂,\mathrm{m},\mathrm{rm}} v_{♂,\mathrm{m}}. \tag{S2.6.5b}$$

We can now use expressions (S2.6.4) and (S2.6.5) in order to derive two identities linking the reproductive values of various classes. We start by equating the right hand sides of the two expressions for $v_{1,\mathrm{mr}}$ above (equations (S2.6.4a) and (S2.6.5a)), and simplify to obtain

$$\left(\Pi_{♀,\mathrm{r,rr}} - \Pi^{\circ}_{♀,\mathrm{m,mr}}\right) v_{♀,\mathrm{m}} = \Pi^{\circ}_{♂,\mathrm{m,mr}} v_{♂,\mathrm{m}}$$

$$\left(1 - q_{♀,\mathrm{m,mr}}\right) \Pi_{♀,\mathrm{r,rr}} v_{♀,\mathrm{m}} = q_{♂,\mathrm{m,mr}} \Pi_{♂,\mathrm{r,rr}} v_{♂,\mathrm{m}}$$

$$\Pi_{♀,\mathrm{r,rr}} v_{♀,\mathrm{m}} = \Pi_{♂,\mathrm{r,rr}} v_{♂,\mathrm{m}} \tag{S2.6.6}$$

$$v_{1,\mathrm{mr}} = v_{1,\mathrm{rm}}, \tag{S2.6.7}$$

where the second line follows from substituting the expressions for neutral productivities (S2.5.10); the third line follows because identity (S1.2.1d) implies $1 - q_{♀,\mathrm{m,mr}} = q_{♂,\mathrm{m,mr}}$; and the last line follows from equation (S2.6.4). Equation (S2.6.6) links the reproductive values of female and male reproductives, and can be interpreted as stating that the reproductive value of a mutant reproductive of a given sex is proportional to the number of resident reproductives of the opposite sex and inversely proportional to the number of resident reproductives of the same sex (S2.2.7). In turn, equation (S2.6.7) states that a consequence of this is that the reproductive values of a mutant young couple is the same, whether the female in the couple is mutant (i.e., the female is mutant and the male is resident) or the male in the couple is mutant (i.e., the female is resident and the male is mutant). Although we derived identities (S2.6.6) and (S2.6.7) by equating the expressions for $v_{1,\mathrm{mr}}$ above (equations (S2.6.4a) and (S2.6.5a)) we could have alternatively derived them by equating the two expressions for $v_{1,\mathrm{rm}}$ (equations (S2.6.4b) and (S2.6.5b)) and simplifying in a similar way.

We can now proceed to obtain expressions for the reproductive values in terms of our variables and parameters. First, because of our choice of letting $v_{♂,\mathrm{m}} = 1$, isolating $v_{♀,\mathrm{m}}$ from equation (S2.6.6) leads to

$$v_{♂,\mathrm{m}} = 1, \tag{S2.6.8a}$$

$$v_{♀,\mathrm{m}} = \frac{\Pi_{♂,\mathrm{r,rr}}}{\Pi_{♀,\mathrm{r,rr}}}, \tag{S2.6.8b}$$

for the reproductive values of unmated mutants. Thus, the reproductive value of unmated mutant females equals the resident sex ratio (S2.2.7). Second, substituting (S2.6.8) into (S2.6.4) and simplifying, we obtain

$$v_{1,\mathrm{rm}} = v_{1,\mathrm{mr}} = \Pi_{♂,\mathrm{r,rr}}, \tag{S2.6.9}$$

for the reproductive value of young couples. Finally, substituting (S2.6.8) into equations (S2.6.3e) and (S2.6.3f), using the expressions for neutral reproductive rates (S2.5.9), and simplifying, we obtain

$$v_{2,\mathrm{rm}} = q_{♀,\mathrm{m,rm}} F_{2,♀,\mathrm{r,rr}} \frac{\Pi_{♂,\mathrm{r,rr}}}{\Pi_{♀,\mathrm{r,rr}}} + q_{♂,\mathrm{m,rm}} F_{2,♂,\mathrm{r,rr}}, \tag{S2.6.10a}$$

$$v_{2,\mathrm{mr}} = q_{♀,\mathrm{m,mr}} F_{2,♀,\mathrm{r,rr}} \frac{\Pi_{♂,\mathrm{r,rr}}}{\Pi_{♀,\mathrm{r,rr}}} + q_{♂,\mathrm{m,mr}} F_{2,♂,\mathrm{r,rr}}, \tag{S2.6.10b}$$

for the reproductive value of old couples.

As stated above, we will later show (in section 2.7) that the generic selection gradient (S2.4.4) can be simplified so that it only depends on two entries of $\mathbf{v}$, namely the reproductive values of unmated females and males, which in turn depend only on the resident sex ratio (equations (S2.6.8)). We will then use the simplified notation

$$v_{\ell} \equiv v_{\ell,\mathrm{m}} \tag{S2.6.11}$$

for $\ell \in \{♀, ♂\}$ to refer to the reproductive values of unmated individuals. From equations (S2.6.8), (S2.6.11), and (S2.2.7), we have

$$v_{♂} \equiv v_{♂,\mathrm{m}} = 1, \tag{S2.6.12a}$$

$$v_{♀} \equiv v_{♀,\mathrm{m}} = \frac{\Pi_{♂,\mathrm{r,rr}}}{\Pi_{♀,\mathrm{r,rr}}} = \frac{N^*_{♂,\mathrm{r}}}{N^*_{♀,\mathrm{r}}}, \tag{S2.6.12b}$$

which are respectively the neutral reproductive values of unmated (mutant) males and females. So, in our model, the modulating effect of reproductive value on selection is encapsulated by the sex ratio.

More explicitly, substituting the expression for resident sex-specific productivity (S1.6.18) into (S2.6.12), via (S1.6.16), the sex-specific reproductive values are given by

$$v_{♂} = 1, \tag{S2.6.13a}$$

$$v_{♀} = \frac{\sigma_{1,♂} f_1 (1 - p_{♂,\mathrm{r,rr}}) s_1 + \sigma_{2,♂} \Pi_2 (f_{2,\mathrm{rr}}, h_{\mathrm{rr}})}{\sigma_{1,♀} f_1 (1 - p_{♀,\mathrm{r,rr}}) s_1 + \sigma_{2,♀} \Pi_2 (f_{2,\mathrm{rr}}, h_{\mathrm{rr}})}. \tag{S2.6.13b}$$

Hence, the reproductive value of females and males is the same ($v_{♀} = v_{♂} = 1$) if both sexes help (G = B, so $p_{♂,\mathrm{r,rr}} = p_{♀,\mathrm{r,rr}}$) and the sex proportion is unbiased in both broods ($\sigma_{a,\ell} = 1/2$ for $a \in \{1,2\}$ and $\ell \in \{♀, ♂\}$), for both diploids and haplodiploids. In contrast, females have a higher reproductive value than males ($v_{♀} > v_{♂} = 1$) if females help more than males ($p_{♂,\mathrm{r,rr}} < p_{♀,\mathrm{r,rr}}$) and the sex proportion is unbiased in both broods ($\sigma_{a,\ell} = 1/2$ for $a \in \{1,2\}$ and $\ell \in \{♀, ♂\}$), for both diploids and haplodiploids (see also [12, 6]).

Still more explicitly, using Fig. S5 and equations (S1.4.1), (S1.3.5), and (S1.1.5), the reproductive value of females (S2.6.13b) for each model case is given by

$$v_{♀} = \begin{cases} \dfrac{\sigma_{1,♂} f_1 (1-p) s_1 + \sigma_{2,♂} \Pi_2 (f_2, f_1 p)}{\sigma_{1,♀} f_1 (1-p) s_1 + \sigma_{2,♀} \Pi_2 (f_2, f_1 p)} & \text{for } \mathrm{C} \in \{\mathrm{O}, \mathrm{M}\} \text{ and } \mathrm{G} = \mathrm{B} \\[2ex] \dfrac{\sigma_{1,♂} f_1 s_1 + \sigma_{2,♂} \Pi_2 (f_2, f_1 \sigma_1 p)}{\sigma_{1,♀} f_1 (1-p) s_1 + \sigma_{2,♀} \Pi_2 (f_2, f_1 \sigma_1 p)} & \text{for } \mathrm{C} \in \{\mathrm{O}, \mathrm{M}\} \text{ and } \mathrm{G} = \mathrm{F} \\[2ex] \dfrac{\sigma_{1,♂} f_1 (1-p(x,y)) s_1 + \sigma_{2,♂} \Pi_2 (f_2, f_1 p(x,y))}{\sigma_{1,♀} f_1 (1-p(x,y)) s_1 + \sigma_{2,♀} \Pi_2 (f_2, f_1 p(x,y))} & \text{for } \mathrm{C} = \mathrm{S} \text{ and } \mathrm{G} = \mathrm{B} \\[2ex] \dfrac{\sigma_{1,♂} f_1 s_1 + \sigma_{2,♂} \Pi_2 (f_2, f_1 \sigma_1 p(x,y))}{\sigma_{1,♀} f_1 (1-p(x,y)) s_1 + \sigma_{2,♀} \Pi_2 (f_2, f_1 \sigma_1 p(x,y))} & \text{for } \mathrm{C} = \mathrm{S} \text{ and } \mathrm{G} = \mathrm{F}. \end{cases}$$

**Stable distribution, u.** Let us now calculate the stable distribution (i.e., the right eigenvector)

$$\mathbf{u}^\intercal = \left(u_{♀,\mathrm{m}}, u_{♂,\mathrm{m}}, u_{1,\mathrm{rm}}, u_{1,\mathrm{mr}}, u_{2,\mathrm{rm}}, u_{2,\mathrm{mr}}\right). \tag{S2.6.14}$$

From the definition of a right eigenvector, and since the leading eigenvalue of $\mathbf{J}^\circ_{\mathrm{mut}}$ is equal to one, we have

$$\mathbf{J}^\circ_{\mathrm{mut}} \mathbf{u} = \mathbf{u}. \tag{S2.6.15}$$

Performing the matrix multiplication stated in (S2.6.15) with $\mathbf{J}^{\circ}_{\text{mut}}$ given by (S2.4.6), we obtain the following system of linear equations

$$u_{♀,\text{m}} = F^{\circ}_{1,♀,\text{m,rm}} u_{1,\text{rm}} + F^{\circ}_{2,♀,\text{m,rm}} u_{2,\text{rm}} + F^{\circ}_{1,♀,\text{m,mr}} u_{1,\text{mr}} + F^{\circ}_{2,♀,\text{m,mr}} u_{2,\text{mr}}, \tag{S2.6.16a}$$

$$u_{♂,\text{m}} = F^{\circ}_{1,♂,\text{m,rm}} u_{1,\text{rm}} + F^{\circ}_{2,♂,\text{m,rm}} u_{2,\text{rm}} + F^{\circ}_{1,♂,\text{m,mr}} u_{1,\text{mr}} + F^{\circ}_{2,♂,\text{m,mr}} u_{2,\text{mr}}, \tag{S2.6.16b}$$

$$u_{1,\text{rm}} = \frac{u_{♂,\text{m}}}{\Pi^{\circ}_{♂,\text{r,rr}}}, \tag{S2.6.16c}$$

$$u_{1,\text{mr}} = \frac{u_{♀,\text{m}}}{\Pi^{\circ}_{♀,\text{r,rr}}}, \tag{S2.6.16d}$$

$$u_{2,\text{rm}} = s^{\circ}_{M,\text{rm}} u_{1,\text{rm}}, \tag{S2.6.16e}$$

$$u_{2,\text{mr}} = s^{\circ}_{M,\text{mr}} u_{1,\text{mr}}. \tag{S2.6.16f}$$

We manipulate these equations in a similar way to what we did for the system describing the reproductive values of our model. First, we isolate $u_{♂,\text{m}}$ and $u_{♀,\text{m}}$ from, respectively, equations (S2.6.16c) and (S2.6.16c), and use (S2.5.11) to obtain

$$u_{♀,\text{m}} = \Pi_{♀,\text{r,rr}} u_{1,\text{mr}}. \tag{S2.6.17a}$$

$$u_{♂,\text{m}} = \Pi_{♂,\text{r,rr}} u_{1,\text{rm}}, \tag{S2.6.17b}$$

Second, we substitute (S2.6.16e) and (S2.6.16f) into (S2.6.16a) and (S2.6.16b), and use the definition of the productivities $\Pi_{\ell,i,k}$ (S1.6.1) to get

$$u_{♀,\text{m}} = \Pi^{\circ}_{♀,\text{m,rm}} u_{1,\text{rm}} + \Pi^{\circ}_{♀,\text{m,mr}} u_{1,\text{mr}}, \tag{S2.6.18a}$$

$$u_{♂,\text{m}} = \Pi^{\circ}_{♂,\text{m,rm}} u_{1,\text{rm}} + \Pi^{\circ}_{♂,\text{m,mr}} u_{1,\text{mr}}. \tag{S2.6.18b}$$

Finally, we use expressions (S2.6.17) and (S2.6.18) to derive an identity linking the stable proportions of young couples of types rm and mr. We start by equating the right hand sides of the two expressions for $u_{♀,\text{m}}$ above (equations (S2.6.17a) and (S2.6.18a)), and simplify to obtain

$$\begin{aligned}
\left(\Pi_{♀,\text{r,rr}} - \Pi^{\circ}_{♀,\text{m,mr}}\right) u_{1,\text{mr}} &= \Pi^{\circ}_{♀,\text{m,rm}} u_{1,\text{rm}} \\
\left(1 - q_{♀,\text{m,mr}}\right) \Pi_{♀,\text{r,rr}} u_{1,\text{mr}} &= q_{♀,\text{m,rm}} \Pi_{♀,\text{r,rr}} u_{1,\text{rm}} \\
q_{♂,\text{m,mr}} u_{1,\text{mr}} &= q_{♀,\text{m,rm}} u_{1,\text{rm}} \\
\frac{u_{1,\text{mr}}}{u_{1,\text{rm}}} &= \frac{q_{♀,\text{m,rm}}}{q_{♂,\text{m,mr}}}
\end{aligned} \tag{S2.6.19}$$

where the second line follows from substituting the expressions for neutral productivities (S2.5.10); the third line follows because identity (S1.2.1d) implies $1 - q_{♀,\text{m,mr}} = q_{♂,\text{m,mr}}$; and the last line rearranges, where the ratio of the transmission probabilities is the one given by (S1.2.2a).

As stated above, we will later (section 2.7) show that the selection gradient (S2.4.4) can be simplified so that it only depends on two entries of $\mathbf{u}$, namely the stable proportions of mutant young couples of either type, which in turn depend only on the transmission probabilities (equations (S2.6.20c) and (S2.6.20d)). Thus, it will be convenient to normalize the right eigenvector $\mathbf{u}$ in such a way that $u_{1,\text{rm}} + u_{1,\text{mr}} = 1$, so that $u_{1,k}$ refers to the proportion of mutant young couples that are of type $k$. Imposing this constraint, equations (S2.6.19),

(S2.6.17), (S2.6.16e), and (S2.6.16f) lead to

$$u_{♀,\mathrm{m}} = \frac{q_{♀,\mathrm{m,rm}}}{q_{♀,\mathrm{m,rm}} + q_{♂,\mathrm{m,mr}}} \Pi_{♀,\mathrm{r,rr}}, \tag{S2.6.20a}$$

$$u_{♂,\mathrm{m}} = \frac{q_{♂,\mathrm{m,mr}}}{q_{♀,\mathrm{m,rm}} + q_{♂,\mathrm{m,mr}}} \Pi_{♂,\mathrm{r,rr}}, \tag{S2.6.20b}$$

$$u_{1,\mathrm{rm}} = \frac{q_{♂,\mathrm{m,mr}}}{q_{♀,\mathrm{m,rm}} + q_{♂,\mathrm{m,mr}}}, \tag{S2.6.20c}$$

$$u_{1,\mathrm{mr}} = \frac{q_{♀,\mathrm{m,rm}}}{q_{♀,\mathrm{m,rm}} + q_{♂,\mathrm{m,mr}}}, \tag{S2.6.20d}$$

$$u_{2,\mathrm{rm}} = \frac{q_{♂,\mathrm{m,mr}}}{q_{♀,\mathrm{m,rm}} + q_{♂,\mathrm{m,mr}}} s_{M,\mathrm{rr}}, \tag{S2.6.20e}$$

$$u_{2,\mathrm{mr}} = \frac{q_{♀,\mathrm{m,rm}}}{q_{♀,\mathrm{m,rm}} + q_{♂,\mathrm{m,mr}}} s_{M,\mathrm{rr}}, \tag{S2.6.20f}$$

where we have also used the fact that $s^{\circ}_{M,\mathrm{rm}} = s^{\circ}_{M,\mathrm{mr}} = s_{M,\mathrm{rr}}$ (S2.5.6).

Since the simplified selection gradient will only depend $u_{1,\mathrm{rm}}$ and $u_{1,\mathrm{mr}}$, we will henceforth use the simplified notation

$$u_k \equiv u_{1,k} \tag{S2.6.21}$$

for $k \in \{\mathrm{rm}, \mathrm{mr}\}$, and term the vector

$$\tilde{\mathbf{u}}^{\mathsf{T}} = \left(u_{♀}, u_{♂}\right) \tag{S2.6.22}$$

the *stable sex distribution* of a neutral mutant allele among young parents, which in turn depends only on the transmission asymmetry. From equations (S2.6.20c), (S2.6.20d), (S2.6.21), and (S2.6.22), we have

$$u_{♀} \equiv u_{\mathrm{mr}} = \frac{q_{♀,\mathrm{m,rm}}}{q_{♀,\mathrm{m,rm}} + q_{♂,\mathrm{m,mr}}}, \tag{S2.6.23a}$$

$$u_{♂} \equiv u_{\mathrm{rm}} = \frac{q_{♂,\mathrm{m,mr}}}{q_{♀,\mathrm{m,rm}} + q_{♂,\mathrm{m,mr}}}, \tag{S2.6.23b}$$

as expressions for the neutral stable proportions of couples of type rm and mr. So, the modulating effect of the stable distribution on selection in our model is encapsulated by the transmission asymmetry.

**Link between the stable distribution, u, and "genetic reproductive values".** Because of our choice regarding the normalization of the leading eigenvector **u**, the stable proportions (S2.6.23) give a well-defined probability distribution. For diploids (P = D) and from Fig. (S4), the stable sex distribution is

$$\tilde{\mathbf{u}}^{\mathsf{T}} = \left(u_{♀}, u_{♂}\right) = \left(u_{1,\mathrm{mr}}, u_{1,\mathrm{rm}}\right) = (1/2, 1/2), \tag{S2.6.24}$$

while for haplodiploids (P = HD) and from Fig. (S4), it is

$$\tilde{\mathbf{u}}^{\mathsf{T}} = \left(u_{♀}, u_{♂}\right) = \left(u_{1,\mathrm{mr}}, u_{1,\mathrm{rm}}\right) = (2/3, 1/3). \tag{S2.6.25}$$

Hence, in a diploid population, a neutral mutation is asymptotically equally likely to be in a young mother or a young father, but in a haplodiploid population it is twice as likely to be in a young mother than in a young father. The asymmetry in the haplodiploid case is a consequence of the sex-related transmission asymmetry of such genetic system (see equation (S2.6.19)).

The entries of the stable sex distribution ($u_\ell$ for $\ell \in \{♀, ♂\}$; equations (S2.6.24) and (S2.6.25)) coincide with the "genetic reproductive values" or "sex-specific reproductive values" that often appear in the literature of social insects and social evolution ([13, 14, 15, 16]; see also pp. 39-41 of [17] and pp. 190-191 of [18]). Such genetic reproductive values are typically used to weigh sex-specific fitness effects so that allele frequency does not change without selection. They are interpreted as describing that, irrespectively of the sex ratio, in a haplodiploid population a male is worth half as much as a female in transmitting genes because he can pass on his genes only through daughters, while a female passes on her genes through both daughters and sons. Genetic reproductive values are often calculated as the normalized dominant left eigenvector of a right stochastic (rows sum to one) "gene flow" matrix ($A$ on p. 151 of [15]) or as the normalized dominant right eigenvector of a left stochastic (columns sum to one) matrix (**P** on p. 40 of [17]).

The stable sex distribution can also be obtained as follows. Let us define the *transmission matrix*

$$\mathbf{Q} = \begin{pmatrix} q_{♀,♀} & q_{♀,♂} \\ q_{♂,♀} & q_{♂,♂} \end{pmatrix} \equiv \begin{pmatrix} q_{♀,\mathrm{m,mr}} & q_{♀,\mathrm{m,rm}} \\ q_{♂,\mathrm{m,mr}} & q_{♂,\mathrm{m,rm}} \end{pmatrix} \tag{S2.6.26}$$

where $q_{\ell,\ell'}$ stands for the probability that a mutant parent of sex $\ell'$ transmits its mutant allele to an offspring of sex $\ell$ when the mutant allele is rare (and hence the second parent is of resident genotype). By (S1.2.1d), **Q** is left stochastic (i.e., its columns sum to one) and hence its dominant eigenvalue is one. Direct calculation shows that $\tilde{\mathbf{u}}^\intercal = (u_♀, u_♂) \equiv (u_{1,\mathrm{mr}}, u_{1,\mathrm{rm}})$ is a dominant right eigenvector of **Q**. Note also that since $\tilde{\mathbf{u}}$ is both a right eigenvector of **Q** and a probability distribution, we have that

$$\sum_{k\in\{\mathrm{rm,mr}\}} u_k q_{\ell,\mathrm{m},k} = \sum_{k\in\{♀,♂\}} u_k q_{\ell,k} = u_\ell \quad \forall \ell \in \{♀, ♂\}, \tag{S2.6.27}$$

that is, the neutral asymptotic probability that an individual of sex $\ell$ is a mutant is also equal to $u_\ell$. For diploids (P = D) and from Fig. (S4),

$$\mathbf{Q} = \begin{pmatrix} 1/2 & 1/2 \\ 1/2 & 1/2 \end{pmatrix}, \tag{S2.6.28}$$

for which equation (S2.6.24) is a dominant right eigenvector. For haplodiploids (P = HD) and from Fig. (S4),

$$\mathbf{Q} = \begin{pmatrix} 1/2 & 1 \\ 1/2 & 0 \end{pmatrix}, \tag{S2.6.29}$$

for which equation (S2.6.25) is a dominant right eigenvector.

Thus, for the specific values of the transmission probabilities under diploidy or haploidiploidy, our transmission matrix **Q** coincides with the matrix **P** of [17] (p. 40) and with the transpose of the gene-flow matrix $A$ of [15] (p. 151). In any case, the $(2/3, 1/3)$ weights can be interpreted as the stable sex distribution.

#### 2.7 Selection gradient (generic, simplified form)

Having calculated the left eigenvector **v** and right eigenvector **u** associated to the leading eigenvalue of $\mathbf{J}^\circ_{\mathrm{mut}}$, we can proceed to simplify the selection gradient $\mathcal{S}_\zeta(\mathbf{z})$ of a generic trait $\zeta$ (where $\zeta \in \{p, z\}$ for offspring and maternal control, whereas $\zeta \in \{x, y, z\}$ for shared control).

Our starting point is the generic expression of the selection gradient of $\zeta$ given by (S2.4.4). Taking the partial derivatives of the elements of the mutant submatrix $\mathbf{J}_{\text{mut}}$ (S2.3.4) with respect to the mutant trait value $\zeta_{\text{m}}$, and since the resident productivities $\Pi_{♀,\text{r},\text{rr}}$ and $\Pi_{♂,\text{r},\text{rr}}$ appearing in the first two columns of $\mathbf{J}_{\text{mut}}$ are independent of $\zeta_{\text{m}}$, we have

$$\mathcal{S}_{\zeta}(\mathbf{z}) = \frac{1}{\mathbf{v}^{\intercal}\mathbf{u}} \sum_{k\in\{\text{rm},\text{mr}\}} \left[ v_{2,k} \left.\frac{\partial s_{M,k}}{\partial \zeta_{\text{m}}}\right|_{\mathbf{z}_{\text{m}}=\mathbf{z}} u_{1,k} + \sum_{\ell\in\{♀,♂\}} v_{\ell,\text{m}} \sum_{a\in\{1,2\}} \left.\frac{\partial F_{a,\ell,\text{m},k}}{\partial \zeta_{\text{m}}}\right|_{\mathbf{z}_{\text{m}}=\mathbf{z}} u_{a,k} \right]. \tag{S2.7.1}$$

From equations (S2.6.16e) and (S2.6.16f), $u_{2,k} = s^{\circ}_{M,k} u_{1,k}$ holds for $k \in \{\text{rm},\text{mr}\}$. Substituting this expression into (S2.7.1) and collecting the $u_{1,k}$'s yields

$$\mathcal{S}_{\zeta}(\mathbf{z}) = \frac{1}{\mathbf{v}^{\intercal}\mathbf{u}} \sum_{k\in\{\text{rm},\text{mr}\}} \left\{ v_{2,k} \left.\frac{\partial s_{M,k}}{\partial \zeta_{\text{m}}}\right|_{\mathbf{z}_{\text{m}}=\mathbf{z}} + \sum_{\ell\in\{♀,♂\}} v_{\ell,\text{m}} \left[ \left.\frac{\partial F_{1,\ell,\text{m},k}}{\partial \zeta_{\text{m}}}\right|_{\mathbf{z}_{\text{m}}=\mathbf{z}} + \left.\frac{\partial F_{2,\ell,\text{m},k}}{\partial \zeta_{\text{m}}}\right|_{\mathbf{z}_{\text{m}}=\mathbf{z}} s^{\circ}_{M,k} \right] \right\} u_{1,k}. \tag{S2.7.2}$$

Also, from equations (S2.6.3e) and (S2.6.3f), $v_{2,k} = F^{\circ}_{2,♀,\text{m},k} v_{♀,\text{m}} + F^{\circ}_{2,♂,\text{m},k} v_{♂,\text{m}}$ hold for $k \in \{\text{rm},\text{mr}\}$. Substituting this expression into equation (S2.7.2) and collecting the $v_{\ell,\text{m}}$'s yields

$$\mathcal{S}_{\zeta}(\mathbf{z}) = \frac{1}{\mathbf{v}^{\intercal}\mathbf{u}} \sum_{\ell\in\{♀,♂\}} v_{\ell,\text{m}} \sum_{k\in\{\text{rm},\text{mr}\}} \left[ \left.\frac{\partial F_{1,\ell,\text{m},k}}{\partial \zeta_{\text{m}}}\right|_{\mathbf{z}_{\text{m}}=\mathbf{z}} + F^{\circ}_{2,\ell,\text{m},k} \left.\frac{\partial s_{M,k}}{\partial \zeta_{\text{m}}}\right|_{\mathbf{z}_{\text{m}}=\mathbf{z}} + \left.\frac{\partial F_{2,\ell,\text{m},k}}{\partial \zeta_{\text{m}}}\right|_{\mathbf{z}_{\text{m}}=\mathbf{z}} s^{\circ}_{M,k} \right] u_{1,k}. \tag{S2.7.3}$$

Finally, from the definition of productivities $\Pi_{\ell,i,k}$ (S1.6.1), by using the simplified notation for sex-specific reproductive values (S2.6.11) and stable sex distribution (S2.6.21), and by the product rule of derivatives, equation (S2.7.3) can be more succinctly written as

$$\mathcal{S}_{\zeta}(\mathbf{z}) = \frac{1}{\mathbf{v}^{\intercal}\mathbf{u}} \sum_{\ell\in\{♀,♂\}} \sum_{k\in\{\text{rm},\text{mr}\}} v_{\ell} \left.\frac{\partial \Pi_{\ell,\text{m},k}}{\partial \zeta_{\text{m}}}\right|_{\mathbf{z}_{\text{m}}=\mathbf{z}} u_k. \tag{S2.7.4}$$

Since $\mathbf{v}^{\intercal}\mathbf{u} > 0$ holds, the selection gradient of $\zeta$ is positive (i.e., $\zeta$ is favored by selection) if and only if

$$\sum_{k\in\{\text{rm},\text{mr}\}} u_k \sum_{\ell\in\{♀,♂\}} \left.\frac{\partial \Pi_{\ell,\text{m},k}}{\partial \zeta_{\text{m}}}\right|_{\mathbf{z}_{\text{m}}=\mathbf{z}} v_{\ell} > 0. \tag{S2.7.5}$$

This condition has an intuitive interpretation: a trait $\zeta$ is favored by selection if and only if the effect of a mutation in the trait on the mutant productivity of a couple, averaged over the stable sex distribution of parents and weighted by the sex-specific reproductive values of offspring, is positive.

In addition to providing a natural interpretation for the action and direction of natural selection, equation (S2.7.4) is convenient for our subsequent analysis because all important terms (those appearing on the left-hand side of (S2.7.5)) are written in terms of (marginal) productivities, sex-specific reproductive values, and the stable sex distribution, thus abstracting away the additional complication of having age classes for couples. Note also that the sex-specific reproductive values (S2.6.13) depend in general on the sex proportions of the two broods, on whether both sexes help or only females do, and on the evolving traits, but not on the transmission probabilities and hence on the genetic system. In contrast, the stable sex distribution (S2.6.23) depends exclusively on the transmission probabilities and hence on the genetic system but not on any other feature of the model.

#### 2.8 Selection gradient of traits affecting helping

##### 2.8.1 Derivation of the general expression

**General expression.** Consider a trait $\zeta$ affecting the probability of helping, that is, either $\zeta = p$ for model cases of offspring and maternal control, or $\zeta \in \{x, y\}$ for model cases of shared control. In this section, we obtain expressions for the selection gradient of these traits by explicitly calculating the derivatives appearing in equation (S2.7.4).

Evaluating the productivity $\Pi_{\ell,i,k}$ (S1.6.17) at $i = \text{m}$, and differentiating the resulting expression with respect to $\zeta_\text{m}$ using the chain rule, we obtain

$$\begin{aligned}
\left.\frac{\partial \Pi_{\ell,\text{m},k}}{\partial \zeta_\text{m}}\right|_{\mathbf{z}_\text{m}=\mathbf{z}} &= \left.\left(\frac{\partial}{\partial \zeta_\text{m}} q_{\ell,\text{m},k}\left[f_1\sigma_{1,\ell}(1-p_{\ell,\text{m},k})s_1+\sigma_{2,\ell}\Pi_{2,k}\right]\right)\right|_{\mathbf{z}_\text{m}=\mathbf{z}}\\
&= q_{\ell,\text{m},k}\left(-f_1\sigma_{1,\ell}\left.\frac{\partial p_{\ell,\text{m},k}}{\partial \zeta_\text{m}}\right|_{\mathbf{z}_\text{m}=\mathbf{z}} s_1+\sigma_{2,\ell}\left.\frac{\partial \Pi_{2,k}}{\partial h_k}\right|_{\mathbf{z}_\text{m}=\mathbf{z}}\times\left.\frac{\partial h_k}{\partial \zeta_\text{m}}\right|_{\mathbf{z}_\text{m}=\mathbf{z}}\right)\\
&= q_{\ell,\text{m},k}\left(-f_1\sigma_{1,\ell}\left.\frac{\partial p_{\ell,\text{m},k}}{\partial \zeta_\text{m}}\right|_{\mathbf{z}_\text{m}=\mathbf{z}} s_1+\sigma_{2,\ell}\frac{\partial \Pi_2}{\partial h}(f_2,h)\times f_1\sum_{\ell'\in\{♀,♂\}}\sigma_{1,\ell'}\sum_{i'\in\{\text{r},\text{m}\}}q_{\ell',i',k}\left.\frac{\partial p_{\ell',i',k}}{\partial \zeta_\text{m}}\right|_{\mathbf{z}_\text{m}=\mathbf{z}}\right)\\
&= f_1 q_{\ell,\text{m},k}\left(-\sigma_{1,\ell}\left.\frac{\partial p_{\ell,\text{m},k}}{\partial \zeta_\text{m}}\right|_{\mathbf{z}_\text{m}=\mathbf{z}} s_1+\sigma_{2,\ell}\frac{\partial \Pi_2}{\partial h}(f_2,h)\times\sum_{\ell'\in\{♀,♂\}}\sigma_{1,\ell'}\sum_{i'\in\{\text{r},\text{m}\}}q_{\ell',i',k}\left.\frac{\partial p_{\ell',i',k}}{\partial \zeta_\text{m}}\right|_{\mathbf{z}_\text{m}=\mathbf{z}}\right),
\end{aligned} \tag{S2.8.1}$$

where we have used the expression for $h_k$ given in (S1.3.3b), and the fact that the functional form for late productivity $\Pi_{2,k}$ is the same for all types $k$ (equation (S1.6.16)), which together with our notational conventions allows us to write

$$\left.\frac{\partial \Pi_{2,k}}{\partial h_k}\right|_{\mathbf{z}_\text{m}=\mathbf{z}} = \frac{\partial \Pi_2}{\partial h}(f_{2,k}^\circ, h_k^\circ) = \frac{\partial \Pi_2}{\partial h}(f_2, h).$$

Substituting (S2.8.1) into (S2.7.4) and rearranging, we obtain

$$\mathcal{S}_\zeta(\mathbf{z}) = \frac{1}{\mathbf{v}^\intercal\mathbf{u}} f_1\left(-\iota s_1+\kappa\frac{\partial \Pi_2}{\partial h}(f_2,h)\right), \tag{S2.8.2}$$

where

$$\iota = \sum_{\ell\in\{♀,♂\}}\sigma_{1,\ell}\sum_{k\in\{\text{rm},\text{mr}\}}u_k q_{\ell,\text{m},k}\left.\frac{\partial p_{\ell,\text{m},k}}{\partial \zeta_\text{m}}\right|_{\mathbf{z}_\text{m}=\mathbf{z}} v_\ell, \tag{S2.8.3a}$$

$$\kappa = \sum_{\ell\in\{♀,♂\}}\sigma_{1,\ell}\sum_{\ell'\in\{♀,♂\}}\sigma_{2,\ell'}\sum_{k\in\{\text{rm},\text{mr}\}}u_k\sum_{i'\in\{\text{r},\text{m}\}}q_{\ell,i',k}\left.\frac{\partial p_{\ell,i',k}}{\partial \zeta_\text{m}}\right|_{\mathbf{z}_\text{m}=\mathbf{z}} q_{\ell',\text{m},k}v_{\ell'}. \tag{S2.8.3b}$$

We call coefficients $\iota$ and $\kappa$ the *structure coefficients*. Since $\boldsymbol{\sigma}_1^\intercal$ and $\tilde{\mathbf{u}}^\intercal$ are probability distributions, (S2.8.3a) shows that $\iota$ is the effect of a mutation on helping evaluated at neutrality ($\partial p_{\ell,\text{m},k}/\partial\zeta_\text{m}|_{\mathbf{z}_\text{m}=\mathbf{z}}$), averaged over the sexes of parents ($u_k$) and of potentially helping offspring ($\sigma_{1,\ell}$), and weighted by the probability that a sex-$\ell$ potentially helping offspring has the mutation ($q_{\ell,\text{m},k}$) and by such offspring's reproductive value ($v_\ell$). Thus, $\iota$ is a weighted average of a helping mutation's phenotypic effect, with the weight given by the probability that candidate helpers have the mutation and by their reproductive value. Similarly, (S2.8.3b) shows that $\kappa$ is the effect of a mutation on helping evaluated at neutrality ($\partial p_{\ell,i',k}/\partial\zeta_\text{m}|_{\mathbf{z}_\text{m}=\mathbf{z}}$), averaged over the sexes of parents ($u_k$), of potentially helping offspring ($\sigma_{1,\ell}$), and of potentially helped offspring ($\sigma_{2,\ell}$), and over the probability that a potentially helping offspring has the mutation ($q_{\ell,i',k}$), and weighted by the probability that a sex-$\ell'$ potentially helped offspring has the mutation ($q_{\ell',\text{m},k}$) and by such offspring's reproductive value ($v_{\ell'}$). Thus, $\kappa$ is a weighted average of a helping mutation's phenotypic effect, with the weight given by the probability that candidate recipients of help have the mutation and by their reproductive value.

We now provide an interpretation for the remaining terms in large parentheses in equation (S2.8.2).

**Marginal cost and benefit of helping.** The factors $-s_1$ and $\partial\Pi_2(f_2,h)/\partial h$ appearing in (S2.8.2) have immediate interpretations in terms of marginal effects of the expected number of helpers on a couple's productivity. First, $\partial\Pi_2(f_2,h)/\partial h$ is the marginal effect of the expected number of helpers on the late productivity of a couple. Second, $s_1$ is the marginal effect of the expected number of helpers on the early productivity of a couple (as it can be verified from equation (S1.6.9)). To underline the fact that the marginal effect on early productivity is always negative (because $s_1 > 0$), while the marginal effect on late productivity is always positive (since, given our assumptions on the vital rates given in section 1.4, $\Pi_2$ is increasing in $h$) and for subsequent use, we introduce the following definitions and notation. We define

$$C = -\frac{\mathrm{d}\Pi_1(h)}{\mathrm{d}h} = s_1, \tag{S2.8.4}$$

as the *(marginal) cost of helping*, and

$$B = \frac{\partial\Pi_2}{\partial h}(f_2(z),h). \tag{S2.8.5}$$

as the *(marginal) benefit of helping* or the *marginal late productivity of helping*.

Note that the marginal cost of helping $C$ equals the constant $s_1$ for all the model cases we consider. In contrast, the marginal benefit of helping is a function of the evolving traits and of the neutral expected number of helpers $h$ and hence takes a different form for each model case, depending on who controls the helping probability and on the sex of the helpers. To make this dependence explicit, hereafter we write $B^{\mathrm{C,G}}$ for the benefit of helping when help control is of type C (where $\mathrm{C} \in \{\mathrm{O,M,S}\}$) and when the helpers' sex is G (where $\mathrm{G} \in \{\mathrm{B,F}\}$). Explicitly, using equations (S1.3.5) and (S1.1.5), the marginal benefit of helping (S2.8.5) for each model case is given by

$$B^{\mathrm{C,G}} = \begin{cases} \dfrac{\partial\Pi_2}{\partial h}(f_2, f_1 p) & \text{for } \mathrm{C} \in \{\mathrm{O,M}\} \text{ and } \mathrm{G} = \mathrm{B} \\[2ex] \dfrac{\partial\Pi_2}{\partial h}(f_2, f_1\sigma_1 p) & \text{for } \mathrm{C} \in \{\mathrm{O,M}\} \text{ and } \mathrm{G} = \mathrm{F} \\[2ex] \dfrac{\partial\Pi_2}{\partial h}(f_2, f_1 p(x,y)) & \text{for } \mathrm{C} = \mathrm{S} \text{ and } \mathrm{G} = \mathrm{B} \\[2ex] \dfrac{\partial\Pi_2}{\partial h}(f_2, f_1\sigma_1 p(x,y)) & \text{for } \mathrm{C} = \mathrm{S} \text{ and } \mathrm{G} = \mathrm{F}. \end{cases} \tag{S2.8.6}$$

**Critical benefit-cost ratio.** With the above definitions of helping cost and benefit, equation (S2.8.2) becomes

$$\mathcal{S}_\zeta(\mathbf{z}) = \frac{1}{\mathbf{v}^\intercal\mathbf{u}} f_1 \left(-\iota C + \kappa B\right). \tag{S2.8.7}$$

Since $f_1/\mathbf{v}^\intercal\mathbf{u} > 0$, the selection gradient of $\zeta$ is positive, and $\zeta$ is under positive directional selection when

$$-\iota C + \kappa B > 0, \tag{S2.8.8}$$

or equivalently,

$$\frac{B}{C} > \left(\frac{B}{C}\right)^* \quad \text{if } \kappa > 0, \text{ or} \tag{S2.8.9a}$$

$$\frac{B}{C} < \left(\frac{B}{C}\right)^* \quad \text{if } \kappa < 0 \tag{S2.8.9b}$$

where the *critical benefit-cost ratio* $(B/C)^*$ equals the ratio of the structure coefficients $\iota$ and $\kappa$ (S2.8.3):

$$\left(\frac{B}{C}\right)^* = \frac{\iota}{\kappa}. \tag{S2.8.10}$$

The case $\kappa > 0$ holds when the trait is the helping probability or maternal influence ($\zeta \in \{p, x\}$) because in that case $\partial p/\partial\zeta > 0$. In turn, the case $\kappa < 0$ holds when the trait is offspring resistance ($\zeta = y$) because in that case $\partial p/\partial\zeta < 0$.

As with the marginal benefit of helping $B$, the structure coefficients $\iota$ and $\kappa$ depend on who controls the helping probability (C) and the helpers' sex (G). To make this dependence explicit, and similarly to how we did for the benefit of helping, hereafter we write $\mathcal{S}_\zeta^{\mathrm{C,G}}$, $\iota_\zeta^{\mathrm{C,G}}$, $\kappa_\zeta^{\mathrm{C,G}}$, and $(B/C)_\zeta^{*\mathrm{C,G}}$ for the selection gradient, the structure coefficients, and the critical benefit-cost ratio for trait $\zeta$, under help control C and helpers' sex G.

##### 2.8.2 Derivation for each model case

We now obtain explicit expressions for the structure coefficients and the critical benefit-cost ratios under the model cases we consider.

**Offspring control, both sexes help (O-B).** For offspring control, $\zeta = p$, and hence $\zeta_\mathrm{m} = p_\mathrm{m}$. Then, in the case of offspring control, and if both sexes help (see Fig. S5)

$$\left.\frac{\partial p_{\ell,i,k}}{\partial p_\mathrm{m}}\right|_{\mathbf{z}_\mathrm{m}=\mathbf{z}} = [i = \mathrm{m}], \quad \forall k \in \{\mathrm{rm}, \mathrm{mr}\} \text{ and } \forall \ell \in \{♀, ♂\}, \tag{S2.8.11}$$

where [ ] is the Iverson bracket, such that

$$[P] = \begin{cases} 1 & \text{if } P \text{ is true} \\ 0 & \text{otherwise.} \end{cases} \tag{S2.8.12}$$

Substituting (S2.8.11) into (S2.8.3) and simplifying using equation (S2.6.27) yields:

$$\iota_p^{\mathrm{O,B}} = \sum_{\ell \in \{♀,♂\}} \sigma_{1,\ell} u_\ell v_\ell, \tag{S2.8.13a}$$

$$\kappa_p^{\mathrm{O,B}} = \sum_{\ell \in \{♀,♂\}} \sigma_{1,\ell} \sum_{\ell' \in \{♀,♂\}} \sigma_{2,\ell'} \sum_{k \in \{\mathrm{rm},\mathrm{mr}\}} u_k q_{\ell,\mathrm{m},k} q_{\ell',\mathrm{m},k} v_{\ell'}. \tag{S2.8.13b}$$

We will provide an interpretation of $\iota_\zeta^{\mathrm{C,G}}$ and $\kappa_\zeta^{\mathrm{C,G}}$ later (section 3.2.4), which applies to all the cases we consider and which recovers an inclusive fitness interpretation.

The critical benefit-cost ratio is then given by

$$\left(\frac{B}{C}\right)_p^{*\mathrm{O,B}} = \frac{\sum_{\ell \in \{♀,♂\}} \sigma_{1,\ell} u_\ell v_\ell}{\sum_{\ell \in \{♀,♂\}} \sigma_{1,\ell} \sum_{\ell' \in \{♀,♂\}} \sigma_{2,\ell'} \sum_{k \in \{\mathrm{rm},\mathrm{mr}\}} u_k q_{\ell,\mathrm{m},k} q_{\ell',\mathrm{m},k} v_{\ell'}}. \tag{S2.8.14}$$

**Offspring control, only females help (O-F).** For offspring control, but now if only females help, we have (see Fig. S5)

$$\left.\frac{\partial p_{\ell,i,k}}{\partial p_\mathrm{m}}\right|_{\mathbf{z}_\mathrm{m}=\mathbf{z}} = [\ell = ♀ \text{ and } i = \mathrm{m}], \quad \forall k \in \{\mathrm{rm}, \mathrm{mr}\}. \tag{S2.8.15}$$

Substituting this expression into equation (S2.8.3) and simplifying using equation (S2.6.27) yields:

$$\iota_p^{\mathrm{O,F}} = \sigma_{1,♀} u_♀ v_♀, \tag{S2.8.16a}$$

$$\kappa_p^{\mathrm{O,F}} = \sigma_{1,♀} \sum_{\ell' \in \{♀,♂\}} \sigma_{2,\ell'} \sum_{k \in \{\mathrm{rm,mr}\}} u_k q_{♀,\mathrm{m},k} q_{\ell',\mathrm{m},k} v_{\ell'}. \tag{S2.8.16b}$$

The critical benefit-cost ratio thus reduces to

$$\left(\frac{B}{C}\right)_p^{*\mathrm{O,F}} = \frac{u_♀ v_♀}{\sum_{\ell' \in \{♀,♂\}} \sigma_{2,\ell'} \sum_{k \in \{\mathrm{rm,mr}\}} u_k q_{♀,\mathrm{m},k} q_{\ell',\mathrm{m},k} v_{\ell'}}. \tag{S2.8.17}$$

**Maternal control, both sexes help (M-B).** For maternal control with both sexes helping, we have (see Fig. S5)

$$\left.\frac{\partial p_{\ell,i,k}}{\partial p_{\mathrm{m}}}\right|_{\mathbf{z}_{\mathrm{m}}=\mathbf{z}} = [k = \mathrm{mr}] \ \forall \ell \in \{♀,♂\} \text{ and } \forall i \in \{\mathrm{r,m}\}. \tag{S2.8.18}$$

Substituting this expression into (S2.8.3) yields:

$$\iota_p^{\mathrm{M,B}} = u_{\mathrm{mr}} \sum_{\ell \in \{♀,♂\}} \sigma_{1,\ell} q_{\ell,\mathrm{m,mr}} v_\ell, \tag{S2.8.19a}$$

$$\begin{aligned}
\kappa_p^{\mathrm{M,B}} &= u_{\mathrm{mr}} \sum_{\ell \in \{♀,♂\}} \sigma_{1,\ell} \sum_{\ell' \in \{♀,♂\}} \sigma_{2,\ell'} \sum_{i' \in \{\mathrm{r,m}\}} q_{\ell,i',\mathrm{mr}} q_{\ell',\mathrm{m,mr}} v_{\ell'} \\
&= u_{\mathrm{mr}} \sum_{\ell \in \{♀,♂\}} \sigma_{1,\ell} \sum_{\ell' \in \{♀,♂\}} \sigma_{2,\ell'} q_{\ell',\mathrm{m,mr}} v_{\ell'} \\
&= u_{\mathrm{mr}} \sum_{\ell' \in \{♀,♂\}} \sigma_{2,\ell'} q_{\ell',\mathrm{m,mr}} v_{\ell'},
\end{aligned} \tag{S2.8.19b}$$

where we have used identities (S1.2.1c) and (S1.1.3).

The critical benefit-cost ratio is then

$$\left(\frac{B}{C}\right)_p^{*\mathrm{M,B}} = \frac{\sum_{\ell \in \{♀,♂\}} \sigma_{1,\ell} q_{\ell,\mathrm{m,mr}} v_\ell}{\sum_{\ell' \in \{♀,♂\}} \sigma_{2,\ell'} q_{\ell',\mathrm{m,mr}} v_{\ell'}}. \tag{S2.8.20}$$

**Maternal control, only females help (M-F).** For maternal control of the helping trait and if only females help, we have (see Fig. S5)

$$\left.\frac{\partial p_{\ell,i,k}}{\partial p_{\mathrm{m}}}\right|_{\mathbf{z}_{\mathrm{m}}=\mathbf{z}} = [k = \mathrm{mr} \text{ and } \ell = ♀] \ \ \forall i \in \{\mathrm{r,m}\}. \tag{S2.8.21}$$

Following the same steps as in the previous case (M-B), we obtain

$$\iota_p^{\mathrm{M,F}} = \sigma_{1,♀} u_{\mathrm{mr}} q_{♀,\mathrm{m,mr}} v_♀, \tag{S2.8.22a}$$

$$\kappa_p^{\mathrm{M,F}} = \sigma_{1,♀} u_{\mathrm{mr}} \sum_{\ell' \in \{♀,♂\}} \sigma_{2,\ell'} q_{\ell',\mathrm{m,mr}} v_{\ell'}, \tag{S2.8.22b}$$

with the critical benefit-cost ratio simplifying to

$$\left(\frac{B}{C}\right)_p^{*\mathrm{M,F}} = \frac{q_{♀,\mathrm{m,mr}} v_♀}{\sum_{\ell' \in \{♀,♂\}} \sigma_{2,\ell'} q_{\ell',\mathrm{m,mr}} v_{\ell'}}. \tag{S2.8.23}$$

**Shared control, both sexes help (S-B).** Consider now shared control, so that $\zeta \in \{x, y\}$ where $x$ is maternal influence and $y$ is offspring resistance.

Let us first calculate the structure coefficients and the critical benefit-cost ratio for maternal influence $x$. If both sexes help, then (see Fig. S5):

$$\left.\frac{\partial p_{\ell,i,k}}{\partial x_{\mathrm{m}}}\right|_{\mathbf{z}_{\mathrm{m}}=\mathbf{z}} = \frac{\partial p}{\partial x}(x,y)[k=\mathrm{mr}] \;\; \forall \ell \in \{♀,♂\} \text{ and } \forall i \in \{\mathrm{r},\mathrm{m}\}. \tag{S2.8.24}$$

Substituting this expression into equation (S2.8.3) and simplifying following the same steps as when calculating the coefficients for the case M-B yields:

$$\iota_x^{\mathrm{S,B}} = \frac{\partial p}{\partial x}(x,y)\iota_p^{\mathrm{M,B}}, \tag{S2.8.25a}$$

$$\kappa_x^{\mathrm{S,B}} = \frac{\partial p}{\partial x}(x,y)\kappa_p^{\mathrm{M,B}}, \tag{S2.8.25b}$$

where $\iota_p^{\mathrm{M,B}}$ and $\kappa_p^{\mathrm{M,B}}$ are as given by equation (S2.8.19). Hence, using (S2.8.7), it follows that

$$\mathcal{S}_x^{\mathrm{S,B}}(\mathbf{z}) = \frac{\partial p}{\partial x}(x,y)\mathcal{S}_p^{\mathrm{M,B}}(\mathbf{z}). \tag{S2.8.26}$$

Moreover, the critical benefit-cost ratio for maternal influence $x$ is

$$\left(\frac{B}{C}\right)_x^{*\mathrm{S,B}} = \left(\frac{B}{C}\right)_p^{*\mathrm{M,B}}, \tag{S2.8.27}$$

where $(B/C)_p^{*\mathrm{M,B}}$ is the critical benefit-cost ratio for $p$ for the case of maternal control and helpers from both sexes, as given by equation (S2.8.20).

Let us now calculate the structure coefficients and critical benefit-cost ratio for offspring resistance $y$. If both sexes help, then (see Fig. S5):

$$\left.\frac{\partial p_{\ell,i,k}}{\partial y_{\mathrm{m}}}\right|_{\mathbf{z}_{\mathrm{m}}=\mathbf{z}} = \frac{\partial p}{\partial y}(x,y)[i=\mathrm{m}] \;\; \forall k \in \{\mathrm{rm},\mathrm{mr}\} \text{ and } \forall \ell \in \{♀,♂\}. \tag{S2.8.28}$$

Substituting this expression into (S2.8.3) and simplifying following the same steps as when calculating the coefficients for the case O-B yields:

$$\iota_y^{\mathrm{S,B}} = \frac{\partial p}{\partial y}(x,y)\iota_p^{\mathrm{O,B}}, \tag{S2.8.29a}$$

$$\kappa_y^{\mathrm{S,B}} = \frac{\partial p}{\partial y}(x,y)\kappa_p^{\mathrm{O,B}}, \tag{S2.8.29b}$$

where $\iota_p^{\mathrm{O,B}}$ and $\kappa_p^{\mathrm{O,B}}$ are as given by equation (S2.8.13). Hence, using (S2.8.7), it follows that

$$\mathcal{S}_y^{\mathrm{S,B}}(\mathbf{z}) = \frac{\partial p}{\partial y}(x,y)\mathcal{S}_p^{\mathrm{O,B}}(\mathbf{z}). \tag{S2.8.30}$$

Moreover, the critical benefit-cost ratio for offspring resistance $y$ is

$$\left(\frac{B}{C}\right)_y^{*\mathrm{S,B}} = \left(\frac{B}{C}\right)_p^{*\mathrm{O,B}}, \tag{S2.8.31}$$

where $(B/C)_p^{*\mathrm{O,B}}$ is the critical benefit-cost ratio for $p$ for the case of offspring control and helpers from both sexes, as given by equation (S2.8.14).

**Shared control, only females help (S-F).** For maternal influence $x$, when only females help, we have (see Fig. S5)

$$\left.\frac{\partial p_{\ell,i,k}}{\partial x_{\mathrm{m}}}\right|_{\mathbf{z}_{\mathrm{m}}=\mathbf{z}} = \frac{\partial p}{\partial x}(x,y)[k=\mathrm{mr} \text{ and } \ell = ♀] \;\; \forall i \in \{\mathrm{r},\mathrm{m}\}. \tag{S2.8.32}$$

Substituting this expression into (S2.8.3) and simplifying following the same steps as when calculating the coefficients for the case M-F yields:

$$\iota_x^{\mathrm{S,F}} = \frac{\partial p}{\partial x}(x,y)\iota_p^{\mathrm{M,F}}, \tag{S2.8.33a}$$

$$\kappa_x^{\mathrm{S,F}} = \frac{\partial p}{\partial x}(x,y)\kappa_p^{\mathrm{M,F}}, \tag{S2.8.33b}$$

where $\iota_p^{\mathrm{M,F}}$ and $\kappa_p^{\mathrm{M,F}}$ are as given by equation (S2.8.22). Hence, using (S2.8.7), it follows that

$$\mathcal{S}_x^{\mathrm{S,F}}(\mathbf{z}) = \frac{\partial p}{\partial x}(x,y)\mathcal{S}_p^{\mathrm{M,F}}(\mathbf{z}). \tag{S2.8.34}$$

Moreover, we can write the critical benefit-cost ratio for maternal influence $x$ as

$$\left(\frac{B}{C}\right)_x^{*\mathrm{S,F}} = \left(\frac{B}{C}\right)_p^{*\mathrm{M,F}}, \tag{S2.8.35}$$

where $(B/C)_p^{*\mathrm{M,F}}$ is the critical benefit-cost ratio for $p$ for the case of maternal control when only females help, as given by equation (S2.8.23).

For offspring resistance $y$, we also have (see Fig. S5)

$$\left.\frac{\partial p_{\ell,i,k}}{\partial y_{\mathrm{m}}}\right|_{\mathbf{z}_{\mathrm{m}}=\mathbf{z}} = \frac{\partial p}{\partial y}(x,y)[\ell = ♀ \text{ and } i = \mathrm{m}] \quad \text{for all } k \in \{\mathrm{rm}, \mathrm{mr}\}. \tag{S2.8.36}$$

Substituting this expression into (S2.8.3) and simplifying following the same steps as when calculating the coefficients for the case O-F yields:

$$\iota_y^{\mathrm{S,F}} = \frac{\partial p}{\partial y}(x,y)\iota_p^{\mathrm{O,F}}, \tag{S2.8.37a}$$

$$\kappa_y^{\mathrm{S,F}} = \frac{\partial p}{\partial y}(x,y)\kappa_p^{\mathrm{O,F}}, \tag{S2.8.37b}$$

where $\iota_p^{\mathrm{O,F}}$ and $\kappa_p^{\mathrm{O,F}}$ are as given by equation (S2.8.16). Hence, using (S2.8.7), it follows that

$$\mathcal{S}_y^{\mathrm{S,F}}(\mathbf{z}) = \frac{\partial p}{\partial y}(x,y)\mathcal{S}_p^{\mathrm{O,F}}(\mathbf{z}). \tag{S2.8.38}$$

Moreover, we can write the critical benefit-cost ratio for offspring resistance $y$ as

$$\left(\frac{B}{C}\right)_y^{*\mathrm{S,F}} = \left(\frac{B}{C}\right)_p^{*\mathrm{O,F}}, \tag{S2.8.39}$$

where $(B/C)_p^{*\mathrm{O,F}}$ is the critical benefit-cost ratio for $p$ for the case of offspring control when only females help, as given by equation (S2.8.17).

##### 2.8.3 Summary

Summarizing, for model cases of offspring or maternal control of helping, the selection gradient of $p$ is

$$\mathcal{S}_p^{\mathrm{C,G}}(\mathbf{z}) = \frac{1}{\mathbf{v}^{\intercal}\mathbf{u}} f_1 \left(-\iota_p^{\mathrm{C,G}} C + \kappa_p^{\mathrm{C,G}} B\right), \tag{S2.8.40}$$

for $\mathrm{C} \in \{\mathrm{O}, \mathrm{M}\}$ and $\mathrm{G} \in \{\mathrm{B}, \mathrm{F}\}$. The structure coefficients $\iota_p^{\mathrm{C,G}}$ and $\kappa_p^{\mathrm{C,G}}$ are listed in Fig. S7A. This follows from (S2.8.7), (S2.8.13), (S2.8.16), (S2.8.19), and (S2.8.22).

For model cases of shared control, the selection gradients of $x$ and $y$ are

$$\mathcal{S}_x^{\mathrm{S,G}}(\mathbf{z}) = \frac{\partial p}{\partial x}(x,y)\mathcal{S}_p^{\mathrm{M,G}}(\mathbf{z}) \tag{S2.8.41a}$$

$$\mathcal{S}_y^{\mathrm{S,G}}(\mathbf{z}) = \frac{\partial p}{\partial y}(x,y)\mathcal{S}_p^{\mathrm{O,G}}(\mathbf{z}), \tag{S2.8.41b}$$

for $\mathrm{G} \in \{\mathrm{B}, \mathrm{F}\}$. This follows from (S2.8.26), (S2.8.30), (S2.8.34), and (S2.8.38).

#### 2.9 Selection gradient of reproductive effort

Finally, let us calculate the selection gradient of reproductive effort, $\zeta = z$, using equation (S2.7.4). Evaluating the expression for productivity $\Pi_{\ell,i,k}$ (S1.6.17) at $i = \mathrm{m}$, and differentiating the resulting expression with respect to $z_\mathrm{m}$ using the chain rule, we obtain

$$\begin{aligned}\left.\frac{\partial \Pi_{\ell,\mathrm{m},k}}{\partial z_\mathrm{m}}\right|_{\mathbf{z}_\mathrm{m}=\mathbf{z}} &= \left.\left(\frac{\partial}{\partial z_\mathrm{m}} q_{\ell,\mathrm{m},k}\left[\sigma_{1,\ell} f_1 (1-p_{\ell,\mathrm{m},k}) s_1 + \sigma_{2,\ell}\Pi_{2,k}\right]\right)\right|_{\mathbf{z}_\mathrm{m}=\mathbf{z}} \\ &= q_{\ell,\mathrm{m},k}\sigma_{2,\ell} \left.\frac{\partial \Pi_{2,k}}{\partial f_{2,k}}\right|_{\mathbf{z}_\mathrm{m}=\mathbf{z}} \left.\frac{\partial f_{2,k}}{\partial z_k}\right|_{\mathbf{z}_\mathrm{m}=\mathbf{z}} \left.\frac{\partial z_k}{\partial z_\mathrm{m}}\right|_{\mathbf{z}_\mathrm{m}=\mathbf{z}} \\ &= q_{\ell,\mathrm{m},k}\sigma_{2,\ell}\frac{\partial \Pi_2}{\partial f_2}(f_2,h)\frac{\mathrm{d}f_2}{\mathrm{d}z}(z)[k=\mathrm{mr}],\end{aligned} \tag{S2.9.1}$$

where the last equality follows from our assumptions on the functional form of the late productivity and late fertility of a couple (equations (S1.6.16) and (S1.4.1)) and from differentiating $z_k$ with respect to the mutant trait. Substituting (S2.9.1) into (S2.7.4) and simplifying, we obtain

$$\mathcal{S}_z(\mathbf{z}) = \frac{1}{\mathbf{v}^\intercal\mathbf{u}}\frac{\partial \Pi_2}{\partial f_2}(f_2,h)\frac{\mathrm{d}f_2}{\mathrm{d}z}(z)\, u_\mathrm{mr} \sum_{\ell'\in\{♀,♂\}} \sigma_{2,\ell'} q_{\ell',\mathrm{m},\mathrm{mr}} v_{\ell'}. \tag{S2.9.2}$$

The selection gradient of reproductive effort is a product of factors that can interpreted similarly as for the selection gradient of traits affecting helping. First, this selection gradient is proportional to the *marginal productivity of late fertility*

$$D = \frac{\partial \Pi_2}{\partial f_2}(f_2,h), \tag{S2.9.3}$$

that is, the marginal effect on a couple's lifetime productivity from a marginal increase in late fertility: since early productivity is independent from late fertility, the marginal effect on lifetime productivity from a marginal increase in late fertility equals the marginal effect on late productivity. As with the marginal benefit of helping (S2.8.6), the marginal productivity of late fertility depends on who controls the helping probability and on the sex of helpers via the neutral expected number of helpers, $h$. Thus, we follow a similar notational convention and write $D^{\mathrm{C,G}}$ for the marginal productivity of late fertility when help control is of type C and when the helpers' sex is G. Specifically, we have

$$D^{\mathrm{C,G}} = \begin{cases} \frac{\partial \Pi_2}{\partial f_2}(f_2(z), f_1 p) & \text{for C} \in \{\mathrm{O,M}\} \text{ and G} = \mathrm{B} \\ \frac{\partial \Pi_2}{\partial f_2}(f_2(z), f_1 \sigma_1 p) & \text{for C} \in \{\mathrm{O,M}\} \text{ and G} = \mathrm{F} \\ \frac{\partial \Pi_2}{\partial f_2}(f_2(z), f_1 p(x,y)) & \text{for C} = \mathrm{S} \text{ and G} = \mathrm{B} \\ \frac{\partial \Pi_2}{\partial f_2}(f_2(z), f_1 \sigma_1 p(x,y)) & \text{for C} = \mathrm{S} \text{ and G} = \mathrm{F} \end{cases}. \tag{S2.9.4}$$

Second, this selection gradient is proportional to the structure coefficient

$$\kappa_z^{\mathrm{C,G}} = u_\mathrm{mr} \sum_{\ell'\in\{♀,♂\}} \sigma_{2,\ell'} q_{\ell',\mathrm{m},\mathrm{mr}} v_{\ell'}. \tag{S2.9.5}$$

Although this structure coefficient has a similar form to the structure coefficient $\kappa_p^{\mathrm{M,B}}$ (S2.8.19b), reproductive values $v_{\ell'}$ depend on help control C and the helpers' sex G (S2.6.13), so $\kappa_z^{\mathrm{C,G}}$ and $\kappa_p^{\mathrm{M,B}}$ may be different.

With these two notational conventions, the selection gradient of reproductive effort for each model case is given by

$$\mathcal{S}_{z}^{\mathrm{C,G}}(\mathbf{z}) = \frac{1}{\mathbf{v}^{\intercal}\mathbf{u}} \frac{\mathrm{d}f_2}{\mathrm{d}z}(z)\kappa_{z}^{\mathrm{C,G}} D^{\mathrm{C,G}}. \tag{S2.9.6}$$

Since the factors $f_1/\mathbf{v}^{\intercal}\mathbf{u}$, $\kappa_{z}^{\mathrm{C,G}}$, and $\mathrm{d}f_2(z)/\mathrm{d}z$ are all strictly positive (e.g., (S1.4.3)), a necessary and sufficient condition for the selection gradient of reproductive effort $z$ to be positive, and for $z$ to be under positive directional selection is that the marginal productivity of fertility is positive, that is that

$$D^{\mathrm{C,G}} > 0 \tag{S2.9.7}$$

holds.

### 3 Inclusive fitness effects

Inclusive fitness describes selection in terms of how the phenotype of individual actors affects the personal fitness of recipients [19, 20, 21, 22]. In general, the inclusive fitness effect is the sum of the effects of a focal individual's phenotype on the fitness of recipients, where each effect is weighted by the relatedness of the actor to the recipient and by the reproductive value of the recipient.

In this section, we show that the sign of the selection gradient of all the traits in our model can be rewritten as the sign of an inclusive fitness effect. To do this, we proceed in six steps. First, we define *social classes*, *actors*, and *recipients* within a given nest, and introduce notation to refer to them (Social classes, actors, and recipients; section 3.1). Second, we define *reproductive worth*, which is an inclusive fitness measure of reproductive valuation of social partners, and show that the structure coefficients can be written in terms of such measure (Reproductive worth; section 3.2). Third, we define *relative reproductive worth*, which is a measure of relative reproductive valuation of social partners (Relative reproductive worth; section 3.3). Fourth, we define personal fitness functions to calculate inclusive fitness benefits and costs for a trait affecting helping (Individual cost and benefit of helping; section 3.4). Fifth, we write the selection gradient of a trait affecting helping in terms of the trait's *inclusive fitness effect* (Inclusive fitness effect for a trait affecting helping and Hamilton's rule; section 3.5). Finally, we define the inclusive fitness benefit for reproductive effort and write this trait's selection gradient in terms of the trait's inclusive fitness effect (Inclusive fitness effect for reproductive effort; section 3.6).

#### 3.1 Social classes, actors, and recipients

In the following, we introduce some notation to refer to the different sets of individuals (or *social classes*) of a "focal" nest in our model, and to distinguish between sets comprising actors and sets comprising recipients.

**Social classes.** We denote by $M$ the singleton whose only member is the mother of the nest; and by $O_{a\ell}$ the set of sex-$\ell$ offspring produced in brood $a$. The set of $a$-th brood offspring is denoted by $O_a$, where $O_a = O_{a♀} \cup O_{a♂}$. We illustrate these social classes in Fig. S6.

**Actors.** Actors are individuals that genetically control the trait $\zeta$ in consideration. In our model the set of actors $A$ is thus either (i) the mother's singleton $M$ (if helping is under maternal control, C = M; or if helping is under shared control, C = S, and the trait is maternal influence, $\zeta = x$), (ii) the set of first-brood offspring $O_1$ (if both sexes help, G = B, and either helping is under offspring control, C = O, or helping is under shared control, C = S, and the trait is resistance, $\zeta = y$), or (iii) the set of first-brood female offspring $O_{1♀}$ (if only females help, G = F, and either helping is under offspring control, C = O, or helping is under shared control, C = S, and the trait is resistance, $\zeta = y$). In short,

$$A = \begin{cases} M & \text{if C = M or (C = S and } \zeta = x) \\ O_1 & \text{if G = B and [C = O or (C = S and } \zeta = y)] \\ O_{1♀} & \text{if G = F and [C = O or (C = S and } \zeta = y)]. \end{cases} \tag{S3.1.1}$$

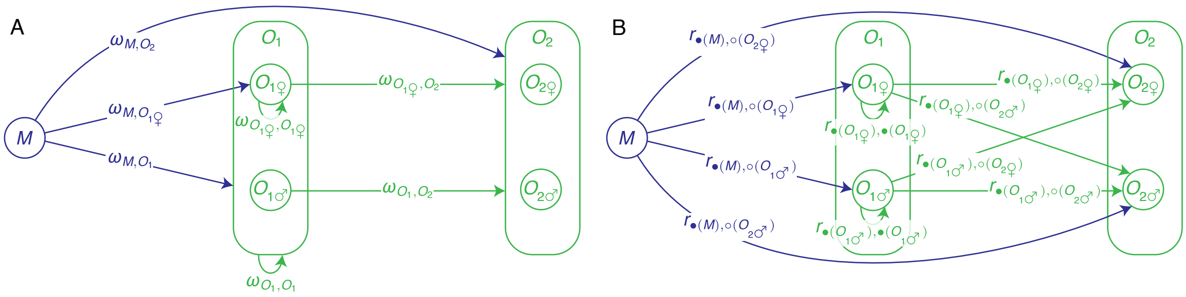

Figure S6: Social classes. Panels A and B show Venn diagrams illustrating the social classes in a given nest resulting in our model. (A) Reproductive worth coefficients resulting in our model, as shown in Fig. S7. (B) Relatedness coefficients involved in our model, as shown in section 3.2.1.

Moreover, we denote by $A_\ell$ the subset of sex-$\ell$ individuals in $A$, e.g., $A_♀ = O_{1♀}$ and $A_♂ = \emptyset$ if $A = O_{1♀}$, where $\emptyset$ is the empty set.

**Recipients.** Recipients are individuals whose fitness is affected by the trait. There are two types of recipients: individuals that can help (which we call *candidate helpers*), and individuals that can be helped (which we call *payees*). In our model the set of candidate helpers $H$ is thus either the set of first-brood offspring $O_1$ (if both sexes help, G = B), or (ii) the set of first-brood female offspring $O_{1♀}$ (if only females help, G = F). A candidate helper is not necessarily a helper and a payee is not necessarily helped (e.g., if $p = 0$). We will see that the set of payees $P$ is the set of second-brood offspring $P = O_2$ in all cases. Consequently, the set of recipients $R$ is either (i) the set of first-brood offspring $O_1$ (the candidate helpers if both sexes help, G = B), (ii) the set of first-brood female offspring $O_{1♀}$ (the candidate helpers if only females help, G = F), or (iii) the set of second-brood offspring $O_2$ (the payees). In short,

$$H = \begin{cases} O_1 & \text{if G = B} \\ O_{1♀} & \text{if G = F} \end{cases} \tag{S3.1.2a}$$

$$P = O_2 \tag{S3.1.2b}$$

$$R = \begin{cases} H & \text{for candidate helpers} \\ P & \text{for candidate recipients of help (payees).} \end{cases} \tag{S3.1.2c}$$

Moreover, we denote by $R_\ell$ the subset of sex-$\ell$ individuals in $R$.

#### 3.2 Reproductive worth

**Sampling experiment.** Consider a neutral ($\mathbf{z}_\mathrm{m} = \mathbf{z}$) rare mutant subpopulation introduced at a resident equilibrium. As ecological time $t$ advances, this mutant subpopulation asymptotically reaches a stable distribution proportional to $\mathbf{u}$ (S2.6.14); since the mutation is neutral, the mutation's frequency remains constant. Now consider sampling uniformly at random one young neutral mutant nest at ecological time $t \to \infty$. Having sampled a nest, we draw an individual actor uniformly at random from the subset $A_\ell$ of sex-$\ell$ actors in the

| A<br>$-\iota c+\kappa b>0$<br>Structure coefficients | B<br>Substituting relatedness | C<br>Substituting $\phi_\ell$ and $\sigma_\ell$ | D<br>Substituting reproductive worth |
|---|---|---|---|
| Offspring control, both sexes help (OB) | | | |
| $\iota_p^{\mathrm{OB}}=\sum_{\ell\in\{♀,♂\}}\sigma_{1,\ell}u_\ell v_\ell$ | $=\sum_{\ell\in\{♀,♂\}}\underbrace{\sigma_{1,\ell}u_\ell}_{\phi_\ell(O_1)} r_{\bullet(O_{1\ell}),\bullet(O_{1\ell})}v_\ell$ | $=\sum_{\ell\in\{♀,♂\}}\phi_\ell(O_1)r_{\bullet(O_{1\ell}),\bullet(O_{1\ell})}v_\ell$ | $=\omega_{O_1,O_1}$ |
| $\kappa_p^{\mathrm{OB}}=\sum_{\ell\in\{♀,♂\}}\sigma_{1,\ell}\sum_{\ell'\in\{♀,♂\}}\sigma_{2,\ell'}\underbrace{\sum_{k\in\{\mathrm{mr},\mathrm{rm}\}}u_k q_{\ell,\mathrm{m},k}q_{\ell',\mathrm{m},k}}_{u_\ell r_{\bullet(O_{1\ell}),\circ(O_{2\ell'})}}v_{\ell'}$ | $=\sum_{\ell\in\{♀,♂\}}\underbrace{\sigma_{1,\ell}u_\ell}_{\phi_\ell(O_1)}\sum_{\ell'\in\{♀,♂\}}\underbrace{\sigma_{2,\ell'}}_{\sigma_{\ell'}(O_2)}r_{\bullet(O_{1\ell}),\circ(O_{2\ell'})}v_{\ell'}$ | $=\sum_{\ell\in\{♀,♂\}}\phi_\ell(O_1)\sum_{\ell'\in\{♀,♂\}}\sigma_{\ell'}(O_2)r_{\bullet(O_{1\ell}),\circ(O_{2\ell'})}v_{\ell'}$ | $=\omega_{O_1,O_2}$ |
| Offspring control, only females help (OF) | | | |
| $\iota_p^{\mathrm{OF}}=\sigma_{1,♀}u_♀ v_♀$ | $=\sigma_{1,♀}\underbrace{u_♀}_{\phi_♀(O_{1♀})}r_{\bullet(O_{1♀}),\bullet(O_{1♀})}v_♀$ | $=\sigma_{1,♀}\sum_{\ell\in\{♀,♂\}}\phi_\ell(O_{1♀})r_{\bullet(O_{1\ell}),\bullet(O_{1\ell})}v_\ell$ | $=\sigma_{1,♀}\omega_{O_{1♀},O_{1♀}}$ |
| $\kappa_p^{\mathrm{OF}}=\sigma_{1,♀}\sum_{\ell'\in\{♀,♂\}}\sigma_{2,\ell'}\underbrace{\sum_{k\in\{\mathrm{mr},\mathrm{rm}\}}u_k q_{♀,\mathrm{m},k}q_{\ell',\mathrm{m},k}}_{u_♀ r_{\bullet(O_{1♀}),\circ(O_{2\ell'})}}v_{\ell'}$ | $=\sigma_{1,♀}\underbrace{u_♀}_{\phi_♀(O_{1♀})}\sum_{\ell'\in\{♀,♂\}}\underbrace{\sigma_{2,\ell'}}_{\sigma_{\ell'}(O_2)}r_{\bullet(O_{1♀}),\circ(O_{2\ell'})}v_{\ell'}$ | $=\sigma_{1,♀}\sum_{\ell\in\{♀,♂\}}\phi_\ell(O_{1♀})\sum_{\ell'\in\{♀,♂\}}\sigma_{\ell'}(O_2)r_{\bullet(O_{1\ell}),\circ(O_{2\ell'})}v_{\ell'}$ | $=\sigma_{1,♀}\omega_{O_{1♀},O_2}$ |
| Maternal control, both sexes help (MB) | | | |
| $\iota_p^{\mathrm{MB}}=u_{\mathrm{mr}}\sum_{\ell\in\{♀,♂\}}\sigma_{1,\ell}\underbrace{q_{\ell,\mathrm{m},\mathrm{mr}}}_{r_{\bullet(M),\circ(O_{1\ell})}}v_\ell$ | $=\underbrace{u_♀}_{\phi_♀(M)}\sum_{\ell\in\{♀,♂\}}\underbrace{\sigma_{1,\ell}}_{\sigma_\ell(O_1)}r_{\bullet(M),\circ(O_{1\ell})}v_\ell$ | $=\sum_{\ell\in\{♀,♂\}}\phi_\ell(M)\sum_{\ell'\in\{♀,♂\}}\sigma_{\ell'}(O_1)r_{\bullet(M_\ell),\circ(O_{1\ell'})}v_{\ell'}$ | $=\omega_{M,O_1}$ |
| $\kappa_p^{\mathrm{MB}}=u_{\mathrm{mr}}\sum_{\ell'\in\{♀,♂\}}\sigma_{2,\ell'}\underbrace{q_{\ell',\mathrm{m},\mathrm{mr}}}_{r_{\bullet(M),\circ(O_{2\ell'})}}v_{\ell'}$ | $=\underbrace{u_♀}_{\phi_♀(M)}\sum_{\ell'\in\{♀,♂\}}\underbrace{\sigma_{2,\ell'}}_{\sigma_{\ell'}(O_2)}r_{\bullet(M),\circ(O_{2\ell'})}v_{\ell'}$ | $=\sum_{\ell\in\{♀,♂\}}\phi_\ell(M)\sum_{\ell'\in\{♀,♂\}}\sigma_{\ell'}(O_2)r_{\bullet(M_\ell),\circ(O_{2\ell'})}v_{\ell'}$ | $=\omega_{M,O_2}$ |
| Maternal control, only females help (MF) | | | |
| $\iota_p^{\mathrm{MF}}=\sigma_{1,♀}u_{\mathrm{mr}}\underbrace{q_{♀,\mathrm{m},\mathrm{mr}}}_{r_{\bullet(M),\circ(O_{1♀})}}v_♀$ | $=\sigma_{1,♀}\underbrace{u_♀}_{\phi_♀(M)}r_{\bullet(M),\circ(O_{1♀})}v_♀$ | $=\sigma_{1,♀}\sum_{\ell\in\{♀,♂\}}\phi_\ell(M)\sum_{\ell'\in\{♀,♂\}}\sigma_{\ell'}(O_{1♀})r_{\bullet(M_\ell),\circ(O_{1\ell'})}v_{\ell'}$ | $=\sigma_{1,♀}\omega_{M,O_{1♀}}$ |
| $\kappa_p^{\mathrm{MF}}=\sigma_{1,♀}u_{\mathrm{mr}}\sum_{\ell'\in\{♀,♂\}}\sigma_{2,\ell'}\underbrace{q_{\ell',\mathrm{m},\mathrm{mr}}}_{r_{\bullet(M),\circ(O_{2\ell'})}}v_{\ell'}$ | $=\sigma_{1,♀}\underbrace{u_♀}_{\phi_♀(M)}\sum_{\ell'\in\{♀,♂\}}\underbrace{\sigma_{2,\ell'}}_{\sigma_{\ell'}(O_2)}r_{\bullet(M),\circ(O_{2\ell'})}v_{\ell'}$ | $=\sigma_{1,♀}\sum_{\ell\in\{♀,♂\}}\phi_\ell(M)\sum_{\ell'\in\{♀,♂\}}\sigma_{\ell'}(O_2)r_{\bullet(M_\ell),\circ(O_{2\ell'})}v_{\ell'}$ | $=\sigma_{1,♀}\omega_{M,O_2}$ |

Figure S7: Structure coefficients in terms of reproductive worth. (A) Structure coefficients when helping is under offspring or maternal control, where either both sexes or only females help. Such structure coefficients after substituting for (B) relatedness; (C) the probability that an actor is mutant and of a given sex, and the probability that a recipient is of a given sex; and (D) reproductive worth. The structure coefficients when helping is under shared control, where either both sexes or only females help, are given by $\iota_x^{\mathrm{SC}}=(\partial p/\partial x)\iota_p^{\mathrm{MC}}$, $\kappa_x^{\mathrm{SC}}=(\partial p/\partial x)\kappa_p^{\mathrm{MC}}$, $\iota_y^{\mathrm{SC}}=(\partial p/\partial y)\iota_p^{\mathrm{OC}}$, and $\kappa_y^{\mathrm{SC}}=(\partial p/\partial y)\kappa_p^{\mathrm{OC}}$.

nest; we denote this individual by $\bullet(A_\ell)$. Then, we draw a recipient uniformly at random from the subset $R_{\ell'}$ of sex-$\ell'$ recipients in the nest; we denote this individual by $\circ(R_{\ell'})$.

**Definition of reproductive worth.** Based on the sampling experiment defined above, we define the *reproductive worth* for a random actor in $A$ of a random recipient in $R$ as

$$\omega_{A,R}=\begin{cases}\sum_{\ell\in\{♀,♂\}}\phi_\ell(A)r_{\bullet(A_\ell),\bullet(A_\ell)}v_\ell & \text{if } A=R\\ \sum_{\ell\in\{♀,♂\}}\phi_\ell(A)\sum_{\ell'\in\{♀,♂\}}\sigma_{\ell'}(R)r_{\bullet(A_\ell),\circ(R_{\ell'})}v_{\ell'} & \text{if } A\neq R,\end{cases}\tag{S3.2.1}$$

where (i) $r_{\bullet(A_\ell),\circ(R_{\ell'})}$ is the relatedness of actor $\bullet(A_\ell)$ to recipient $\circ(R_{\ell'})$, defined as the conditional probability that $\circ(R_{\ell'})$ is mutant given that $\bullet(A_\ell)$ is mutant (see section 3.2.1); (ii) $\phi_\ell(A)$ is the probability that an individual in $A$ is mutant and of sex $\ell$ (see section 3.2.2); and (iii) $\sigma_{\ell'}(R)$ is the probability that an individual in $R$ is of sex $\ell'$ (see section 3.2.3). Note that if the actor set is equal to the recipient set ($A=R$), reproductive worth is defined so that the random actor and the random recipient are the same individual (i.e., the focal individual $\bullet(A_\ell)$) so the relevant relatedness is $r_{\bullet(A_\ell),\bullet(A_\ell)}$. Given these definitions, reproductive worth $\omega_{A,R}$ is an inclusive fitness measure of how a random actor values its own reproduction (if $A=R$) or the reproduction of a random recipient (if $A\neq R$).

A

| Relatedness, $r$ of | to | Diploids | Haplodiploids |
|---|---|---|---|
| mother | daugther | $\frac{1}{2}$ | $\frac{1}{2}$ |
| | son | $\frac{1}{2}$ | $\frac{1}{2}$ |
| sister | sister | $\frac{1}{2}$ | $\frac{3}{4}$ |
| | brother | $\frac{1}{2}$ | $\frac{1}{4}$ |
| brother | sister | $\frac{1}{2}$ | $\frac{1}{2}$ |
| | brother | $\frac{1}{2}$ | $\frac{1}{2}$ |

B

| Relative worth, $\rho$ for | of | Diploids | Haplodiploids |
|---|---|---|---|
| mother | 2nd-brood offspring relative to 1st-brood offspring | 1 | 1 |
| 1st-brood sibling | 2nd-brood sibling relative to self | $\frac{1}{2}$ | $\frac{1}{2}$ |

when both sexes help and brood sex proportions are unbiased

Figure S8: Relatedness and relative reproductive worth. (A) Values of the relatedness coefficient $r$ we obtain. Taken from (S3.2.5), (S3.2.8), and (S3.2.9). (B) Values of relative reproductive worth $\rho$ when both sexes help (G = B) and brood sex proportions are unbiased ($\sigma_1 = \sigma_2 = 1/2$). Taken from (S3.3.4) and (S3.3.7).

**Outline.** In subsections 3.2.1, 3.2.2, and 3.2.3, we give details about the calculation of all the building blocks of our notion of reproductive worth. Then, in subsection 3.2.4 we show how to use these calculations to rewrite the structure coefficients $\iota$ and $\kappa$ in terms of reproductive worth, which we then use to obtain an inclusive fitness interpretation of the selection gradients.

##### 3.2.1 Relatedness

We define the relatedness $r_{i,j}$ of individual $i$ to individual $j$ as the conditional probability that $i$ is mutant given that $j$ is mutant, that is

$$\begin{aligned} r_{i,j} &= \Pr(j\text{'s genotype} = \text{m} \mid i\text{'s genotype} = \text{m}) \\ &= \frac{\Pr(i\text{'s genotype} = \text{m and } j\text{'s genotype} = \text{m})}{\Pr(i\text{'s genotype} = \text{m})}. \end{aligned} \tag{S3.2.2}$$

Our measure of relatedness takes the following values, summarized in Fig. S8A.

**Self-self ($r_{\bullet(A_\ell),\bullet(A_\ell)}$).** For any set of actors $A$, the relatedness of an actor to itself is

$$r_{\bullet(A_\ell),\bullet(A_\ell)} = 1, \tag{S3.2.3}$$

which is obtained from (S3.2.2) by letting $i = j = \bullet(A_\ell)$.

**Mother-offspring ($r_{\bullet(M),\circ(O_{a\ell})}$).** The relatedness of a mother to her offspring of sex $\ell$ is

$$r_{\bullet(M),\circ(O_{a\ell})} = \frac{u_♀ q_{\ell,♀}}{u_♀} = q_{\ell,♀} \;\; \forall a \in \{1,2\}. \tag{S3.2.4}$$

Indeed, the mother is a mutant with probability $u_{♀}$ so that both mother and offspring are mutants with probability $u_{♀}q_{\ell,♀}$. Simplifying, the relatedness of mother to offspring equals the transmission probability $q_{\ell,♀} \equiv q_{\ell,\mathrm{m,mr}}$.

For both diploids and haplodiploids, and from Fig. S4, we then get

$$\left(r_{\bullet(M),\circ(O_{1♀})}, r_{\bullet(M),\circ(O_{1♂})}\right) = \left(r_{\bullet M,\circ(O_{2♀})}, r_{\bullet(M),\circ(O_{2♂})}\right) = (1/2, 1/2)\,. \tag{S3.2.5}$$

Hence, irrespective of the genetic system and of the sex of the offspring, the relatedness of a mother to a random offspring is one half.

**Sibling-sibling ($r_{\bullet(O_{1\ell}),\circ(O_{2\ell'})}$).** Consider the relatedness of an individual to a (full) sibling. The conditional probability that a (second-brood) sibling of sex $\ell'$ is mutant given that a (first-brood) offspring of sex $\ell$ is mutant is given by

$$r_{\bullet(O_{1\ell}),\circ(O_{2\ell'})} = \frac{\sum_{k\in\{♀,♂\}} u_k q_{\ell,k} q_{\ell',k}}{\sum_{k\in\{♀,♂\}} u_k q_{\ell,k}} = \frac{\sum_{k\in\{♀,♂\}} u_k q_{\ell,k} q_{\ell',k}}{u_\ell}, \tag{S3.2.6}$$

where the second equality makes use of equation (S2.6.27). Indeed, a first-brood offspring is a mutant if either the mother is a mutant that transmits her mutant allele to the offspring (which happens with probability $u_{♀}q_{\ell,♀}$) or if the father is a mutant that transmits his mutant allele to the offspring (which happens with probability $u_{♂}q_{\ell,♂}$). Summing up the two probabilities, we obtain the total probability that a first-brood individual is a mutant, which is equal to $u_\ell$. This explains the denominator of the expression above. To calculate the numerator, we follow a similar logic, now noting that both offspring are mutants if either the mother is a mutant that transmits her mutant allele to both offspring (which happens with probability $u_{♀}q_{\ell,♀}q_{\ell',♀}$) or if the father is a mutant that transmits his mutant allele to both offspring (which happens with probability $u_{♂}q_{\ell,♂}q_{\ell',♂}$). Summing up the two probabilities we obtain the total probability that both offspring are mutants. The ratio of the two probabilities gives the conditional probability that both actor and recipient are mutants given that the actor is a mutant.

Note that, for a given sex of the actor, $\ell \in \{♀,♂\}$, $r_{\bullet(O_{1\ell}),\circ(O_{2\ell'})}$ defines a probability distribution over the possible sexes of the recipient, $\ell' \in \{♀,♂\}$. Indeed

$$\begin{aligned}
\sum_{\ell'\in\{♀,♂\}} r_{\bullet(O_{1\ell}),\circ(O_{2\ell'})} &= \sum_{\ell'\in\{♀,♂\}} \frac{\sum_{k\in\{♀,♂\}} u_k q_{\ell,k} q_{\ell',k}}{u_\ell} \\
&= \frac{1}{u_\ell} \sum_{k\in\{♀,♂\}} u_k q_{\ell,k} \sum_{\ell'\in\{♀,♂\}} q_{\ell',k} \\
&= \frac{1}{u_\ell} \sum_{k\in\{♀,♂\}} u_k q_{\ell,k} \\
&= 1,
\end{aligned} \tag{S3.2.7}$$

where the first line substitutes the formula given in equation (S3.2.6), the second line rearranges, the third line applies identity (S1.2.1d), and the last equality results from applying (S2.6.27).

For diploids, and from (S2.6.24) and Fig. S4, we obtain

$$\left(r_{\bullet(O_{1♀}),\circ(O_{2♀})}, r_{\bullet(O_{1♀}),\circ(O_{2♂})}, r_{\bullet(O_{1♂}),\circ(O_{2♀})}, r_{\bullet(O_{1♂}),\circ(O_{2♂})}\right) = (1/2, 1/2, 1/2, 1/2)\,, \tag{S3.2.8}$$

so that the relatedness of an individual to any sibling is, irrespective of the sexes of actor and recipient, equal to one half.

For haplodiploids, and from (S2.6.24) and Fig. S4, we get

$$\left(r_{\bullet(O_{1♀}),\circ(O_{2♀})}, r_{\bullet(O_{1♀}),\circ(O_{2♂})}, r_{\bullet(O_{1♂}),\circ(O_{2♀})}, r_{\bullet(O_{1♂}),\circ(O_{2♂})}\right) = (3/4, 1/4, 1/2, 1/2)\,. \tag{S3.2.9}$$

Here, the asymmetry of the transmission probabilities for the case of haplodiploids makes a female offspring more related to a sister than to a brother, while a male offspring is equally related to both sisters and brothers.

**Connection to other relatedness coefficients.** Our relatedness coefficients are conceptually most similar to the *weighted pedigree relatedness coefficients* of [23] (p. 190; $G'$ in their notation). Such weighted relatedness involves pedigree relatedness weighted by the so-called genetic reproductive values (which we have seen to arise in our model as the stable sex distribution rather than as reproductive values). Indeed, the stable sex distribution is part of our relatedness coefficients $r$ (equation (S3.2.4) and (S3.2.6)). Hamilton's notion of *complete* or *life-for-life relatedness coefficients* includes both the stable sex distribution (described by a factor 2 multiplying $c$ in his cross-sex formulas in Table 1; [14]), and the sex ratio (his $c$), which we have seen to arise in our model as reproductive values. Accordingly, the values for our relatedness coefficients (equations (S3.2.5), (S3.2.8), and (S3.2.9)) numerically recover the standard values for Hamilton's life-for-life relatedness coefficients for the case of singly-mated, outbred queens, and unbiased sex ratio; e.g., p. 81 of [24].

##### 3.2.2 Probability that an actor is mutant and of a given sex

$\phi_\ell(A)$ in (S3.2.1) denotes the probability that an actor (i.e., an individual in $A$) is mutant and of sex $\ell$. This probability takes the following values.

**Actors are first-brood offspring ($A = O_1$).** If the set of actors is the set of first-brood offspring, the probability that an actor is mutant and of sex $\ell$ is

$$\phi_\ell(O_1) = \sigma_{1,\ell} u_\ell, \tag{S3.2.10}$$

since a first-brood offspring is of sex $\ell$ with probability $\sigma_{1,\ell}$ and it is a mutant with probability $u_\ell$ due to equation (S2.6.27).

**Actors are first-brood female offspring ($A = O_{1♀}$).** If the set of actors is the set of first-brood female offspring, the probability that an actor is mutant and of sex $\ell$ is given by

$$\phi_\ell(O_{1♀}) = \begin{cases} u_\ell & \text{if } \ell = ♀ \\ 0 & \text{if } \ell = ♂. \end{cases} \tag{S3.2.11}$$

Indeed, a first-brood female offspring is of sex ♀ with probability 1 and it is mutant with probability $u_\ell$ due to equation (S2.6.27); by definition, a first-brood female offspring is of sex ♂ with probability 0.

**Actors are mothers ($A = M$).** If the set of actors is the mother singleton, the probability that an actor is mutant and of sex $\ell$ is

$$\phi_\ell(M) = \begin{cases} u_\ell & \text{if } \ell = \venus \\ 0 & \text{if } \ell = \mars. \end{cases} \tag{S3.2.12}$$

Indeed, a mother is of sex ♀ with probability 1 and it is mutant with probability $u_\ell$ due to equation (S2.6.22); by definition, a mother is of sex ♂ with probability 0.

##### 3.2.3 Probability that a recipient is of a given sex

$\sigma_{\ell'}(R)$ in (S3.2.1) denotes the probability that a recipient (i.e., an individual in $R$) is of sex $\ell'$. This probability takes the following value.

**Recipients are $a$-th brood offspring ($R = O_a$).** Consider the case where the set of recipients is the set of $a$-th brood offspring. The probability that an $a$-th brood offspring is of sex $\ell'$ is

$$\sigma_{\ell'}(O_a) = \sigma_{a,\ell'}. \tag{S3.2.13}$$

##### 3.2.4 Structure coefficients in terms of reproductive worth

We can obtain an inclusive fitness interpretation of the selection gradients by rewriting the structure coefficients $\iota$ and $\kappa$ in terms of reproductive worth (S3.2.1), for each of our model cases. These equivalences and their derivation are summarized in Fig. S7. Substituting equations (S3.2.3), (S3.2.4), and (S3.2.6) into Fig. S7A yields Fig. S7B. Substituting equations (S3.2.10), (S3.2.11), (S3.2.12), and (S3.2.13) into Fig. S7B yields Fig. S7C. In turn, substituting equation (S3.2.1) into Fig. S7C yields Fig. S7D which expresses the structure coefficients in terms of reproductive worth.

Overall, we have shown that the structure coefficients can be written in terms of reproductive worth with the calculated expressions for the probability that an actor of a given sex and a recipient of a given sex carry a mutation given that the actor carries it ($r_{\bullet(A_\ell),\circ(R'_\ell)}$ and $r_{\bullet(A_\ell),\bullet(A_\ell)}$), the probability that an actor is mutant given that it is of a given sex ($\phi_\ell(A)$), and the probability that a recipient is of a given sex ($\sigma_{\ell'}(R)$) (Fig. S7). In doing this, we find that candidate recipients of help (i.e., the payees) are second-brood offspring for all our model cases (Fig. S7). For instance, even if helping increases couple survival, payees are still second-brood offspring and the relevant relatedness is that toward such offspring rather than toward the couple.

#### 3.3 Relative reproductive worth

In order to write more compact expressions, we define the *relative reproductive worth*, $\rho_{A,H,P}$, for a random actor in $A$ relative to a random candidate helper in $H$ of a random payee in $P$ as

$$\rho_{A,H,P} = \frac{\omega_{A,P}}{\omega_{A,H}}, \tag{S3.3.1}$$

that is, as the ratio between the reproductive worth $\omega_{A,P}$ (measuring how much a random actor from $A$ values the reproduction of a random payee from $P$) and the reproductive worth $\omega_{A,H}$ (measuring how much a random actor from $A$ values the reproduction of a random candidate helper from $H$). In the main text, we write $\rho_A$

as short-hand of $\rho_{A,H,P}$. Our measure of relative reproductive worth can be seen as a generalization of the concept of life-for-life relatedness coefficients introduced by [14] to allow for actors and recipients to be of both sexes.

Relative reproductive worth $\rho_{A,H,P}$ takes the following values, summarized for the cases when both sexes help and brood sex proportions are unbiased in Fig. S8B.

**Sibling-sibling-sibling for both females and males ($\rho_{O_1,O_1,O_2}$).** The relative reproductive worth $\rho_{O_1,O_1,O_2}$ for a random first-brood offspring actor relative to itself of a random second-brood offspring recipient is given by

$$\rho_{O_1,O_1,O_2} = \frac{\omega_{O_1,O_2}}{\omega_{O_1,O_1}} = \frac{\sum_{\ell \in \{♀,♂\}} \phi_\ell(O_1) \sum_{\ell' \in \{♀,♂\}} \sigma_{2,\ell'} r_{\bullet(O_{1\ell}),\circ(O_{2\ell'})} v_{\ell'}}{\sum_{\ell \in \{♀,♂\}} \phi_\ell(O_1) v_\ell}. \tag{S3.3.2}$$

This expression greatly simplifies for two particular but relevant cases. First, for diploids, and via Fig. S7, we get

$$\rho_{O_1,O_1,O_2} = \frac{1}{2} \frac{\sum_{\ell \in \{♀,♂\}} \sigma_{2,\ell} v_\ell}{\sum_{\ell \in \{♀,♂\}} \sigma_{1,\ell} v_\ell}. \tag{S3.3.3}$$

Second, if both sexes help (G = B) and brood sex proportions are unbiased (i.e., $\sigma_1 = \sigma_2 = 1/2$), so that $v_♀ = v_♂ = 1$ also holds, (S3.3.2) can be simplified as

$$\begin{aligned} \rho_{O_1,O_1,O_2} &= \frac{\sum_{\ell \in \{♀,♂\}} \phi_\ell(O_1) \sum_{\ell' \in \{♀,♂\}} \sigma_{2,\ell'} r_{\bullet(O_{1\ell}),\circ(O_{2\ell'})}}{\sum_{\ell \in \{♀,♂\}} \phi_\ell(O_1)} \\ &= \frac{1}{2} \sum_{\ell \in \{♀,♂\}} u_\ell \sum_{\ell' \in \{♀,♂\}} r_{\bullet(O_{1\ell}),\circ(O_{2\ell'})} \\ &= \frac{1}{2} \sum_{\ell \in \{♀,♂\}} u_\ell \\ &= \frac{1}{2}, \end{aligned} \tag{S3.3.4}$$

where the first line follows from substituting (S3.3.2) with $v_♀ = v_♂ = 1$; the second line substitutes $\sigma_{2,♀} = \sigma_{2,♀} = 1/2$, and identifies $\phi_\ell(O_1) = u_\ell$; the third line applies identity (S3.2.7); and the fourth line simplifies.

**Mother-offspring-offspring ($\rho_{M,O_1,O_2}$).** The relative reproductive worth $\rho_{M,O_1,O_2}$ for a mother relative to a random candidate first-brood offspring helper of a random second-brood offspring payee is given by

$$\rho_{M,O_1,O_2} = \frac{\omega_{M,O_2}}{\omega_{M,O_1}} = \frac{\sum_{\ell \in \{♀,♂\}} \sigma_{2,\ell} r_{\bullet(M),\circ(O_{2\ell})} v_\ell}{\sum_{\ell \in \{♀,♂\}} \sigma_{1,\ell} r_{\bullet(M),\circ(O_{1\ell})} v_\ell}, \tag{S3.3.5}$$

which, for both diploids and haplodiploids, simplifies to

$$\rho_{M,O_1,O_2} = \frac{\sum_{\ell \in \{♀,♂\}} \sigma_{2,\ell} v_\ell}{\sum_{\ell \in \{♀,♂\}} \sigma_{1,\ell} v_\ell}. \tag{S3.3.6}$$

If, additionally, both sexes help (G = B) and brood sex proportions are unbiased (i.e., $\sigma_1 = \sigma_2 = 1/2$), so that $v_♀ = v_♂ = 1$ also holds, then

$$\rho_{M,O_1,O_2} = 1. \tag{S3.3.7}$$

**Sibling-sibling-sibling for females ($\rho_{O_{1♀},O_{1♀},O_2}$).** The relative reproductive worth $\rho_{O_{1♀},O_{1♀},O_2}$ for a random first-brood female offspring actor relative to herself of a random second-brood offspring payee is given by

$$\rho_{O_{1♀},O_{1♀},O_2} = \frac{\omega_{O_{1♀},O_2}}{\omega_{O_{1♀},O_{1♀}}} = \sum_{\ell \in \{♀,♂\}} \sigma_{2,\ell} r_{\bullet(O_{1♀}),\circ(O_{2\ell})} \frac{v_\ell}{v_♀}. \tag{S3.3.8}$$

If only female offspring were produced, then $\sigma_{2,♀} = 1$ and $\sigma_{2,♂} = 0$ so the relative reproductive worth for a random first-brood female offspring actor relative to herself of a random second-brood sister payee reduces to

$$\rho_{O_{1♀},O_{1♀},O_2} = r_{\bullet(O_{1♀}),\circ(O_{2♀})},$$

as stated in the main text.

**Mother-daughter-offspring ($\rho_{M,O_{1♀},O_2}$).** The relative reproductive worth $\rho_{M,O_{1♀},O_2}$ for a mother relative to a random candidate first-brood daughter helper of a random second-brood offspring payee is given by

$$\rho_{M,O_{1♀},O_2} = \frac{\omega_{M,O_2}}{\omega_{M,O_{1♀}}} = \frac{\sum_{\ell \in \{♀,♂\}} \sigma_{2,\ell} r_{\bullet(M),\circ(O_{2\ell})} v_\ell}{r_{\bullet(M),\circ(O_{1♀})} v_♀}. \tag{S3.3.9}$$

If only female offspring were produced, then $\sigma_{2,♀} = 1$ and $\sigma_{2,♂} = 0$ so the relative reproductive worth for a mother relative to a random candidate first-brood daughter helper of a random second-brood daughter payee reduces to

$$\rho_{M,O_{1♀},O_2} = \frac{r_{\bullet(M),\circ(O_{2♀})} v_♀}{r_{\bullet(M),\circ(O_{1♀})} v_♀} = \frac{r_{\bullet(M),\circ(O_{2♀})}}{r_{\bullet(M),\circ(O_{1♀})}} = 1,$$

as stated in the main text.

#### 3.4 Individual cost and benefit of helping

The cost $C$ (S2.8.4) and the benefit $B$ (S2.8.5) of helping refer to the marginal effects of changing the number of helpers on either the early or the late productivity of a couple. These quantities can also be written in terms of inclusive fitness, which considers the effect that an individual candidate helper $i \in H$ has, respectively, on its own personal fitness and on the fitness of its payees (all members of $P$). Such individual interpretations of cost and benefit of helping are the last building block we need in order to interpret the selection gradients from an inclusive fitness perspective.

For these purposes, let us define the personal fitness of a first or second-brood offspring as their personal contribution to the stages of unmated reproductives. Now consider a focal individual $i$ belonging to the set of candidate helpers $H$. Denoting by $p_i$ the probability that $i$ becomes a helper, the personal fitness of $i$ is then given by

$$W_{1,i} = (1 - p_i) s_1, \tag{S3.4.1}$$

while the expected total fitness of individuals belonging to $P$ is

$$W_2 = \Pi_2(f_2, h). \tag{S3.4.2}$$

The marginal effects of the trait $\zeta$ affecting helping of a focal candidate helper on its own personal fitness and on the total fitness of its second-brood offspring are then respectively given by

$$-c_\zeta \equiv \frac{\partial W_{1,i}}{\partial \zeta_i} = \frac{\partial W_{1,i}}{\partial p_i}\frac{\partial p_i}{\partial \zeta_i} = -s_1\frac{\partial p}{\partial \zeta} = -C\frac{\partial p}{\partial \zeta}, \tag{S3.4.3a}$$

$$b_\zeta \equiv \frac{\partial W_2}{\partial \zeta_i} = \frac{\partial W_2}{\partial p_i}\frac{\partial p_i}{\partial \zeta_i} = \frac{\partial \Pi_2}{\partial h}(f_2,h)\frac{\partial h}{\partial p_i}\frac{\partial p}{\partial \zeta} = \frac{\partial \Pi_2}{\partial h}(f_2,h)\frac{\partial p_i}{\partial \zeta_i} = B\frac{\partial p}{\partial \zeta}, \tag{S3.4.3b}$$

where we have used the fact that $\partial h/\partial p_i = 1$, since the number of helpers can be written as

$$h = p_i + \sum_{j\in H, j\neq i} p_j$$

and the probabilities $p_\ell$ for all $\ell \in H$ are assumed to be independent.

Thus, the benefit $B$ and cost $C$ equal the inclusive fitness benefit $b_\zeta$ and cost $c_\zeta$ when the trait is the helping probability $\zeta = p$.

#### 3.5 Inclusive fitness effect for a trait affecting helping and Hamilton's rule

We have obtained expressions for the selection gradient of a trait $\zeta$ affecting helping for all the model cases we study in terms of structure coefficients (equations (S2.8.40) and (S2.8.41)). We have also shown how such structure coefficients translate into inclusive fitness measures of reproductive valuation (Fig. S7). Finally, we have also obtained expressions for the individual benefit and cost (equations (S3.4.3)). Using these results and the definition of the maximum number of helpers $\bar{h}$ (equation (S1.1.5)), it follows that the selection gradient of a trait $\zeta$ affecting helping for all the model cases we study can be written as

$$\mathcal{S}_\zeta^{\mathrm{C,G}} = \frac{\bar{h}}{\mathbf{v}^\intercal\mathbf{u}}\mathcal{H}_\zeta^{\mathrm{C,G}}, \tag{S3.5.1}$$

where we define the inclusive fitness effect of a trait $\zeta$ affecting helping as

$$\mathcal{H}_\zeta^{\mathrm{C,G}} = -\omega_{A,H}c_\zeta + \omega_{A,P}b_\zeta. \tag{S3.5.2}$$

Indeed, $\mathcal{H}_\zeta^{\mathrm{C,G}}$ is the marginal effect of a candidate helper's phenotype on the candidate helper's personal fitness $(-c_\zeta)$ weighted by how much a random actor values the reproduction of a random candidate helper $(\omega_{A,H})$ plus the marginal effect of a candidate helper's phenotype on the fitness of payees $(b_\zeta)$ weighted by how much a random actor values the reproduction of a random payee $(\omega_{A,P})$.

Therefore, for all the model cases we consider, a trait $\zeta$ affecting helping is favored by selection if and only if

$$\underbrace{-\omega_{A,H}c_\zeta + \omega_{A,P}b_\zeta}_{\mathcal{H}_\zeta^{\mathrm{C,G}}} > 0. \tag{S3.5.3}$$

Condition (S3.5.3) constitutes a Hamilton's rule for the model cases we consider [19, 20, 21, 22]. Resistance is thus a selfish trait (both $c_y < 0$ and $b_y < 0$) according to the terminology of [25].

Dividing by $\omega_{A,H}$ (which is strictly positive), a trait $\zeta$ affecting helping is favored by selection if and only if

$$-c_\zeta + \rho_{A,H,P}b_\zeta > 0,$$

where $\rho_{A,H,P}$ is the relative worth for a random actor in $A$ relative to a random candidate helper in $H$ of a random payee in $P$. Specifically, if the trait is the helping probability $\zeta = p$, helping is favored by selection if and only if

$$-C + \rho_{A,H,P} B > 0. \tag{S3.5.4}$$

Then, for all the model cases we consider, the critical benefit-cost ratio (S2.8.10) can be alternatively written as

$$\left(\frac{B}{C}\right)^* = \frac{1}{\rho_{A,H,P}}. \tag{S3.5.5}$$

#### 3.6 Inclusive fitness effect for reproductive effort

We have obtained the selection gradient of reproductive effort $z$ for all the model cases we study in terms of the structure coefficient $\kappa_z^{\mathrm{C,G}}$ (S2.9.6). We have shown how this structure coefficient translates into an inclusive fitness measure of reproductive valuation; specifically, it equals $\omega_{M,O_2}$ (Fig. S7). We can define the individual benefit for a mother of increasing her reproductive effort $z$ as

$$b_z \equiv \frac{\partial W_2}{\partial z} = \frac{\partial \Pi_2}{\partial f_2}\frac{\mathrm{d}f_2}{\mathrm{d}z} = D\frac{\mathrm{d}f_2}{\mathrm{d}z}. \tag{S3.6.1}$$

Using these results, it follows that the selection gradient of reproductive effort $z$ for all the model cases we study is

$$\mathcal{S}_z^{\mathrm{C,G}} = \frac{1}{\mathbf{v}^\intercal \mathbf{u}} \mathcal{H}_z^{\mathrm{C,G}}, \tag{S3.6.2}$$

where we define the inclusive fitness effect of reproductive effort $z$ as

$$\mathcal{H}_z^{\mathrm{C,G}} = \omega_{M,O_2} b_z. \tag{S3.6.3}$$

### 4 Conflict dissolution and benefit-cost ratio zones

In this section, we define conflict dissolution and show that it can also be understood in terms of benefit-cost ratios zones. To do this, we proceed in three steps. First, we define zones for the benefit-cost ratio in which a party (i.e., the mother or the offspring) favors or disfavors increasing helping (Benefit-cost ratio zones considering the interest of a single party; section 4.1). Second, we define benefit-cost ratio zones considering simultaneously the interests of both mother and offspring, and define the zone of parent-offspring conflict over helping (Benefit-cost ratio zones simultaneously considering the interest of mother and offspring; section 4.2). Third, we define *conflict dissolution* and show how it can be understood in terms of benefit-cost ratio zones (Conflict dissolution; section 4.3).

#### 4.1 Benefit-cost ratio zones considering the interest of a single party

To define the benefit-cost ratio zones, recall the following. We have obtained that an increasing helping probability $p$ is favored by selection if and only if

$$\frac{B}{C} > \left(\frac{B}{C}\right)^{*} \tag{S4.1.1}$$

(equations (S2.8.8) and (S2.8.9a) since $\kappa > 0$ for $\zeta = p$). We have also obtained that the critical benefit-cost ratio $(B/C)^{*}$ can be written in inclusive fitness terms as

$$\left(\frac{B}{C}\right)^{*} = \frac{1}{\rho_{A,H,P}}$$

for all the model cases we consider (equation (S3.5.4)). Finally, we have seen that the critical benefit-cost ratio depends on the model case, which when useful we highlight by writing $(B/C)^{*} = (B/C)_{p}^{*^{\mathrm{C,G}}}$ for the helping probability $p$ (Fig. S9A).

When helping is under the control of a single party, that is, when helping is under offspring or maternal control, we have the following benefit-cost ratio zones (Fig. S9B):

1. Low benefit-cost ratio ($B/C < (B/C)^{*}$). In this zone, the selection gradient of helping, $\mathcal{S}_{p}(\mathbf{z})$, is negative, so helping is disfavored by selection. As the helping trait is either under maternal or offspring control, we say that helping is disfavored by the party controlling helping.

2. High benefit-cost ratio ($B/C > (B/C)^{*}$). In this zone, the selection gradient of helping, $\mathcal{S}_{p}(\mathbf{z})$, is positive, so helping is favored by selection. We say that helping is favored by the party controlling helping.

We can show that if the genetic system is diploid, if only females help, or if brood sex proportions are unbiased, that is, if at least one of the following conditions is satisfied:

$$\mathrm{P} = \mathrm{D}, \tag{S4.1.2a}$$

$$\mathrm{G} = \mathrm{F}, \tag{S4.1.2b}$$

$$\sigma_1 = \sigma_2 = 1/2, \tag{S4.1.2c}$$

then

$$\left(\frac{B}{C}\right)_{p}^{*^{\mathrm{M,G}}} < \left(\frac{B}{C}\right)_{p}^{*^{\mathrm{O,G}}} \tag{S4.1.3}$$

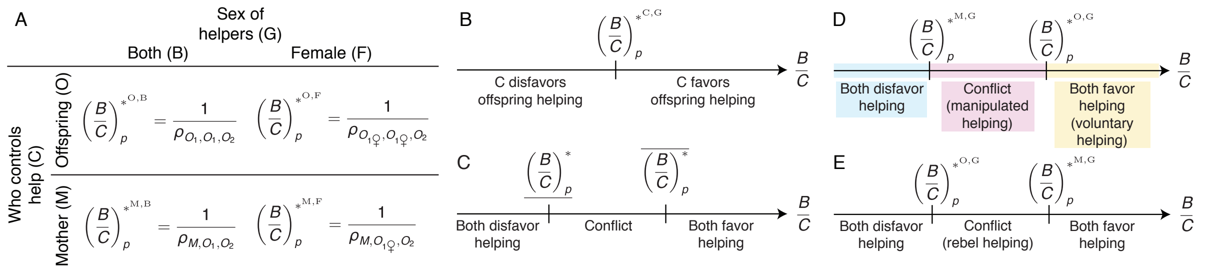

Figure S9: Benefit-cost ratio zones. (A) Critical benefit-cost ratio for helping for all model cases and its corresponding inclusive fitness interpretation (equations (S3.5.5), (S3.1.1), and (S3.1.2)). (B) Benefit-cost ratio zones considering helping control by a single party. Who controls help is given by C (for C ∈ {O, M}, where O stands for offspring control and M stands for maternal control). (C-E) Benefit-cost ratio zones simultaneously considering helping control by mother and offspring, (D) when condition (S4.1.3) holds and (E) when the reverse of condition (S4.1.3) holds. Throughout, we consider only the case when (S4.1.3) holds (D).

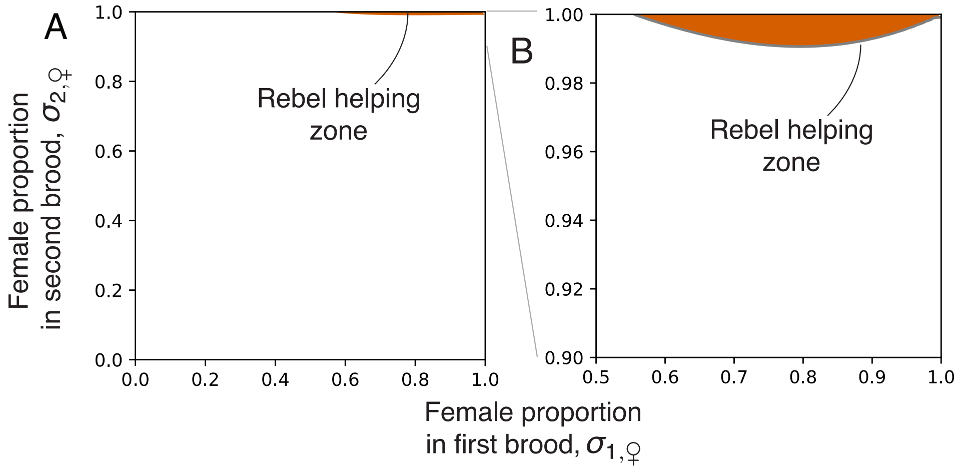

Figure S10: Rebel helping zone. In the case of haplodiploids where both sexes help, the reverse of inequality (S4.1.3) holds in the red zone. (A) In full brood-sex-proportion space. (B) In "zoomed" brood-sex-proportion space. Parameter values are: $f_1 = 30$, $f_2 = 60$, $s_1 = 0.2$, $s_2 = 0.5$, $s_M = 0.9$, and $p = 0.5$.

holds, in which case the helping zone is greater when helping is under maternal control than under offspring control. Note that at least one out of the three assumptions listed in (S4.1.2) holds in all of our model cases, except for the case of haplodiploids where both sexes help (HD-C-B) with biased sex proportions ($\sigma_1 \neq \sigma_2$). In such a case, the reverse of inequality (S4.1.3) can hold in a thin band of extremely female biased sex proportions (Fig. S10). Yet, such a case might be of limited biological interest as known real populations of haplodiploids where both sexes help are characterized by unbiased sex proportions [5, 6].

We now show that if any of the assumptions listed in (S4.1.2) holds, then (S4.1.3) holds. First, let us consider case (S4.1.2b) (i.e., only females help), for which (S4.1.3) takes the form

$$\left(\frac{B}{C}\right)_p^{*\mathrm{M,F}} < \left(\frac{B}{C}\right)_p^{*\mathrm{O,F}}. \qquad \text{(S4.1.4)}$$

Via the expressions in Fig. S9A, and using (S3.3.8) and (S3.3.9), inequality (S4.1.4) simplifies to

$$\frac{r_{\bullet(M),\circ(O_{1♀})}}{\sum_{\ell\in\{♀,♂\}}\sigma_{2,\ell}r_{\bullet(M),\circ(O_{2\ell})}v_\ell} < \frac{1}{\sum_{\ell\in\{♀,♂\}}\sigma_{2,\ell}r_{\bullet(O_{1♀}),\circ(O_{2\ell})}v_\ell}. \tag{S4.1.5}$$

For both diploids and haplodiploids, we have that $r_{\bullet(M),\circ(O_{a\ell})} = 1/2$ for all $\ell \in \{♀,♂\}$ and all $a \in \{1,2\}$ (from equation (S3.2.5)), so (S4.1.5) simplifies to

$$\frac{1}{\sum_{\ell\in\{♀,♂\}}\sigma_{2,\ell}v_\ell} < \frac{1}{\sum_{\ell\in\{♀,♂\}}\sigma_{2,\ell}r_{\bullet(O_{1♀}),\circ(O_{2\ell})}v_\ell},$$

which rearranging yields

$$\sum_{\ell\in\{♀,♂\}}\sigma_{2,\ell}\left(1-r_{\bullet(O_{1♀}),\circ(O_{2\ell})}\right)v_\ell > 0.$$

This holds true since $r_{\bullet(O_{1♀}),\circ(O_{2\ell})} < 1$ always holds (from equations (S3.2.8) and (S3.2.9)). We conclude that (S4.1.4) is true for both diploids and haplodiploids.

Now, let us consider case (S4.1.2a) (i.e., the genetic system is diploid). Since (S4.1.4) has been established irrespectively of the genetic system, we only need to consider the case where both sexes help (G = B), that is

$$\left(\frac{B}{C}\right)_p^{*\text{M,B}} < \left(\frac{B}{C}\right)_p^{*\text{O,B}}. \tag{S4.1.6}$$

for diploids. This inequality follows by substituting from (S3.3.3) and (S3.3.6).

Finally, let us assume that (S4.1.2c) holds (i.e., brood sex proportions are unbiased). Since (S4.1.4) has been established irrespectively of the brood sex proportions, we only need to consider the case where both sexes help (G = B). Then, via equations (S3.3.4) and (S3.3.7), we have that

$$\left(\frac{B}{C}\right)_p^{*\text{M,B}} = 1, \tag{S4.1.7a}$$

$$\left(\frac{B}{C}\right)_p^{*\text{O,B}} = 2 \tag{S4.1.7b}$$

holds, and (S4.1.3) is satisfied.

#### 4.2 Benefit-cost ratio zones simultaneously considering the interest of mother and offspring

Considering the interests of both mother and offspring simultaneously, we have two critical benefit-cost ratios: one corresponding to helping under maternal control ($(B/C)_p^{*\text{M,G}}$) and one corresponding to helping under offspring control ($(B/C)_p^{*\text{O,G}}$). Defining the *minimum critical benefit-cost ratio*,

$$\underline{\left(\frac{B}{C}\right)^*} \equiv \min\left(\left(\frac{B}{C}\right)_p^{*\text{O,G}}, \left(\frac{B}{C}\right)_p^{*\text{M,G}}\right), \tag{S4.2.1}$$

and the *maximum critical benefit-cost ratio*,

$$\overline{\left(\frac{B}{C}\right)^*} \equiv \max\left(\left(\frac{B}{C}\right)_p^{*\text{O,G}}, \left(\frac{B}{C}\right)_p^{*\text{M,G}}\right), \tag{S4.2.2}$$

we have the following three benefit-cost ratios zones (Fig. S9C-E):

1. Low benefit-cost ratio ($B/C < \underline{(B/C)^*}$). In this zone, the selection gradients of helping under maternal control and under offspring control, $\mathcal{S}_p^{\mathrm{M,G}}(\mathbf{z})$ and $\mathcal{S}_p^{\mathrm{O,G}}(\mathbf{z})$, are both negative. Hence, we say that helping is disfavored from both the mother's and offspring's perspective.

2. Intermediate benefit-cost-ratio ($\underline{(B/C)^*} < B/C < \overline{(B/C)^*}$). In this zone, the selection gradients of helping under maternal control and under offspring control, $\mathcal{S}_p^{\mathrm{M,G}}(\mathbf{z})$ and $\mathcal{S}_p^{\mathrm{O,G}}(\mathbf{z})$, have opposite sign. Thus, helping is favored (resp. disfavored) from the mother's perspective and disfavored (resp. favored) from the offspring's perspective. Hence, we say that there is parent-offspring conflict over helping. There are two possibilities:

   (a) If $(B/C)_p^{*\mathrm{M,G}} < (B/C)_p^{*\mathrm{O,G}}$ holds, so that $\underline{(B/C)^*} = (B/C)_p^{*\mathrm{M,G}}$ and $\overline{(B/C)^*} = (B/C)_p^{*\mathrm{O,G}}$, the selection gradient of helping under maternal control, $\mathcal{S}_p^{\mathrm{M,G}}(\mathbf{z})$, is positive and the selection gradient of helping under offspring control, $\mathcal{S}_p^{\mathrm{O,G}}(\mathbf{z})$, is negative. Hence, helping is favored from the mother's perspective but is disfavored from the offspring's perspective. We call "manipulated helping" the helping that is in this zone.

   (b) If $(B/C)_p^{*\mathrm{O,G}} < (B/C)_p^{*\mathrm{M,G}}$ holds, so that $\underline{(B/C)^*} = (B/C)_p^{*\mathrm{O,G}}$ and $\overline{(B/C)^*} = (B/C)_p^{*\mathrm{M,G}}$, the selection gradient of helping under maternal control, $\mathcal{S}_p^{\mathrm{M,G}}(\mathbf{z})$, is negative and the selection gradient of helping under offspring control, $\mathcal{S}_p^{\mathrm{O,G}}(\mathbf{z})$, is positive. Hence, helping is disfavored from the mother's perspective but is favored from the offspring's perspective. We call "rebel helping" the helping that is in this zone. As shown above, this case only occurs for haplodiploids where both sexes help and with extremely female biased sex proportions (Fig. S10). We do not study this case.

3. High benefit-cost ratio ($B/C > \overline{(B/C)^*}$). In this zone, the selection gradients of helping under maternal and under offspring control, $\mathcal{S}_p^{\mathrm{M,G}}(\mathbf{z})$ and $\mathcal{S}_p^{\mathrm{O,G}}(\mathbf{z})$, are both positive. Hence, helping is favored from both the mother's and the offspring's perspective. We call "voluntary helping" the helping that is in this zone.

##### 4.3 Conflict dissolution

We say that conflict dissolution occurs if there are evolutionary times $\tau_0 < \tau_{\mathrm{end}}$ such that

$$\mathcal{S}_p^{\mathrm{M,G}}(\mathbf{z}(\tau_0)) > 0, \quad \mathcal{S}_p^{\mathrm{O,G}}(\mathbf{z}(\tau_0)) < 0, \quad \mathcal{S}_p^{\mathrm{M,G}}(\mathbf{z}(\tau_{\mathrm{end}})) > 0, \quad \text{and} \quad \mathcal{S}_p^{\mathrm{O,G}}(\mathbf{z}(\tau_{\mathrm{end}})) > 0, \tag{S4.3.1}$$

that is, helping is favored by the mother and disfavored by offspring at time $\tau_0$, and helping is favored by both mother and offspring at time $\tau_{\mathrm{end}}$. Given equation (S3.5.1), conditions (S4.3.1) are equivalent to

$$\mathcal{H}_p^{\mathrm{M,G}}(\mathbf{z}(\tau_0)) > 0, \quad \mathcal{H}_p^{\mathrm{O,G}}(\mathbf{z}(\tau_0)) < 0, \quad \mathcal{H}_p^{\mathrm{M,G}}(\mathbf{z}(\tau_{\mathrm{end}})) > 0, \quad \text{and} \quad \mathcal{H}_p^{\mathrm{O,G}}(\mathbf{z}(\tau_{\mathrm{end}})) > 0. \tag{S4.3.2}$$

Provided that $\mathcal{H}_p^{\mathrm{O,G}}(\mathbf{z}(\tau))$ is everywhere differentiable with respect to $\tau$, conditions (S4.3.2) imply that the inclusive fitness effect for offspring-controlled helping satisfies the following: there exists an interval $[\tau_1, \tau_2] \subset [\tau_0, \tau_{\mathrm{end}})$ such that

1. $\mathcal{H}_p^{\mathrm{O,G}}(\mathbf{z}(\tau))$ increases with evolutionary time during $[\tau_1, \tau_2]$, that is,

$$\frac{\mathrm{d}\mathcal{H}_p^{\mathrm{O,G}}}{\mathrm{d}\tau}(\mathbf{z}(\tau)) > 0 \qquad \text{(persuasion condition)}$$

   for all $\tau \in [\tau_1, \tau_2]$, and

2. $\mathcal{H}_p^{\text{O,G}}(\mathbf{z}(\tau))$ is null at some evolutionary time within $[\tau_1, \tau_2]$, that is,

$$\mathcal{H}_p^{\text{O,G}}(\mathbf{z}(\tau)) = 0 \qquad \text{(conversion condition)}$$

for some $\tau \in (\tau_1, \tau_2)$.

The persuasion condition and conversion condition state that $\mathcal{H}_p^{\text{O,G}}(\mathbf{z}(\tau))$ changes sign from negative to positive for some $\tau \in [\tau_0, \tau_{\text{end}})$.

Since $\mathcal{H}_p^{\text{O,G}}(\mathbf{z}(\tau)) = \mathcal{H}_p^{\text{O,G}}(p(\tau), z(\tau))$ and from the chain rule, the persuasion condition is equivalent to

$$\frac{\mathrm{d}\mathcal{H}_p^{\text{O,G}}}{\mathrm{d}\tau} = \frac{\partial \mathcal{H}_p^{\text{O,G}}}{\partial p}\frac{\mathrm{d}p}{\mathrm{d}\tau} + \frac{\partial \mathcal{H}_p^{\text{O,G}}}{\partial z}\frac{\mathrm{d}z}{\mathrm{d}\tau} > 0 \tag{S4.3.3}$$

for all $\tau \in [\tau_1, \tau_2]$. Following [26], we say that the derivative

$$\frac{\partial \mathcal{H}_\xi^{\text{C,G}}}{\partial \zeta}$$

measures the evolutionary synergy of $\zeta$ on $\xi$: if the derivative is positive, there is evolutionary synergy; if it is negative, there is evolutionary interference. Motivated by (S4.3.3), we say that there is conflict dissolution via maternal reproductive specialization if there exist $\tau_0 < \tau_{\text{end}}$ such that (S4.3.1) hold and (S4.3.3) implies that

$$\frac{\partial \mathcal{H}_p^{\text{O,G}}}{\partial z}\frac{\mathrm{d}z}{\mathrm{d}\tau} > 0 \tag{S4.3.4}$$

for all $\tau \in [\tau_1, \tau_2]$. Thus, by material implication [i.e., $(A \implies B) \iff (\neg A \vee B)$], to establish that there is conflict dissolution via maternal reproductive specialization, it is sufficient that there is conflict dissolution ((S4.3.1) hold) and that (S4.3.4) holds for all $\tau \in [\tau_1, \tau_2]$. From (S4.3.3) and (S4.3.4), if reproductive effort increases over evolutionary time (i.e., $\mathrm{d}z/\mathrm{d}\tau > 0$), a necessary condition for conflict dissolution via maternal reproductive specialization is that there is evolutionary synergy of reproductive effort on helping, that is

$$\frac{\partial \mathcal{H}_p^{\text{O,G}}}{\partial z} > 0. \tag{S4.3.5}$$

Conflict dissolution can also be understood in terms of the benefit-cost ratio zones. If $(B/C)_p^{*\text{M,G}} < (B/C)_p^{*\text{O,G}}$ (condition (S4.1.3)) holds, conditions (S4.3.1) imply that conflict dissolution occurs if the system makes a transition from the conflict zone to the zone where both mother and offspring favor offspring helping, that is, if there are evolutionary times $\tau_0 < \tau_{\text{end}}$ such that

$$\left.\left(\frac{B}{C}\right)_p^{*\text{M,G}}\right|_{\mathbf{z}(\tau_0)} < \left.\frac{B}{C}\right|_{\mathbf{z}(\tau_0)} < \left.\left(\frac{B}{C}\right)_p^{*\text{O,G}}\right|_{\mathbf{z}(\tau_0)} \quad \text{and} \quad \left.\left(\frac{B}{C}\right)_p^{*\text{M,G}}\right|_{\mathbf{z}(\tau_{\text{end}})} < \left.\left(\frac{B}{C}\right)_p^{*\text{O,G}}\right|_{\mathbf{z}(\tau_{\text{end}})} < \left.\frac{B}{C}\right|_{\mathbf{z}(\tau_{\text{end}})} \tag{S4.3.6}$$

hold.

There are two basic pathways whereby conflict dissolution could happen in models related to ours. First, holding constant the benefit-cost ratio $B/C$, conflict dissolution requires that $(B/C)_p^{*\text{O,G}}$ decreases (equivalently, that its associated relative reproductive worth increases) over evolutionary time. This might occur, for instance, if brood sex proportions evolve in a model with a partially bivoltine life cycle (as in [27]). Second, holding constant the critical benefit-cost ratios $(B/C)_p^{*\text{M,G}}$ and $(B/C)_p^{*\text{O,G}}$ (e.g., if sex brood proportions are unbiased so (S4.1.2c) and hence (S4.1.7) hold), conflict dissolution requires the increase of the benefit-cost ratio $B/C$ over evolutionary time. In general, conflict dissolution might occur by a combination of the two pathways. We focus our analysis and results on the second pathway.

### 5 Evolutionary synergy and trade-off alleviation

We showed in the previous section that a necessary condition for conflict dissolution via maternal reproductive specialization is that there is evolutionary synergy of reproductive effort $z$ on helping $p$ (equation (S4.3.5)) as increasing $z$ evolves. In this section, we show that evolutionary synergy of reproductive effort $z$ on helping $p$ is equivalent to trade-off alleviation by helpers if reproductive effort is optimal. This yields the conclusion that, at an optimal reproductive effort, conflict dissolution via maternal reproductive specialization requires trade-off alleviation by helpers.

To do this, we proceed in four steps. First, we rewrite the selection gradient of reproductive effort in terms of elasticities, which quantify the assumed trade-offs (Selection gradient of reproductive effort in terms of elasticities; section 5.1). Second, we show that, if reproductive effort is optimal, evolutionary synergy of reproductive effort $z$ on helping $p$ is equivalent to a positive marginal effect of late fertility on the benefit of helping, $B$; we also show that, if reproductive effort is optimal, evolutionary synergy of helping $p$ on reproductive effort $z$ is equivalent to a positive marginal effect of helpers on the marginal productivity of late fertility, $D$ (Synergy of reproductive effort on helping and vice-versa; section 5.2). Third, we show that, if reproductive effort is optimal, such synergy is symmetric (evolutionary synergy of reproductive effort $z$ on helping $p$ is equivalent to evolutionary synergy of helping $p$ on reproductive effort $z$) and equivalent to late productivity being *supermodular* (Synergy as supermodularity of late productivity; section 5.3). Finally, we use these results to express the supermodularity of late productivity at an optimal reproductive effort in terms of trade-off alleviation by helpers (Synergy as trade-off alleviation; section 5.4).

#### 5.1 Selection gradient of reproductive effort in terms of elasticities

We begin by rewriting the selection gradient of reproductive effort in terms of elasticities, which offer a convenient way to quantify the trade-offs we have assumed. We have shown in section 2.4 that reproductive effort $z$ is under positive directional selection if the selection gradient of reproductive effort $\mathcal{S}_z(\mathbf{x})$ is positive, that is, if $\mathcal{S}_z(\mathbf{x}) > 0$ . We saw in section 2.9 that this condition is satisfied if and only if

$$D > 0, \tag{S5.1.1}$$

where

$$D = \frac{\partial \Pi_2}{\partial f_2}(f_2, h) \tag{S5.1.2}$$

is the marginal productivity of late fertility. Hence, holding the helping probability $p$ constant, selection leads to a (locally) optimal reproductive effort $z^*$, and corresponding (locally) optimal late fertility

$$f_2^* = f_2(z^*) \tag{S5.1.3}$$

that locally maximizes late productivity $\Pi_2$. Such an optimal $z^*$ satisfies the first-order condition

$$D|_{z^*} = D|_{f_2^*} = \frac{\partial \Pi_2}{\partial f_2}(f_2^*, h) = 0. \tag{S5.1.4}$$

Now, writing the late productivity $\Pi_2$ explicitly in terms of the vital rates (equation (S1.6.15)) and using the product rule of derivatives, we can rewrite equation (S5.1.2) as

$$\begin{aligned} D &= \frac{\partial}{\partial f_2}\left(s_M f_2 s_2\right) \\ &= \frac{\partial s_M}{\partial f_2} f_2 s_2 + s_M s_2 + s_M f_2 \frac{\partial s_2}{\partial f_2} \\ &= s_M s_2 \left(\frac{f_2}{s_M}\frac{\partial s_M}{\partial f_2} + 1 + \frac{f_2}{s_2}\frac{\partial s_2}{\partial f_2}\right) \\ &= s_M s_2 \left(\epsilon_{f_2}(s_M) + 1 + \epsilon_{f_2}(s_2)\right), \end{aligned} \tag{S5.1.5}$$

where we have identified

$$\epsilon_{f_2}(s_M) = \frac{f_2}{s_M}\frac{\partial s_M}{\partial f_2} = \frac{\partial \ln s_M}{\partial \ln f_2}, \text{ and} \tag{S5.1.6a}$$

$$\epsilon_{f_2}(s_2) = \frac{f_2}{s_2}\frac{\partial s_2}{\partial f_2} = \frac{\partial \ln s_2}{\partial \ln f_2}, \tag{S5.1.6b}$$

as, respectively, the (partial) elasticities of $s_M$ and $s_2$ with respect to $f_2$. The elasticity $\epsilon_X(Y)$ is the percent change in $Y$ caused by a marginal percent increase in $X$ holding all other variables constant. From our assumptions on the trade-offs between the vital rates (S1.4.6), at least one of the elasticities (S5.1.6) is negative but neither is positive. Thus, the elasticities (S5.1.6) quantify the trade-offs that we have assumed between vital rates.

From (S5.1.5) and since $s_M s_2 > 0$ (equation (S1.4.5)), a necessary and sufficient condition for $D > 0$ is that

$$\epsilon_{f_2}(s_M) + \epsilon_{f_2}(s_2) > -1. \tag{S5.1.7}$$

Together with (S5.1.4), this implies that the optimal reproductive effort $z^*$ is implicitly given by

$$\left(\epsilon_{f_2}(s_M) + \epsilon_{f_2}(s_2)\right)\Big|_{z=z^*} = -1. \tag{S5.1.8}$$

An elasticity equal to $-1$ means that a percent increase in the input variable leads to an exactly equal percent decrease in the output variable. Hence, condition (S5.1.7) states that a necessary and sufficient condition for reproductive effort to be favored to increase over evolutionary time is that a percent increase in late fertility $f_2$ caused by a marginal increase in reproductive effort leads to a weaker percent decrease in the total effect on maternal survival $s_M$ and second-brood survival $s_2$ (see also [28]).

#### 5.2 Synergy of reproductive effort on helping and vice-versa

We now show that, if reproductive effort is optimal, the evolutionary synergy of reproductive effort $z$ on helping $p$ can be equivalently expressed as either the marginal effect of $f_2$ on $B$ (section 5.2.1) or as the marginal effect of $h$ on $D$ (section 5.2.2).

##### 5.2.1 Synergy of reproductive effort on helping as late-fertility effects on benefit

At an optimal reproductive effort $z^*$, there is evolutionary synergy of reproductive effort $z$ on helping $p$ if

$$\left.\frac{\partial \mathcal{H}_p^{\mathrm{O,G}}}{\partial z}\right|_{z=z^*} > 0. \tag{S5.2.1}$$

Noting that the set of actors is the set of candidate helpers ($A = H$) when helping is under offspring control ($\mathrm{C} = \mathrm{O}$), taking the partial derivative, and by the product rule and the chain rule of derivatives, condition (S5.2.1) can be written as

$$\left(-\frac{\partial \omega_{H,H}}{\partial f_2}\frac{\mathrm{d}f_2}{\mathrm{d}z}C + \frac{\partial \omega_{H,P}}{\partial f_2}\frac{\mathrm{d}f_2}{\mathrm{d}z}B + \omega_{H,P}\frac{\partial B}{\partial f_2}\frac{\mathrm{d}f_2}{\mathrm{d}z}\right)_{z=z^*} > 0,$$

which, since $\mathrm{d}f_2/\mathrm{d}z > 0$, is equivalent to

$$\left(-\frac{\partial \omega_{H,H}}{\partial f_2}C + \frac{\partial \omega_{H,P}}{\partial f_2}B + \omega_{H,P}\frac{\partial B}{\partial f_2}\right)_{f_2=f_2^*} > 0. \tag{S5.2.2}$$

Now, for all $\mathrm{C} \in \{\mathrm{M},\mathrm{O}\}$ and all $\mathrm{G} \in \{\mathrm{B},\mathrm{F}\}$, reproductive worth $\omega_{A,R}$ depends on late fertility $f_2$ only through the reproductive value of females, $v_♀$. More specifically, the partial derivative of $\omega_{A,R}$ with respect to $f_2$ is proportional to the partial derivative of $v_♀$ with respect to $f_2$, which can be readily calculated as

$$\begin{aligned}
\frac{\partial v_♀}{\partial f_2} &= \frac{\partial}{\partial f_2}\left(\frac{\Pi_{♂,\mathrm{r},\mathrm{rr}}}{\Pi_{♀,\mathrm{r},\mathrm{rr}}}\right) \\
&= \frac{\left(\frac{\partial}{\partial f_2}\Pi_{♂,\mathrm{r},\mathrm{rr}}\right)\Pi_{♀,\mathrm{r},\mathrm{rr}} - \left(\frac{\partial}{\partial f_2}\Pi_{♀,\mathrm{r},\mathrm{rr}}\right)\Pi_{♂,\mathrm{r},\mathrm{rr}}}{\Pi^2_{♀,\mathrm{r},\mathrm{rr}}} \\
&= \frac{\left(\sigma_{2,♂}\frac{\partial}{\partial f_2}\Pi_{2,\mathrm{rr}}\right)\Pi_{♀,\mathrm{r},\mathrm{rr}} - \left(\sigma_{2,♀}\frac{\partial}{\partial f_2}\Pi_{2,\mathrm{rr}}\right)\Pi_{♂,\mathrm{r},\mathrm{rr}}}{\Pi^2_{♀,\mathrm{r},\mathrm{rr}}} \\
&= \frac{\left(\sigma_{2,♂}v_♂ - \sigma_{2,♀}v_♀\right)}{\Pi_{♀,\mathrm{r},\mathrm{rr}}}\frac{\partial \Pi_{2,\mathrm{rr}}}{\partial f_2} \\
&= \frac{\left(\sigma_{2,♂}v_♂ - \sigma_{2,♀}v_♀\right)}{\Pi_{♀,\mathrm{r},\mathrm{rr}}}\frac{\partial \Pi_2}{\partial f_2}(f_2,h) \\
&= \frac{\left(\sigma_{2,♂}v_♂ - \sigma_{2,♀}v_♀\right)}{\Pi_{♀,\mathrm{r},\mathrm{rr}}}D,
\end{aligned} \tag{S5.2.3}$$

where the first line follows from substituting equation (S2.6.12b), the second line applies the quotient rule of derivatives, the third line uses the derivatives of expression (S1.6.2) with respect to $f_2$, the fourth line uses the expressions for reproductive values (S2.6.12), the fifth line uses (S1.6.16), and the last line identifies the marginal productivity of late fertility $D$ (S2.9.3) and rearranges. Evaluating (S5.2.3) we then obtain, via (S5.1.4),

$$\left(\frac{\partial v_♀}{\partial f_2}\right)_{f_2=f_2^*} = \left.\frac{\left(\sigma_{2,♂}v_♂ - \sigma_{2,♀}v_♀\right)}{\Pi_{♀,\mathrm{r},\mathrm{rr}}}\right|_{f_2=f_2^*} \times D|_{f_2=f_2^*} = 0, \tag{S5.2.4}$$

so that the partial derivative of the reproductive value of females with respect to late fertility vanishes at an optimal late fertility. It follows that

$$\left(\frac{\partial \omega_{H,H}}{\partial f_2}\right)_{f_2=f_2^*} = \left(\frac{\partial \omega_{H,P}}{\partial f_2}\right)_{f_2=f_2^*} = 0,$$

and, since $\omega_{H,P} > 0$, condition (S5.2.2) simplifies to

$$\left(\frac{\partial B}{\partial f_2}\right)_{f_2=f_2^*} > 0.$$

Summarizing, we have

$$\left.\frac{\partial \mathcal{H}_p^{\mathrm{O,G}}}{\partial z}\right|_{z=z^*} > 0 \Longleftrightarrow \left(\frac{\partial B}{\partial f_2}\right)_{f_2=f_2^*} > 0, \tag{S5.2.5}$$

which states that, at an optimal reproductive effort, there is evolutionary synergy of reproductive effort $z$ on helping $p$ if and only if the marginal benefit of helping is increasing in late fertility, $f_2$.

##### 5.2.2 Synergy of helping on reproductive effort as helper effects on marginal productivity

Likewise, at an optimal reproductive effort $z^*$, there is evolutionary synergy of helping $p$ on reproductive effort $z$ if

$$\left.\frac{\partial \mathcal{H}_z^{\mathrm{C,G}}}{\partial p}\right|_{z=z^*} > 0. \tag{S5.2.6}$$

Taking the derivative of the inclusive fitness effect $\mathcal{H}_z^{\mathrm{C,G}}$ with respect to $p$, this condition can be written as

$$\left(\frac{\partial \omega_{M,O_2}}{\partial p} D + \omega_{M,O_2} \frac{\partial D}{\partial h}\frac{\partial h}{\partial p}\right)_{z=z^*} > 0, \tag{S5.2.7}$$

where $\omega_{M,O_2}$ is the reproductive worth for a mother of a second-brood offspring. Since $D$ vanishes at $z = z^*$, and since $\omega_{M,O_2} > 0$ and $\partial h/\partial p = \bar{h} > 0$, this condition simplifies to

$$\left(\frac{\partial D}{\partial h}\right)_{f_2=f_2^*} > 0.$$

Summarizing, we have that

$$\left.\frac{\partial \mathcal{H}_z^{\mathrm{C,G}}}{\partial p}\right|_{z=z^*} > 0 \iff \left(\frac{\partial D}{\partial h}\right)_{f_2=f_2^*} > 0, \tag{S5.2.8}$$

which states that, at an optimal reproductive effort, there is evolutionary synergy of helping $p$ on reproductive effort $z$ if and only if the marginal productivity of late fertility is increasing in the expected number of helpers, $h$.

#### 5.3 Synergy as supermodularity of late productivity

We now show that, at an optimal reproductive effort, the conditions for evolutionary synergy of helping on reproductive effort (S5.2.1) and for evolutionary synergy of reproductive effort on helping (S5.2.6) are equivalent, and that both are equivalent to the condition that late productivity is supermodular.

This observation is immediate from the fact that the right-hand inequalities in (S5.2.5) and (S5.2.8) are equivalent. Indeed, it follows both from our definitions of marginal benefit of helping $B$ (S2.8.5) and marginal productivity of late fertility $D$ (S2.9.3), and from the symmetry of second derivatives, that

$$\frac{\partial B}{\partial f_2} = \frac{\partial^2 \Pi_2}{\partial f_2 \partial h} = \frac{\partial^2 \Pi_2}{\partial h \partial f_2} = \frac{\partial D}{\partial h}, \tag{S5.3.1}$$

and hence that

$$\frac{\partial^2 \Pi_2}{\partial f_2 \partial h} > 0 \iff \frac{\partial B}{\partial f_2} > 0 \iff \frac{\partial D}{\partial h} > 0. \tag{S5.3.2}$$

Since this identity also holds at an optimal level of late fertility $f_2^*$, we have

$$\frac{\partial^2 \Pi_2}{\partial f_2 \partial h}(f_2^*, h) > 0 \iff \left(\frac{\partial B}{\partial f_2}\right)_{f_2=f_2^*} > 0 \iff \left(\frac{\partial D}{\partial h}\right)_{f_2=f_2^*} > 0. \tag{S5.3.3}$$

Expression (S5.3.1) reminds us of the connection between the partial derivatives of the marginal productivity of one input (e.g., expected number of helpers, $h$) with respect to the other (e.g., the late fertility $f_2$). Expression (S5.3.2) reminds us of the fact that the condition for the marginal productivity of one variable to be increasing in the other is equal to the condition that the cross partial derivatives of the late productivity function $\Pi_2(f_2, h)$ are positive, that is, that the late productivity $\Pi_2$ is supermodular. Supermodularity formalizes a classic way of interpreting the notion of complementarity in economics; namely that having more of one input increases the marginal returns to having more of another input [29]. In our case, supermodularity of $\Pi_2$ means that having more helpers increases the marginal productivity of late fertility, and that having more late fertility (via increased reproductive effort) increases the marginal productivity of helping, that is, that helping and reproductive effort act as *strategic complements*.

In conclusion, we have, via expressions (S5.3.3), (S5.2.5) and (S5.2.8), that

$$\frac{\partial^2 \Pi_2}{\partial f_2 \partial h}(f_2^*, h) > 0 \Longleftrightarrow \left.\frac{\partial \mathcal{H}_p^{\mathrm{O,G}}}{\partial z}\right|_{z=z^*} > 0 \Longleftrightarrow \left.\frac{\partial \mathcal{H}_z^{\mathrm{C,G}}}{\partial p}\right|_{z=z^*} > 0. \tag{S5.3.4}$$

Expression (S5.3.4) states that the supermodularity of the late productivity $\Pi_2$ (i.e., the complementarity between helping and reproductive effort) at an optimal reproductive effort is a necessary and sufficient condition for evolutionary synergy between helping and reproductive effort. Such evolutionary synergy means that helping and reproductive effort are in positive feedback whereby the evolution of reproductive effort increases selection for helping, and the evolution of helping increases selection for reproductive effort.

#### 5.4 Synergy as trade-off alleviation

**Trade-off alleviation.** The condition on the supermodularity of the late productivity function $\Pi_2$ appearing on the left hand side of (S5.3.4) can be given a demographically meaningful interpretation in terms of the way helping by offspring alleviates life-history trade-offs faced by mothers. To do so, note that we can write the cross partial derivative as

$$\begin{aligned}
\frac{\partial^2 \Pi_2}{\partial f_2 \partial h}(f_2^*, h) &= \left(\frac{\partial D}{\partial h}\right)_{f_2=f_2^*} \\
&= \left\{\frac{\partial}{\partial h}\left[s_M s_2\left(\epsilon_{f_2}(s_M) + 1 + \epsilon_{f_2}(s_2)\right)\right]\right\}_{f_2=f_2^*} \\
&= \left[\frac{\partial (s_M s_2)}{\partial h}\left(\epsilon_{f_2}(s_M) + 1 + \epsilon_{f_2}(s_2)\right) + s_M s_2 \frac{\partial\left(\epsilon_{f_2}(s_M) + 1 + \epsilon_{f_2}(s_2)\right)}{\partial h}\right]_{f_2=f_2^*},
\end{aligned} \tag{S5.4.1}$$

where we made use of (S5.3.1) in the first line, of (S5.1.5) in the second line, and of the product rule of derivatives in the third line.

Since at an optimal reproductive effort, $\epsilon_{f_2}(s_M) + 1 + \epsilon_{f_2}(s_2) = 0$ holds (see (S5.1.8)), equation (S5.4.1) simplifies to

$$\frac{\partial^2 \Pi_2}{\partial f_2 \partial h}(f_2^*, h) = \left[s_M s_2 \frac{\partial\left(\epsilon_{f_2}(s_M) + 1 + \epsilon_{f_2}(s_2)\right)}{\partial h}\right]_{f_2=f_2^*}. \tag{S5.4.2}$$

Given that $s_M s_2 > 0$ (see (S1.4.5)), it follows that

$$\frac{\partial^2 \Pi_2}{\partial f_2 \partial h}(f_2^*, h) > 0 \Longleftrightarrow \left.\left(\frac{\partial \epsilon_{f_2}(s_M)}{\partial h} + \frac{\partial \epsilon_{f_2}(s_2)}{\partial h}\right)\right|_{f_2=f_2^*} > 0. \tag{S5.4.3}$$

As previously stated, $\epsilon_{f_2}(s_M)$ and $\epsilon_{f_2}(s_2)$ measure the percent life-history trade-offs faced by a mother by increasing her late fertility $f_2$. Hence, condition (S5.4.3) states that, at an optimal reproductive effort, the condition for $\Pi_2$ to be supermodular is equivalent to the condition that helpers alleviate the proportional life-history trade-offs. Therefore, together with (S4.3.3) and (S5.3.4), condition (S5.4.3) yields the conclusion that conflict dissolution via maternal reproductive specialization requires that helpers alleviate trade-offs as optimal reproductive effort evolves.

**Comparative statics of optimal reproductive effort with respect to the expected number of helpers.** A consequence of the supermodularity of the late productivity function is that a given (locally) optimal reproductive effort $z^*$ is increasing in the expected number of helpers (see, e.g., [29]). That is,

$$\frac{\partial^2 \Pi_2}{\partial f_2 \partial h}(f_2^*, h) > 0 \Longleftrightarrow \frac{\partial z^*}{\partial h} > 0. \tag{S5.4.4}$$

For our purposes, this can be proven using the implicit function theorem as follows. A locally optimal late fertility value $f_2^*$ is implicitly given by (see equation (S5.1.4))

$$\frac{\partial \Pi_2}{\partial f_2}(f_2^*, h) = 0. \tag{S5.4.5}$$

Differentiating with respect to $h$, we have

$$\frac{\partial^2 \Pi_2}{\partial f_2^2}(f_2^*, h)\frac{\partial f_2^*}{\partial h} + \frac{\partial^2 \Pi_2}{\partial h \partial f_2}(f_2^*, h) = 0, \tag{S5.4.6}$$

so that solving for $\partial f_2^* / \partial h$ we get

$$\frac{\partial f_2^*}{\partial h} = -\frac{\dfrac{\partial^2 \Pi_2}{\partial h \partial f_2}(f_2^*, h)}{\dfrac{\partial^2 \Pi_2}{\partial f_2^2}(f_2^*, h)} > 0, \tag{S5.4.7}$$

from which (S5.4.4) follows by the chain rule, because $\dfrac{\partial^2 \Pi_2}{\partial f_2^2}(f_2^*, h) < 0$ holds (as $z^*$ is a local maximum) and $f_2(z)$ is an increasing function.

**Examples of late productivity functions that do not allow for evolutionary synergy.** There are at least two important classes of possible late productivity functions that do not allow for evolutionary synergy: additively separable functions, and multiplicatively separable functions.

First, consider late productivity functions that are additively separable, that is, late productivity functions that could be written as

$$\Pi_2(f_2, h) = \Pi_{2,1}(f_2) + \Pi_{2,2}(h) \tag{S5.4.8}$$

with $\Pi_{2,1} : \mathbb{R}_+^* \to \mathbb{R}_+^*$ and $\Pi_{2,2} : [0, f_1] \to \mathbb{R}_+^*$. A function of the form of (S5.4.8) is not supermodular in any point of its domain, as the cross partial derivative is zero at all points. It then follows that the condition in the left hand side of (S5.3.4) is never satisfied.

Second, consider late productivity functions that are multiplicatively separable, that is, one could find functions $\Pi_{2,1} : \mathbb{R}_+^* \to \mathbb{R}_+^*$ and $\Pi_{2,2} : [0, f_1] \to \mathbb{R}_+^*$ so that

$$\Pi_2(f_2, h) = \Pi_{2,1}(f_2) \times \Pi_{2,2}(h) \tag{S5.4.9}$$

holds. To show that for functions of the form (S5.4.9) there is no evolutionary synergy between helping and fertility at an optimal late fertility level, note first that in the case of a multiplicatively separable $\Pi_2$ function, the first order condition for an optimal reproductive effort (S5.1.4) implies

$$\frac{\mathrm{d}\Pi_{2,1}}{\mathrm{d}f_2}(f_2^*) = 0. \tag{S5.4.10}$$

Note second that evaluating the cross partial derivative of $\Pi_2$ at an optimal fertility level, we obtain

$$\begin{aligned}
\left.\frac{\partial^2\Pi_2}{\partial f_2 \partial h}\right|_{f_2=f_2^*} &= \frac{\partial}{\partial h}\left.\left(\frac{\partial \Pi_2}{\partial f_2}\right)\right|_{f_2=f_2^*} \\
&= \frac{\partial}{\partial h}\left.\left(\Pi_{2,2}(h)\frac{\mathrm{d}\Pi_{2,1}}{\mathrm{d}f_2}\right)\right|_{f_2=f_2^*} \\
&= \left.\left(\frac{\mathrm{d}\Pi_{2,2}}{\mathrm{d}h}\frac{\mathrm{d}\Pi_{2,1}}{\mathrm{d}f_2} + \Pi_{2,2}(h)\frac{\partial}{\partial h}\left(\frac{\mathrm{d}\Pi_{2,1}}{\mathrm{d}f_2}\right)\right)\right|_{f_2=f_2^*} \\
&= \frac{\mathrm{d}\Pi_{2,2}}{\mathrm{d}h}(h)\frac{\mathrm{d}\Pi_{2,1}}{\mathrm{d}f_2}(f_2^*) \\
&= 0
\end{aligned}$$

where the third line applies the product rule of derivatives, the fourth line follows because $\mathrm{d}\Pi_{2,1}/\mathrm{d}f_2$ is independent of $h$ (and hence $\partial(\mathrm{d}\Pi_{22}/\mathrm{d}f_2)/\partial h = 0$), and the last line follows from (S5.4.10).

#### 5.5 Summary of equivalences

To summarize, the following statements are equivalent.

1. Increasing reproductive effort increases selection for helping at optimal reproductive effort:

$$\left.\frac{\partial \mathcal{H}_p^{\mathrm{O,G}}}{\partial z}\right|_{z=z^*} > 0.$$

2. Increasing late fertility increases the benefit of helping at optimal late fertility:

$$\left(\frac{\partial B}{\partial f_2}\right)_{f_2=f_2^*} > 0.$$

3. Increasing helping increases selection for reproductive effort at optimal reproductive effort:

$$\left.\frac{\partial \mathcal{H}_z^{\mathrm{C,G}}}{\partial p}\right|_{z=z^*} > 0.$$

4. Increasing helpers increases the marginal productivity of late fertility at optimal late fertility:

$$\left(\frac{\partial D}{\partial h}\right)_{f_2=f_2^*} > 0.$$

5. Late fertility is supermodular at optimal late fertility:

$$\frac{\partial^2\Pi_2}{\partial f_2 \partial h}(f_2^*, h) > 0.$$

6. Helpers alleviate the total percent trade-off at optimal late fertility:

$$\left.\left(\frac{\partial \epsilon_{f_2}(s_M)}{\partial h} + \frac{\partial \epsilon_{f_2}(s_2)}{\partial h}\right)\right|_{f_2=f_2^*} > 0.$$

7. Helpers increase the optimal reproductive effort:

$$\frac{\partial z^*}{\partial h} > 0.$$

8. Helpers increase the optimal late fertility:

$$\frac{\partial f_2^*}{\partial h} > 0.$$

### 6 Evolutionary dynamics

In this section, we write equations describing the evolutionary dynamics of the evolving traits. To do this, we proceed in two steps. First, in section 6.1 (Canonical equation) we write the evolutionary dynamic equations by postulating that our evolving traits satisfy a form of the canonical equation of adaptive dynamics [2, 30, 31]. Second, in section 6.2 (Resulting evolutionary dynamic equations when traits are genetically uncorrelated) we write the evolutionary dynamic equations that result when traits are genetically uncorrelated.

#### 6.1 Canonical equation

We follow the evolutionary dynamics of the phenotypic vector $\mathbf{z}$. Given our assumptions of $\delta$-weak selection and rare mutation, we expect that, in our model, invasion implies fixation [4] and that the deterministic evolutionary dynamics are to first order approximately given by a form of the canonical equation of adaptive dynamics [2, 30, 31]. Thus, we conjecture that the evolutionary dynamics of $\mathbf{z}$ over evolutionary time $\tau$ are to first order given by

$$\frac{\mathrm{d}\mathbf{z}}{\mathrm{d}\tau} = \mathbf{G}(\mathbf{z})\boldsymbol{\mathcal{S}}(\mathbf{z}), \tag{S6.1.1}$$

with a covariance matrix $\mathbf{G}(\mathbf{z})$ given by

$$\mathbf{G}(\mathbf{z}) = \begin{pmatrix} \mathcal{G}_{pp} & \mathcal{G}_{pz} \\ \mathcal{G}_{zp} & \mathcal{G}_{zz} \end{pmatrix}$$

for model cases of offspring or maternal control, and by

$$\mathbf{G}(\mathbf{z}) = \begin{pmatrix} \mathcal{G}_{xx} & \mathcal{G}_{xy} & \mathcal{G}_{xz} \\ \mathcal{G}_{yx} & \mathcal{G}_{yy} & \mathcal{G}_{yz} \\ \mathcal{G}_{zx} & \mathcal{G}_{zy} & \mathcal{G}_{zz} \end{pmatrix}$$

for model cases of shared control. The $\zeta\xi$-th entry $\mathcal{G}_{\zeta\xi}(\mathbf{z})$ of $\mathbf{G}$ is proportional to the covariance of mutational effects $\mathrm{Cov}[Z_\mathrm{m}-\zeta, \Xi_\mathrm{m}-\xi] = \mathrm{Cov}[Z_\mathrm{m}, \Xi_\mathrm{m}]$, where $Z_\mathrm{m}$ and $\Xi_\mathrm{m}$ are random variables with small variation around their respective expected values $\mathrm{E}[Z_\mathrm{m}] = \zeta$ and $\mathrm{E}[\Xi_\mathrm{m}] = \xi$. The diagonal entries $\mathcal{G}_{\zeta\zeta}(\mathbf{z})$ are non-negative, and we also denote them as $\mathcal{G}_\zeta(\mathbf{z})$. $\mathbf{G}$ is symmetric. If traits are genetically uncorrelated, then $\mathbf{G}$ is diagonal.

#### 6.2 Resulting evolutionary dynamic equations when traits are genetically uncorrelated

When traits are genetically uncorrelated, the resulting evolutionary dynamic equations are

$$\frac{\mathrm{d}\zeta}{\mathrm{d}\tau} = \mathcal{G}_\zeta \frac{\bar{h}}{\mathbf{v}^\intercal\mathbf{u}} \omega_{A,H}\left(-c_\zeta + \rho_{A,H,P} b_\zeta\right) \tag{S6.2.1a}$$

$$\frac{\mathrm{d}z}{\mathrm{d}\tau} = \mathcal{G}_z \frac{1}{\mathbf{v}^\intercal\mathbf{u}} \omega_{M,O_2} b_z, \tag{S6.2.1b}$$

for $\zeta$ affecting helping (i.e., $\zeta \in \{p\}$ for model cases of offspring or maternal control and $\zeta \in \{x, y\}$ for model cases of shared control; using equations (S6.1.1), (S3.5.1), (S3.5.2), (S3.3.1), (S3.6.2), and (S3.6.3)). We now list the resulting dynamic equations for each model case.

**Offspring control, both sexes help.** When helping is under offspring control and both sexes help, and from equations (S6.2.1), (S1.1.5), (S3.4.3), (S3.6.1), (S3.1.1), and (S3.1.2), the evolutionary dynamics equations are

$$\frac{\mathrm{d}p}{\mathrm{d}\tau} = \mathcal{G}_p \frac{f_1}{\mathbf{v}^\mathsf{T}\mathbf{u}} \omega_{O_1,O_1} \left(-C + \rho_{O_1,O_1,O_2} B^{\mathrm{O,B}}\right), \tag{S6.2.2a}$$

$$\frac{\mathrm{d}z}{\mathrm{d}\tau} = \mathcal{G}_z \frac{1}{\mathbf{v}^\mathsf{T}\mathbf{u}} \omega_{M,O_2} \frac{\mathrm{d}f_2}{\mathrm{d}z} D^{\mathrm{O,B}}. \tag{S6.2.2b}$$

For the particular case of unbiased sex proportions in both broods (i.e., $\sigma_{1,♀} = \sigma_{1,♂} = \sigma_{2,♀} = \sigma_{2,♂} = 1/2$) using Fig. S7 and (S3.3.4), equations (S6.2.2) further simplify for both diploids and haplodiploids to

$$\frac{\mathrm{d}p}{\mathrm{d}\tau} = \mathcal{G}_p \frac{f_1}{\mathbf{v}^\mathsf{T}\mathbf{u}} \frac{1}{2} \left(\sum_{\ell\in\{♀,♂\}} u_\ell v_\ell\right) \left(-C + \frac{1}{2} B^{\mathrm{O,B}}\right), \tag{S6.2.3a}$$

$$\frac{\mathrm{d}z}{\mathrm{d}\tau} = \mathcal{G}_z \frac{1}{\mathbf{v}^\mathsf{T}\mathbf{u}} u_♀ \frac{1}{2} \left(\sum_{\ell\in\{♀,♂\}} r_{\bullet(M),\circ(O_{2\ell})} v_\ell\right) \frac{\mathrm{d}f_2}{\mathrm{d}z} D^{\mathrm{O,B}}. \tag{S6.2.3b}$$

For diploids, each of the sums over $\ell$ in parentheses in equations (S6.2.3) equals 1.

**Offspring control, only females help.** When helping is under offspring control and only females help, and from equations (S6.2.1), (S1.1.5), (S3.4.3), (S3.6.1), (S3.1.1), and (S3.1.2), the evolutionary dynamics equations are

$$\frac{\mathrm{d}p}{\mathrm{d}\tau} = \mathcal{G}_p \frac{f_1 \sigma_{1,♀}}{\mathbf{v}^\mathsf{T}\mathbf{u}} \omega_{O_{1♀},O_{1♀}} \left(-C + \rho_{O_{1♀},O_{1♀},O_2} B^{\mathrm{O,F}}\right), \tag{S6.2.4a}$$

$$\frac{\mathrm{d}z}{\mathrm{d}\tau} = \mathcal{G}_z \frac{1}{\mathbf{v}^\mathsf{T}\mathbf{u}} \omega_{M,O_2} \frac{\mathrm{d}f_2}{\mathrm{d}z} D^{\mathrm{O,F}}. \tag{S6.2.4b}$$

**Maternal control, both sexes help.** When helping is under maternal control and both sexes help, and from equations (S6.2.1), (S1.1.5), (S3.4.3), (S3.6.1), (S3.1.1), and (S3.1.2), the evolutionary dynamics equations are

$$\frac{\mathrm{d}p}{\mathrm{d}\tau} = \mathcal{G}_p \frac{f_1}{\mathbf{v}^\mathsf{T}\mathbf{u}} \omega_{M,O_1} \left(-C + \rho_{M,O_1,O_2} B^{\mathrm{M,B}}\right), \tag{S6.2.5a}$$

$$\frac{\mathrm{d}z}{\mathrm{d}\tau} = \mathcal{G}_z \frac{1}{\mathbf{v}^\mathsf{T}\mathbf{u}} \omega_{M,O_2} \frac{\mathrm{d}f_2}{\mathrm{d}z} D^{\mathrm{M,B}}. \tag{S6.2.5b}$$

For the particular case of unbiased sex proportions in both broods (i.e., $\sigma_{1,♀} = \sigma_{1,♂} = \sigma_{2,♀} = \sigma_{2,♂} = 1/2$) using Fig. S7 and (S3.3.7), equations (S6.2.5) further simplify for both diploids and haplodiploids to

$$\frac{\mathrm{d}p}{\mathrm{d}\tau} = \mathcal{G}_p \frac{f_1}{\mathbf{v}^\mathsf{T}\mathbf{u}} u_♀ \frac{1}{2} \left(\sum_{\ell\in\{♀,♂\}} r_{\bullet(M),\circ(O_{1\ell})} v_\ell\right) \left(-C + B^{\mathrm{M,B}}\right) \tag{S6.2.6a}$$

$$\frac{\mathrm{d}z}{\mathrm{d}\tau} = \mathcal{G}_z \frac{1}{\mathbf{v}^\mathsf{T}\mathbf{u}} u_♀ \frac{1}{2} \left(\sum_{\ell\in\{♀,♂\}} r_{\bullet(M),\circ(O_{2\ell})} v_\ell\right) \frac{\mathrm{d}f_2}{\mathrm{d}z} D^{\mathrm{M,B}}. \tag{S6.2.6b}$$

For diploids, each of the sums over $\ell$ in parentheses in equations (S6.2.6) equals 1.

**Maternal control, only females help.** When helping is under maternal control and only females help, and from equations (S6.2.1), (S1.1.5), (S3.4.3), (S3.6.1), (S3.1.1), and (S3.1.2), the evolutionary dynamics equations are

$$\frac{\mathrm{d}p}{\mathrm{d}\tau} = \mathcal{G}_p \frac{f_1 \sigma_{1,♀}}{\mathbf{v}^\mathsf{T}\mathbf{u}} \omega_{M,O_{1♀}} \left(-C + \rho_{M,O_{1♀},O_2} B^{\mathrm{M,F}}\right) \tag{S6.2.7a}$$

$$\frac{\mathrm{d}z}{\mathrm{d}\tau} = \mathcal{G}_z \frac{1}{\mathbf{v}^\mathsf{T}\mathbf{u}} \omega_{M,O_2} \frac{\mathrm{d}f_2}{\mathrm{d}z} D^{\mathrm{M,F}}. \tag{S6.2.7b}$$

**Shared control, both sexes help.** When helping is under shared control and both sexes help, and from equations (S6.2.1), (S1.1.5), (S3.4.3), (S3.6.1), (S3.1.1), and (S3.1.2), the evolutionary dynamics equations are

$$\frac{\mathrm{d}x}{\mathrm{d}\tau} = \mathcal{G}_x \frac{f_1}{\mathbf{v}^\intercal \mathbf{u}} \omega_{M,O_1} \frac{\partial p}{\partial x}(x,y)\left(-C + \rho_{M,O_1,O_2} B^{\mathrm{S,B}}\right) \tag{S6.2.8a}$$

$$\frac{\mathrm{d}y}{\mathrm{d}\tau} = \mathcal{G}_y \frac{f_1}{\mathbf{v}^\intercal \mathbf{u}} \omega_{O_1,O_1} \frac{\partial p}{\partial y}(x,y)\left(-C + \rho_{O_1,O_1,O_2} B^{\mathrm{S,B}}\right) \tag{S6.2.8b}$$

$$\frac{\mathrm{d}z}{\mathrm{d}\tau} = \mathcal{G}_z \frac{1}{\mathbf{v}^\intercal \mathbf{u}} \omega_{M,O_2} \frac{\mathrm{d}f_2}{\mathrm{d}z} D^{\mathrm{S,B}}. \tag{S6.2.8c}$$

For the particular case of unbiased sex proportions in both broods (i.e., $\sigma_{1,♀} = \sigma_{1,♂} = \sigma_{2,♀} = \sigma_{2,♂} = 1/2$) using Fig. S7, (S3.3.7), and (S3.3.4), equations (S6.2.8) further simplify for both diploids and haplodiploids to

$$\frac{\mathrm{d}x}{\mathrm{d}\tau} = \mathcal{G}_x \frac{f_1}{\mathbf{v}^\intercal \mathbf{u}} u_{♀} \frac{1}{2}\left(\sum_{\ell \in \{♀,♂\}} r_{\bullet(M),\circ(O_{1\ell})} v_\ell\right) \frac{\partial p}{\partial x}(x,y)\left(-C + B^{\mathrm{S,B}}\right) \tag{S6.2.9a}$$

$$\frac{\mathrm{d}y}{\mathrm{d}\tau} = \mathcal{G}_y \frac{f_1}{\mathbf{v}^\intercal \mathbf{u}} \frac{1}{2}\left(\sum_{\ell \in \{♀,♂\}} u_\ell v_\ell\right) \frac{\partial p}{\partial y}(x,y)\left(-C + \frac{1}{2} B^{\mathrm{S,B}}\right) \tag{S6.2.9b}$$

$$\frac{\mathrm{d}z}{\mathrm{d}\tau} = \mathcal{G}_z \frac{1}{\mathbf{v}^\intercal \mathbf{u}} u_{♀} \frac{1}{2}\left(\sum_{\ell \in \{♀,♂\}} r_{\bullet(M),\circ(O_{2\ell})} v_\ell\right) \frac{\mathrm{d}f_2}{\mathrm{d}z} D^{\mathrm{S,B}}. \tag{S6.2.9c}$$

For diploids, each of the sums over $\ell$ in parentheses in equations (S6.2.9) equals 1.

**Shared control, only females help.** When helping is under shared control and only females help, and from equations (S6.2.1), (S1.1.5), (S3.4.3), (S3.6.1), (S3.1.1), and (S3.1.2), the evolutionary dynamics equations are

$$\frac{\mathrm{d}x}{\mathrm{d}\tau} = \mathcal{G}_x \frac{f_1 \sigma_{1,♀}}{\mathbf{v}^\intercal \mathbf{u}} \omega_{M,O_{1♀}} \frac{\partial p}{\partial x}(x,y)\left(-C + \rho_{M,O_{1♀},O_2} B^{\mathrm{S,F}}\right) \tag{S6.2.10a}$$

$$\frac{\mathrm{d}y}{\mathrm{d}\tau} = \mathcal{G}_y \frac{f_1 \sigma_{1,♀}}{\mathbf{v}^\intercal \mathbf{u}} \omega_{O_{1♀},O_{1♀}} \frac{\partial p}{\partial y}(x,y)\left(-C + \rho_{O_{1♀},O_{1♀},O_2} B^{\mathrm{S,F}}\right) \tag{S6.2.10b}$$

$$\frac{\mathrm{d}z}{\mathrm{d}\tau} = \mathcal{G}_z \frac{1}{\mathbf{v}^\intercal \mathbf{u}} \omega_{M,O_2} \frac{\mathrm{d}f_2}{\mathrm{d}z} D^{\mathrm{S,F}}. \tag{S6.2.10c}$$

### 7 Specific functional forms

In this section, we specify the functional forms for the vital rates composing late productivity $\Pi_2(h,z)$ (Vital rates composing late productivity; section 7.1) and for the joint phenotype $p(x,y)$ for helping under shared control (Joint helping phenotype; section 7.2) that we use to illustrate our results in the main text.

#### 7.1 Vital rates composing late productivity

We consider the following effects of helping and of reproductive effort. We let helpers increase only the couple survival $s_M(f_2,h)$. In turn, reproductive effort increases the late fertility $f_2(z)$ and decreases only the couple survival $s_M(f_2,h)$. We let second-brood survival $s_2(f_2,h)$ be constant. Specifically, we use the following functional forms for the vital rates composing late productivity:

$$f_2(z) = f_0 z^{\alpha}, \tag{S7.1.1a}$$

$$s_M(f_2,h) = \overline{s_M}(h)\left(1 - \frac{f_2}{\overline{f_2}(h)}\right), \tag{S7.1.1b}$$

$$s_2(f_2,h) = s_2, \tag{S7.1.1c}$$

where $s_2$ denotes a real-valued constant in the interval $(0,1]$, $\overline{s_M}(h)$ and $\overline{f_2}(h)$ are positive increasing functions of $h$, with $\overline{s_M}(\bar{h}) \le 1$, and where the domain $S = S_M \times [0,\bar{h}]$ of $s_M$ (see (S1.4.5a)) is given by

$$S = \left\{(f_2,h) \in \mathbb{R}_+^* \times [0,\bar{h}] : f_2 < \overline{f_2}(h)\right\},$$

so that the image of $s_M$ is the interval $(0,1)$. Thus, for a given $h$, $s_M(f_2,h)$ is a linear function of $f_2$ with negative slope equal to $-\overline{s_M}(h)/\overline{f_2}(h)$ and intercept equal to $\overline{s_M}(h)$. It follows that, for a given $h$, $\overline{s_M}(h)$ is the maximum couple survival that can be achieved (as late fertility $f_2 \to 0$) and $\overline{f_2}(h)$ is the maximum late fertility that can be achieved with a positive couple survival (as $s_M \to 0$). Eq. (S7.1.1b) thus specifies the simplest kind of trade-off between $s_M(f_2,h)$ and $f_2$: a linear trade-off.

Late productivity is given by the product of the three vital rates, hence

$$\Pi_2(f_2,h) = s_M(f_2,h) f_2 s_2. \tag{S7.1.2}$$

The benefit of helping is then

$$B = \frac{\partial \Pi_2}{\partial h} = \left[\frac{\mathrm{d}\overline{s_M}(h)}{\mathrm{d}h}\left(1 - \frac{f_2}{\overline{f_2}(h)}\right) + \overline{s_M}(h)\frac{f_2}{\overline{f_2}^2(h)}\frac{\mathrm{d}\overline{f_2}(h)}{\mathrm{d}h}\right] f_2 s_2, \tag{S7.1.3}$$

which is positive since $\overline{s_M}(h)$ and $\overline{f_2}(h)$ are increasing in $h$.

The marginal productivity of late fertility is given by

$$D = \frac{\partial \Pi_2}{\partial f_2} = s_2 \overline{s_M}(h)\left(1 - \frac{2 f_2}{\overline{f_2}(h)}\right). \tag{S7.1.4}$$

$D$ has a single sign change from positive to negative as $f_2$ increases. This happens at the optimal late fertility rate

$$f_2^*(h) = \frac{\overline{f_2}(h)}{2}, \tag{S7.1.5}$$

obtained at an optimal level of reproductive effort equal to

$$z^*(h) = \left(\frac{\overline{f_2}(h)}{2f_0}\right)^{1/\alpha}. \tag{S7.1.6}$$

Hence, for each value of $h$, the optimal late fertility is half the maximum late fertility. As $\overline{f_2}(h)$ is increasing in $h$, so is $f_2^*(h)$. This is to be expected as there is synergy of optimal reproductive effort on helping, since

$$\left.\frac{\partial \Pi_2}{\partial h \partial f_2}\right|_{f_2=f_2^*} = s_2 \overline{s_M}(h) \frac{1}{\overline{f_2}(h)} \frac{\mathrm{d}\overline{f_2}(h)}{\mathrm{d}h} > 0$$

holds. Further note that $s_M^*(h) = s_M(f_2^*, h) = \overline{s_M}(h)/2$.

We have assumed that $\overline{f_2}(h)$ is strictly increasing. Suppose for a moment that $\overline{f_2}(h) = \bar{f}_2$, where $\bar{f}_2$ is a constant. This is an example where the resulting late productivity function $\Pi_2$ is multiplicatively separable (cf. equation (S5.4.9)). Hence, $f_2^* = \bar{f}_2$ is independent of $h$ and there is not synergy of optimal reproductive effort on helping, as

$$\left.\frac{\partial \Pi_2}{\partial h \partial f_2}\right|_{f_2=f_2^*} = 0.$$

To complete the specification of the vital rates composing late productivity, we use the functions

$$\overline{s_M}(h) = \underline{s_M} + \left(\overline{\overline{s_M}} - \underline{s_M}\right)\frac{h}{\bar{h}}, \tag{S7.1.7a}$$

$$\overline{f_2}(h) = \underline{f_2} + \left(\overline{\overline{f_2}} - \underline{f_2}\right)\frac{h}{\bar{h}}, \tag{S7.1.7b}$$

where the constant $\underline{s_M} \in (0,1]$ gives the smallest possible intercept for couple survival attained at $h = 0$, the constant $\overline{\overline{s_M}} \in [\underline{s_M}, 1]$ gives the largest possible intercept for couple survival attained at $h = \bar{h}$, the constant $\underline{f_2} \in \mathbb{R}_+^*$ gives the smallest possible value of $\overline{f_2}(h)$ attained at $h = 0$, and the constant $\overline{\overline{f_2}} \in [\underline{f_2}, \infty)$ gives the largest possible value of $\overline{f_2}(h)$ attained at $h = \bar{h}$ (the resulting $s_M(f_2, h)$ with the parameter values used is plotted in Fig. 3).

#### 7.2 Joint helping phenotype

Here we specify the function for the joint helping phenotype $p(x, y)$ for model cases of shared control. We suppose that maternal influence $x$ and offspring resistance $y$ engage in a contest to achieve the expression of the helping phenotype $p$. We consider two different kinds of contests. First, we consider simultaneous contests, where maternal influence $x$ and offspring resistance $y$ contest simultaneously to determine the helping probability. For this kind of contest, we assume that the helping probability is given by the probability that the mother wins an imperfectly discriminating contest within the class of contest success functions proposed and axiomatized by [32]. Specifically, we assume

$$p(x, y) = \frac{g_\mathrm{M}(x;\chi)}{1 + g_\mathrm{M}(x;\chi) + g_\mathrm{O}(y;\psi)}, \tag{S7.2.1}$$

where $g_\mathrm{M}(x;\chi)$ and $g_\mathrm{O}(y;\psi)$ are “impact functions” to be specified below, with parameters $\chi > 0$ and $\psi > 0$ measuring the “power” of mother and offspring, respectively.

Second, we also consider sequential contests, where the mother acts first (engaging in a contest "against nature"; e.g., secreting molecules that alter offspring development) and the offspring acts second (e.g., by subsequently readjusting its own development). For these contests we assume the following general form:

$$\begin{aligned} p(x,y) &= \frac{g_\mathrm{M}(x;\chi)}{1+g_\mathrm{M}(x;\chi)}\left(1-\frac{g_\mathrm{O}(y;\psi)}{1+g_\mathrm{O}(y;\psi)}\right) \\ &= \frac{g_\mathrm{M}(x;\chi)}{1+g_\mathrm{M}(x;\chi)+g_\mathrm{O}(y;\psi)+g_\mathrm{M}(x;\chi)g_\mathrm{O}(y;\psi)} \end{aligned} \tag{S7.2.2}$$

for impact functions $g_\mathrm{M}(x;\chi)$ and $g_\mathrm{O}(y;\psi)$.

We assume that the impact functions $g_\mathrm{M}(x;\chi)$ and $g_\mathrm{O}(y;\psi)$ satisfy the following properties:

1. $g_\mathrm{M}(x;\chi)$ and $g_\mathrm{O}(y;\psi)$ are non-negative strictly increasing functions $g_i : \mathbb{R}_+ \to \mathbb{R}_+$, $i \in \{\mathrm{M},\mathrm{O}\}$. This can be interpreted as the impact functions measuring the absolute effort devoted to the contest.

2. $g_\mathrm{M}(x;\chi)$ and $g_\mathrm{O}(y;\psi)$ are strictly increasing in their parameters, that is $\partial g_\mathrm{M}(x;\chi)/\partial\chi > 0$ and $\partial g_\mathrm{M}(x;\psi)/\partial\psi > 0$. This can be interpreted as power increasing the ability of the effort to succeed in the contest.

3. $g_\mathrm{M}(0;\chi) = 0$. This can be interpreted as stating that without maternal influence, the mother devotes no effort to contest offspring helping.

It follows that $p(x,y)$ satisfies:

1. $p(x,y) \in [0,1]$ for all $x \geq 0$, $y \geq 0$ (i.e., the helping probability is well defined).

2. $p(x,y)$ is strictly increasing in $x$ and strictly decreasing in $y$ (i.e., maternal influence and offspring resistance affect the helping probability as required by (S1.2.3)).

3. $p(0,y) = 0$ (i.e., there is no helping in the absence of maternal influence).

4. For given $x \geq 0$ and $y \geq 0$, $p(x,y)$ is strictly increasing in $\chi$ and strictly decreasing in $\psi$ (i.e., the "power" of maternal influence can be increased by increasing $\chi$ and the "power" of offspring resistance can be increased by increasing $\psi$).

It remains to specify the impact function. We consider an exponential function of the kind

$$g_\mathrm{M}(x;\chi) = \mathrm{e}^{\chi x} - 1, \tag{S7.2.3a}$$

$$g_\mathrm{O}(y;\psi) = \mathrm{e}^{\psi y} - 1, \tag{S7.2.3b}$$

which satisfies the required properties and has been used in contest models [33] (we add the $-1$ in the exponential impact function to satisfy $g_\mathrm{M}(0;\chi) = 0$). The resulting joint phenotype is illustrated in Fig. S11.

##### 7.3 Comparison between simultaneous and sequential contests

From equations (S2.8.41), we have that the relative evolutionary speed but not the direction of selection of maternal influence and offspring resistance depend respectively on $\partial p/\partial x$ and $\partial p/\partial y$. Thus, conflict dissolution is promoted by greater $\partial p/\partial x$ and smaller $|\partial p/\partial y|$. We now determine whether conflict dissolution is promoted by simultaneous (S7.2.1) or sequential (S7.2.1) contests.

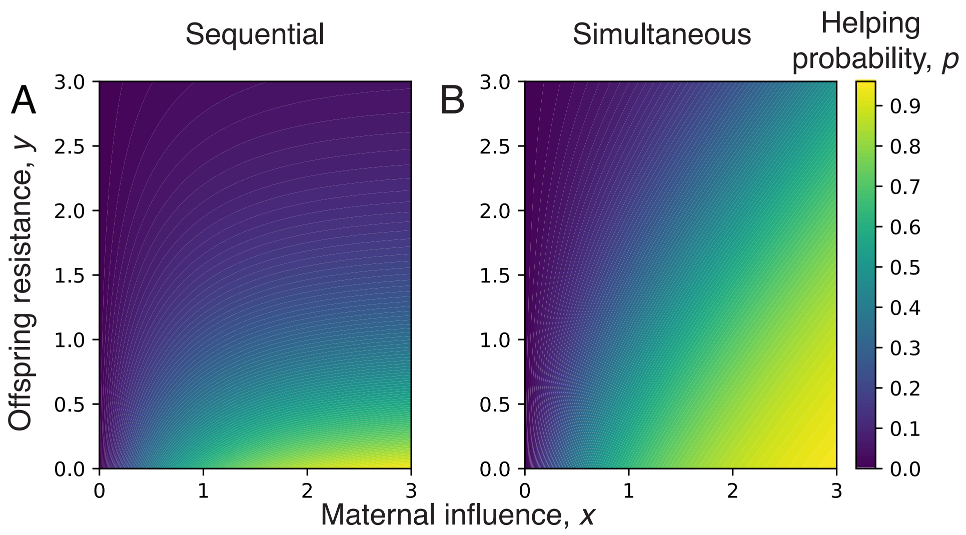

Figure S11: Joint helping phenotype. The helping probability $p(x,y)$ under (A) sequential or (B) simultaneous contests. Parameter values are as in Fig. 2; in particular, mother and offspring have the same power in both panels ($\chi = \psi = 1$).

Denoting by $p_{\text{sim}}(x,y)$ the helping probability defined by (S7.2.1), we obtain

$$\frac{\partial p_{\text{sim}}}{\partial x} = \frac{1+g_{\text{O}}}{(1+g_{\text{M}}+g_{\text{O}})^2}\frac{\partial g_{\text{M}}}{\partial x}.$$

Similarly, denoting by $p_{\text{seq}}(x,y)$ the helping probability defined by (S7.2.2), we obtain

$$\frac{\partial p_{\text{seq}}}{\partial x} = \frac{1}{(1+g_{\text{M}})^2(1+g_{\text{O}})}\frac{\partial g_{\text{M}}}{\partial x}.$$

Thus,

$$\begin{aligned}\frac{\partial p_{\text{sim}}}{\partial x} - \frac{\partial p_{\text{seq}}}{\partial x} &= \frac{1+g_{\text{O}}}{(1+g_{\text{M}}+g_{\text{O}})^2}\frac{\partial g_{\text{M}}}{\partial x} - \frac{1}{(1+g_{\text{M}})^2(1+g_{\text{O}})}\frac{\partial g_{\text{M}}}{\partial x} \\ &= \left[\frac{(1+g_{\text{O}})^2}{(1+g_{\text{M}}+g_{\text{O}})^2} - \frac{1}{(1+g_{\text{M}})^2}\right]\frac{1}{1+g_{\text{O}}}\frac{\partial g_{\text{M}}}{\partial x},\end{aligned}$$

which is positive if

$$\frac{(1+g_{\text{O}})^2}{(1+g_{\text{M}}+g_{\text{O}})^2} - \frac{1}{(1+g_{\text{M}})^2} > 0.$$

This condition holds whenever $g_{\text{O}} > 0$. Therefore, the evolutionary speed of maternal influence is promoted by simultaneous relative to sequential contests, provided that $g_{\text{O}} > 0$, which holds for an exponential impact function (S7.2.3b) if there is some resistance.

Proceeding analogously for resistance, we have that

$$\begin{aligned}\frac{\partial p_{\text{sim}}}{\partial y} &= -\frac{g_{\text{M}}}{(1+g_{\text{M}}+g_{\text{O}})^2}\frac{\partial g_{\text{O}}}{\partial y} \\ \frac{\partial p_{\text{seq}}}{\partial y} &= -\frac{g_{\text{M}}}{(1+g_{\text{M}})(1+g_{\text{O}})^2}\frac{\partial g_{\text{O}}}{\partial y}.\end{aligned}$$

Thus,

$$\begin{aligned}\left|\frac{\partial p_{\text{seq}}}{\partial y}\right| - \left|\frac{\partial p_{\text{sim}}}{\partial y}\right| &= \frac{g_{\text{M}}}{(1+g_{\text{M}})(1+g_{\text{O}})^2}\frac{\partial g_{\text{O}}}{\partial y} - \frac{g_{\text{M}}}{(1+g_{\text{M}}+g_{\text{O}})^2}\frac{\partial g_{\text{O}}}{\partial y} \\ &= \left[\frac{1}{(1+g_{\text{M}})(1+g_{\text{O}})^2} - \frac{1}{(1+g_{\text{M}}+g_{\text{O}})^2}\right] g_{\text{M}}\frac{\partial g_{\text{O}}}{\partial y},\end{aligned}$$

which, provided that $g_{\mathrm{M}} > 0$, is positive if

$$\frac{1}{(1+g_{\mathrm{M}})(1+g_{\mathrm{O}})^2} - \frac{1}{(1+g_{\mathrm{M}}+g_{\mathrm{O}})^2} > 0.$$

This condition reduces to

$$1 + g_{\mathrm{M}} - g_O^2 > 0,$$

which is positive for sufficiently small $g_{\mathrm{O}}$. Therefore, the evolutionary speed of offspring resistance is hindered by simultaneous relative to sequential contests, provided that there is some maternal influence and that resistance is small.

We conclude that conflict dissolution is promoted by simultaneous relative to sequential contests, provided that there is some maternal influence, some offspring resistance, and that resistance is small.

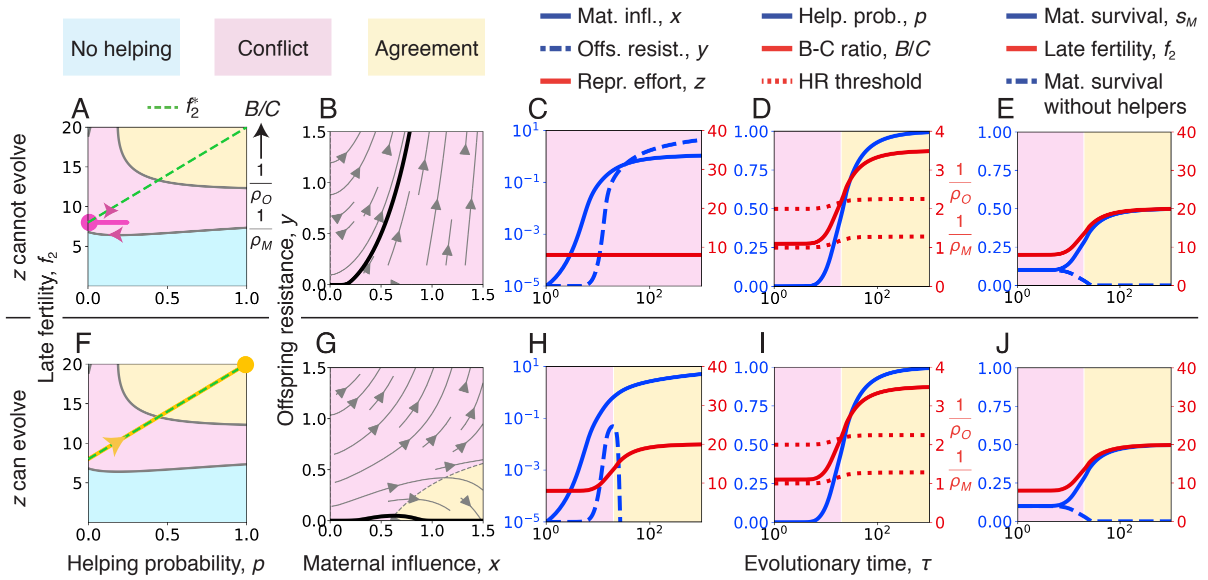

Figure S12: Conflict dissolution via maternal reproductive specialization (evolutionary model) in haplodiploids. Analogous plots to Fig. 2. Same parameter values except that here the genetic system is haplodiploid, only females help, $\underline{f_2} = 16$, and $\overline{\overline{f_2}} = 40$.

### 8 Specification of Fig. 2, and additional figures

The specification of Fig. 2 is the following. The genetic system is diploid, both sexes help, and the determination of the joint helping phenotype is sequential. Functions:

$$f_2(z) = f_0 z^{\alpha}, \tag{S8.0.1a}$$

$$s_M(f_2, h) = \left(\underline{s_M} + \left(\overline{\overline{s_M}} - \underline{s_M}\right)\frac{h}{\bar{h}}\right)\left(1 - \frac{f_2}{\underline{f_2} + \left(\overline{\overline{f_2}} - \underline{f_2}\right)\frac{h}{\bar{h}}}\right), \tag{S8.0.1b}$$

$$p(x, y) = \mathrm{e}^{-\chi x - \psi y}\left(\mathrm{e}^{\chi x} - 1\right), \tag{S8.0.1c}$$

$$\mathcal{G}_\zeta = G_\zeta\left(1 - \mathrm{e}^{-\beta\zeta}\right) \text{ for } \zeta \in \{x, y\}. \tag{S8.0.1d}$$

Parameter values: $f_0 = 1$, $\alpha = 1$, $\underline{s_M} = 0.2$, $\overline{\overline{s_M}} = 1$, $f_1 = 8$, $\underline{f_2} = 36$, $\overline{\overline{f_2}} = 72$, $s_1 = s_2 = 0.1$, $\chi = \psi = 1$, $\sigma_{1♀} = \sigma_{2♀} = 0.5$, $G_x = G_y = 1$, and $\beta = 100$. Traits are genetically uncorrelated: $\mathcal{G}_{xy} = \mathcal{G}_{xz} = \mathcal{G}_{yz} = 0$. Initial conditions for $\mathbf{z}(\tau) = (x(\tau), y(\tau), z(\tau))^\intercal$ are $x(0) = y(0) = 10^{-5}$ and $z(0) = z^*(0)$. For Fig. 2A-E, $z$ is constant. For Fig. 2F-J, $z$ is equal to $z^*(h)$.

Conflict dissolution in haplodiploids is shown in Fig. S12. Promoters of conflict dissolution in haplodiploids are described in Fig. S13. Conflict dissolution with low genetic variance of reproductive effort is shown in Fig. S14. In all cases, $\mathcal{G}_z$ follows the functional form given in eq. (S8.0.1d) with $\beta = 100$.

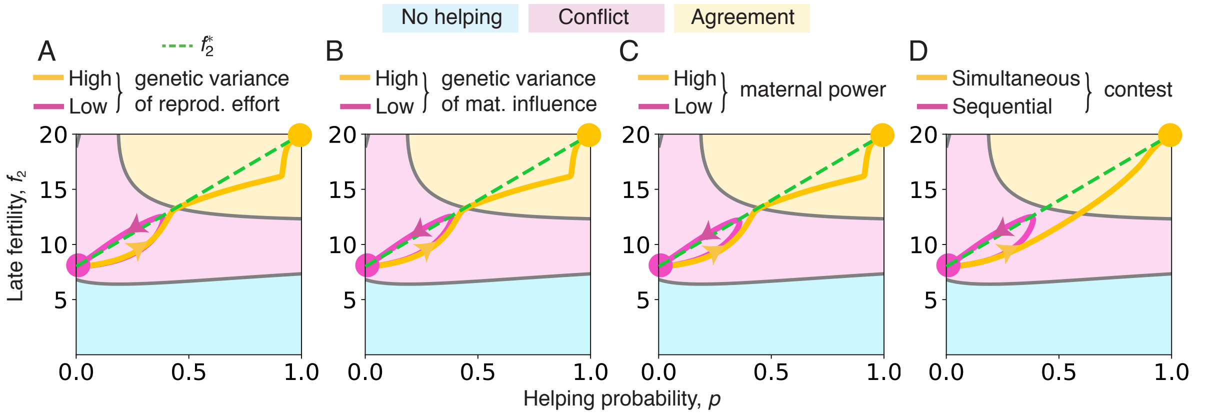

Figure S13: Promoters of conflict dissolution in haplodiploids. Analogous plots to Fig. S12 except that here the genetic system is haplodiploid, only females help, and parameter values are as in Fig. S12 with the following genetic variances. For A, $G_z = 70$ for low genetic variance of $z$ and $G_z = 80$ for high genetic variance of $z$. For B, $G_x = 0.9$ for low genetic variance of $x$ and $G_x = 1$ for high genetic variance of $x$ (and $G_z = 80$ for both). For C, $\chi = 0.9$ for low maternal power and $\chi = 1$ for high maternal power (and $G_z = 80$ for both). For D, sequential contest and simultaneous contest (and $G_z = 70$ for both).

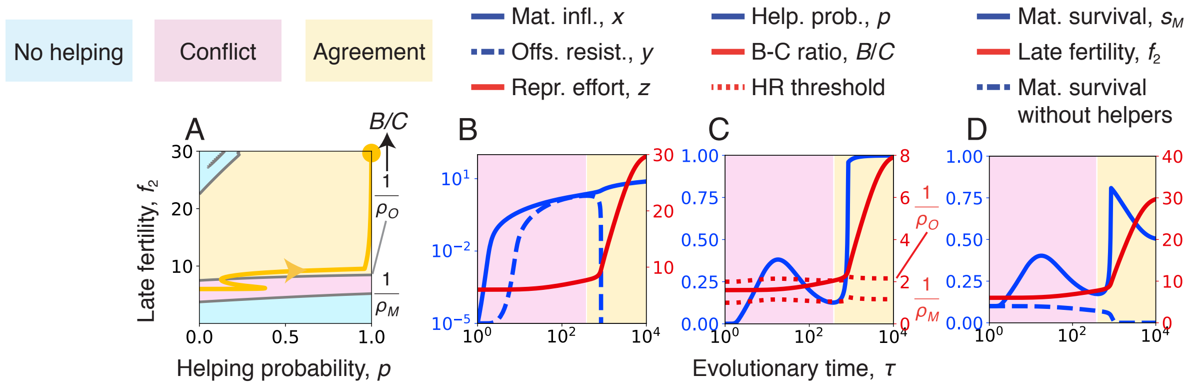

Figure S14: Conflict dissolution with low genetic variance of reproductive effort. The genetic system is haplodiploid and only females help. Analogous plots to Fig. S12F,H,I,J. Same parameter values except that here $f_1 = 6$, $\underline{f_2} = 12$, $\overline{\overline{f_2}} = 60$, and $G_x = G_y = G_z = 1$.
